# Purifying selection purges harmful variants in the rarest pine

**DOI:** 10.1101/2024.08.07.607108

**Authors:** Ren-Gang Zhang, Wei Zhao, Hui Liu, Ruo-Fei Li, Heng Shu, De-Tuan Liu, Zheng-Jiang Liu, Richard Milne, Jun Chen, Xiong-Fang Liu, Hong-Yun Shang, Min-Jie Zhou, Yi-Qing Wang, Hao Yang, Kai-Hua Jia, Xiao-Quan Wang, Hang Sun, Wei-Bang Sun, Loren Riesberg, Yong-Peng Ma

## Abstract

Population genetic theory predicts that severe bottlenecks and extremely small effective population sizes (*N*_e_) should reduce the ability of natural selection to eliminate harmful mutations. Under this framework, deleterious alleles are expected to accumulate and even fix, eroding fitness, constraining evolutionary rescue, and potentially precipitating mutational meltdown. Yet, empirical tests of these predictions in species at the extreme lower bound of *N*_e_ remain rare. We address this gap using *Pinus squamata*, one of the rarest tree species on Earth, with only 35 wild individuals remaining. We generated a near-complete reference genome (29.2 Gb) for this species and performed population genomic analyses across nearly all of its extant individuals, together with two closely related species. *P. squamata* exhibits extraordinarily low nucleotide diversity (π = 3.35 × 10⁻⁵), the lowest reported for any plant. Demographic inference reveals a recent and severe bottleneck (∼20 generations ago) that reduced *N*_e_ to ∼2.7 and resulted in intense inbreeding. Contrary to theoretical expectations, we uncover evidence for highly efficient purifying selection: strongly deleterious mutations are markedly depleted, indicating substantial purging despite the extremely small *N*_e_. Genome-wide patterns further implicate selection at linked sites—including background selection and pseudo-overdominance—as dominant forces shaping genomic variation in the species. These results challenge the prevailing view that drift overwhelms selection in extremely small populations. Instead, they suggest that, under certain genomic and demographic conditions, purifying selection can remain unexpectedly effective, potentially mitigating the risk of mutational meltdown in the rarest species.

## Introduction

The theoretical foundation of conservation genetics centers on small populations, as most taxa of conservation concern currently persist as narrow remnants, and common management actions often begin with only a handful of individuals (Ellstrand and Elam, 1993; Frankham et al., 2010; Ma et al., 2013; Sun et al., 2024). Under the (nearly) neutral theory, small effective population size (*N*_e_) is expected to intensify genetic drift and reduce the efficiency of natural selection (Kimura, 1968; Ohta, 1992), a pattern documented in multiple species with small populations (Bemmels et al., 2025; Leroy et al., 2021). Under such conditions, deleterious alleles are expected to accumulate and potentially become fixed rather than being purged (Lynch et al., 1995). This can erode fitness, constrain evolutionary rescue, and precipitate mutational meltdowns.

Conversely, elevated inbreeding in small populations can expose recessive deleterious alleles to stronger purifying selection, potentially enabling their removal, but at the cost of short-term fitness (Dussex et al., 2023; Glémin, 2003; Hedrick and Garcia-Dorado, 2016). Empirical evidence for such purging is rapidly accumulating (Chen et al., 2025; Garcia-Erill et al., 2025; Grossen et al., 2020; Khan et al., 2021; Kleinman-Ruiz et al., 2022; Yang et al., 2025). As inbreeding enhances purifying selection while drift erodes it, however, their net outcome when *N*_e_ approaches single digits remains unclear. Whether purifying selection can remain efficient enough to counter mutational meltdown in extremely small, highly inbred populations is therefore an open question—one with direct implications for conservation.

*Pinus squamata* X.W. Li (Qiaojia pine), first described in 1992 (Li, 1992), is classified as Critically Endangered on the IUCN Red List and is designated a Plant Species with Extremely Small Populations (PSESP) in China, receiving Grade-I national protection. Field surveys recored just 35 wild trees, all on Yaoshan Mountain, Qiaojia County, Yunnan (Tao et al., 2022). Such a small census population size (*N*_c_) makes it the rarest pine on record. Since 1995, wild seeds have been collected and propagated; today ca. 2,000 and 600 cultivated adults exist for *in situ* (population augmentation) and *ex situ* conservation, respectively, along with thousands of seedlings (Shu, 2024). Despite these conservation efforts, genetic management has been hindered by the species’ very large genome size (∼30 Gb). Nevertheless, *P. squamata* provides an unparalleled model for testing population genetic theory when population size is exceptionally small.

Here, we present a near-complete genome assembly for *P. squamata* and present population genomic data for nearly all extant wild individuals, plus two closely related congeners, *P. bungeana* and *P. gerardiana*. Population genomics analyses reveal a severe, recent bottleneck that has driven genome-wide diversity to record lows and reduced *N*_e_ to single digits. Contrary to the expectation that drift should dominate its evolutionary process under such extremes, we detect an exceptionally strong signature of purifying selection: a markedly higher fraction of strongly deleterious mutations has been purged from the genome relative to closely related species. Our results provide the first molecular evidence that, even when *N*_e_ approaches single digits, purifying selection—not drift—can govern the genomic landscape, challenging a central expectation of conservation genetics theory, and offering new perspectives on the evolution and conservation of plant species with extremely small populations.

## Results

### Genome characteristics of *P. squamata*

To investigate how extreme population contraction has shaped the genome of this critically endangered conifer *Pinus squamata*, we first assembled a chromosome-level reference genome from an individual propagated *ex situ* from wild-collected seeds (GL03) (**Supplementary Figure 1**). *K*-mer analysis (*k* = 19) estimated a genome size of ∼30 Gb with exceptionally low heterozygosity (∼0.001%) (**Supplementary Figure 2**). We generated 431 Gb (14.4×) of PacBio HiFi reads, 1.41 Tb (47.1×) of Oxford Nanopore reads, 4.57 Tb (152.3×) of Hi-C data, and 1.1 Tb (36.7×) of Illumina short reads for assembling the giant genome (**Supplementary Table 1**). We constructed contigs from HiFi and ONT reads and scaffolded them with Hi-C, assembling 12 pseudochromosomes, consistent with the haploid chromosome number in *Pinus* (2*n* = 24; **Fig. 1A-B**, **Supplementary Figures. 3–4**). The final assembly was 29.2 Gb, with a contig N50 and a scaffold N50 of 915.4 Mb and 2.5 Gb, respectively, and only 41 gaps were left (**Supplementary Table 2**). *K*-mer analysis confirmed 99.7% completeness (QV = 51.8), and BUSCO assessment showed 91.3% completeness (**Supplementary Table 2**). We further identified 20 telomeric and 12 centromeric regions (**Supplementary Figure 5**). Together, this assembly represents the most complete conifer genome to date, surpassing previously reported gymnosperm assemblies in both continuity and accuracy (**Supplementary Figure 4**).

**Figure 1.**
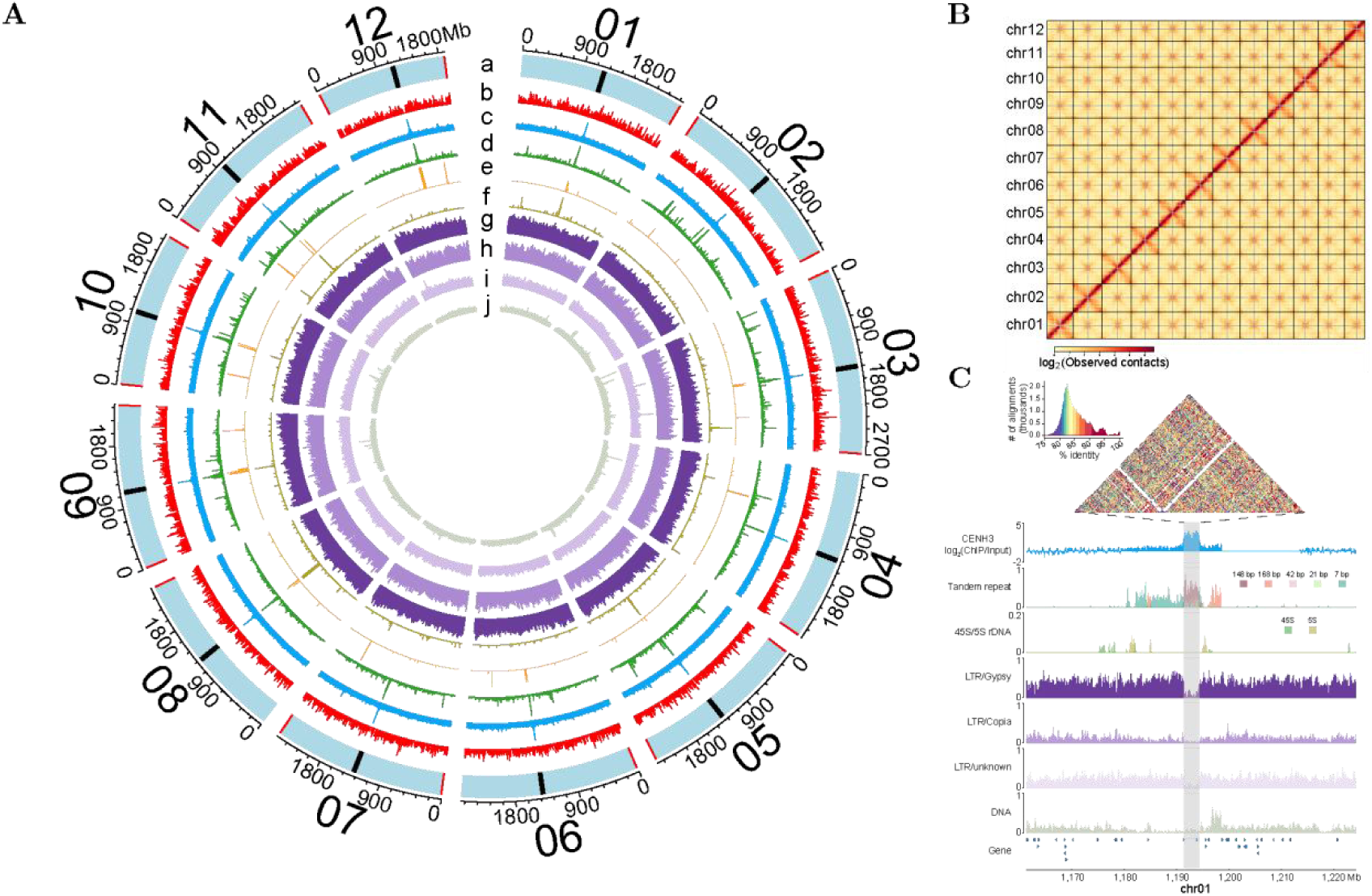
Genome features and centromere characteristics of *P. squamata*. **A**) Circos plot of genomic features: (a) centromere (black) and telomere (red) positions, (b) gene density, (c) CENH3 enrichment [log_2_(ChIP/input)], (d) tandem repeat, (e) 45S rDNA, (f) 5S rDNA, (g) LTR/Gypsy, (h) LTR/Copia, (i) LTR/unknown, (j) DNA transposons. **B**) Hi-C heatmap of the genome assembly. Dashed gray lines across the inter-chromosomal intense interaction signals indicate the positions of centromeres. **C**) Characteristics of a typical centromere from chromosome 1.

Annotation identified 55,413 protein-coding genes (PCGs) based on *ab initio* prediction, transcriptome evidence, and homology support (**Supplementary Table 3**). BUSCO assessment indicated 94.9% completeness of the gene set, and 88.0% of PCGs were functionally annotated using public protein databases (**Supplementary Table 4**). We also identified 24,280 rRNA, 20,398 tRNA, and 4,209 other non-coding RNA genes. Transposable elements (TEs) accounted for 75.1% of the genome, with long terminal repeat retrotransposons (LTR-RTs) being the most abundant, accounting for 59.5% of the genome (**Supplementary Table 5**).

Hi-C contact maps revealed a canonical Rabl configuration across all chromosomes (**Fig. 1B**), characterized by strong intrachromosomal arm interactions and enriched inter-centromeric contacts, indicating spatial clustering of centromeres within the nucleus (Marie-Nelly et al., 2014). Using these interaction patterns, we identified centromere regions and further refined their boundaries based on enrichment of 148-bp tandem repeats (**Supplementary Figures 6–8; Supplementary Table 6**). CENH3 ChIP–seq independently validated centromeric locations, but revealed centromere repositioning to an adjacent site on chromosome 2 (chr02) and chr10 (**Fig. 1C; Supplementary Figure 8**). Sequence similarity analyses showed that centromeric repeats were highly conserved across chromosomes (**Supplementary Figure 6**). The peri-centromeric regions were enriched with telomere-like 7-bp motifs (TTTAAGG), 45S and 5S rRNA loci, and LTR/Gypsy elements (**Supplementary Figures. 8–11**), consistent with cytological observations in *Pinus* (Hizume et al., 2002; Liu et al., 2003; Murray, 2013). The GC content of peri-centromeric rDNA was significantly lower than the genome-wide average (Wilcoxon test, *P* < 0.001; **Supplementary Figure 12**), suggesting ongoing degeneration and transcriptional inactivation, consistent with long-term heterochromatinization and relaxed selective constraints. Notably, this is the first report of centromere sequence characteristics in gymnosperms.

### Genomic consequences of extreme population contraction in *P. squamata*

To characterize population genomic parameters of *P. squamata*, we resequenced nearly all known wild individuals of P. squamata (*n* = 33) (**Supplementary Table 7**), with the exception of two trees located precariously on steep cliffs, which were inaccessible due to safety concerns. For comparison, we included 10 individuals of *P. bungeana*, a relatively widespread conifer native to northern China, and 9 individuals of *P. gerardiana*, a Himalayan species with a widespread but fragmented distribution. These two species are the closest relatives of *P. squamata* (Jin et al., 2021). We performed genome-wide comparative analyses, and found high genetic differentiation and deep divergence among the three species (mean *F*_ST_ = 0.90 and mean *d*_XY_ = 0.015) (**Supplementary Note 1**, **Supplementary Figure 13**), although they form a monophyletic group (Jin et al., 2021).

*P. squamata* displays exceptionally low nucleotide diversity (mean *π* = 3.35 × 10⁻⁵; median *π* = 8.45 × 10⁻^6^, across 1Mb windows), which is over two orders of magnitude lower than that of *P. gerardiana* (mean *π* = 3.07 × 10⁻^3^; median *π* = 2.63 × 10⁻^3^) and *P. bungeana* (mean *π* = 1.70 × 10⁻^3^; median *π* = 1.21 × 10⁻^3^) (**Fig. 2A**). Intriguingly, nucleotide diversity of *P. squamata* is also substantially below the range reported for other sequenced angiosperms, gymnosperms, ferns (**Fig. 2A, Supplementary Table 8**), and animals (Chen et al., 2017; Corbett-Detig et al., 2015; Garcia-Erill et al., 2025; Leffler et al., 2012). Furthermore, *π* in over half of the genome (55.5% of 1-Mb windows) falls below 1 × 10⁻⁵, and it drops below 1 × 10⁻⁶ in 4.8% of the genome (**Fig. 2B**). As a consequence of the extremely low population-level *π*, *P. squamata* also shows markedly reduced individual-level heterozygosity, with an average of 2.24 × 10⁻⁵ heterozygotes per bp (range: 0.95 × 10⁻⁵ – 4.22 × 10⁻⁵; **Fig. 2C**). In contrast, *P. gerardiana* and *P. bungeana* exhibit substantially higher heterozygosity (3.73 × 10⁻³ and 2.23 × 10⁻³, respectively; Wilcoxon test, *P* < 1 × 10⁻⁴). These results indicate that *P. squamata* harbors the lowest genetic diversity observed in plants to date, providing an invaluable natural system to examine the genomic consequences of extreme genetic erosion.

**Figure 2.**
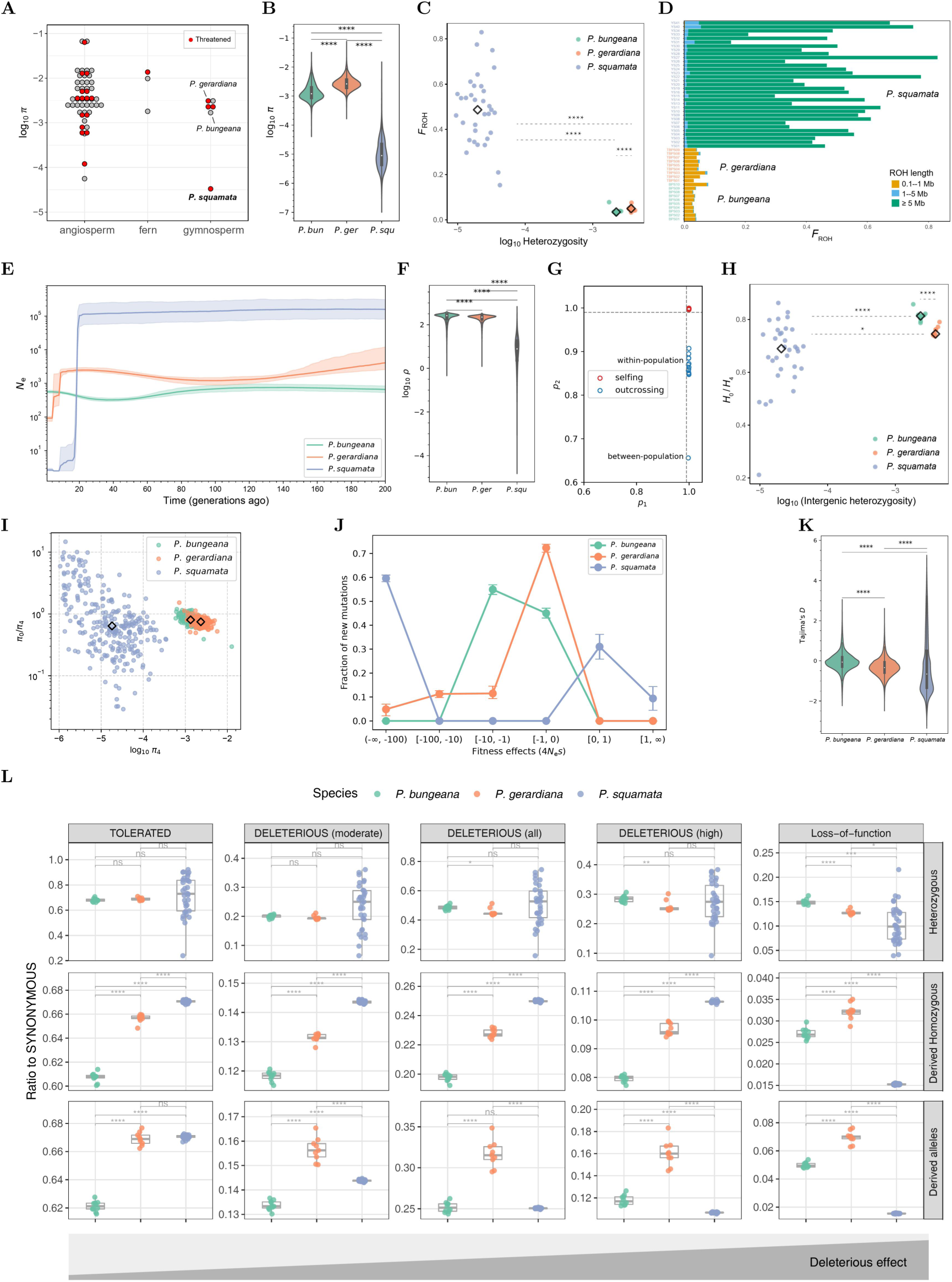
Population comparison of *P. squamata* with two closely related pines. **A**) Nucleotide diversity (*π*) in the three pines and sequenced plants. **B**) genome-wide *π* across 1-Mb windows in the three pines. **C**) Individual heterozygosity and *F*_ROH_ in the three species. **D**) *F*_ROH_ partitioned by ROH length. **E**) Demographic history estimated by GONE over the past 200 generations. **F**) Genome-wide population recombination rate (*ρ*). **G**) Estimation of selfing rate. *p*_1_, the proportion of progeny in which one allele matching the parental alleles; *p*_2_, the proportion of progeny in which both alleles match the parental alleles. **H**) Ratio of the heterozygosity at zero-fold to four-fold degenerate sites (*H*_0_/*H*_4_) and putatively neutral heterozygosity in each individual. *H*_0_/*H*_4_ is an individual-level proxy of purifying selection efficacy. **I**) *π*₀/*π*₄ across 100Mb windows. *π*₀/*π*₄ is a population-level proxy for purifying selection strength; lower values indicate stronger selection suppressing diversity at zero-fold degenerate sites. **J**) Distribution of fitness effects (DFE) modeled on site frequency spectra of four-fold (putatively neutral) and zero-fold (putatively selected) degenerate sites. 4*N*_e_*s* is population selection coefficient. **K**) Genome-wide Tajima’s *D* across 1-Mb windows. **L**) Ratio of non-synonymous alleles across different deleterious effects. Ratios of heterozygous genotypes, derived homozygous genotypes and total derived alleles reflect masked load, realized load, and total genetic load, respectively. Statistical test: *, *P* < 0.05; ****, *P* < 1e-4, Wilcoxon rank-sum test. Diamond symbols indicate median values in panels C, H and I.

To characterize the history of inbreeding, we identified runs of homozygosity (ROH)—contiguous genomic regions that are identical by descent. Long ROHs (>5 Mb) typically reflect recent inbreeding, while shorter ROHs may result from older inbreeding or demographic contractions (Ceballos et al., 2018). In *P. squamata,* ROH regions covered half of the genome (mean *F*_ROH_ = 0.49), far exceeding the values in *P. gerardiana* (mean *F*_ROH_= 0.050) and *P. bungeana* (0.038) (**Fig. 2C**). Notably, most ROH segments in *P. squamata* exceeded 5 Mb suggesting widespread and recent inbreeding (**Fig. 2D**).

Collectively, the extensive ROH, reduced heterozygosity, and extremely low nucleotide diversity point to severe genetic erosion in *P. squamata*, likely driven by demographic contraction. To investigate this further, we reconstructed recent demographic histories by using GONE (Santiago et al., 2020). Results showed that *P. squamata* maintained a relatively high effective population size (median *N*_e_ ∼1.0 × 10^5^) until ∼20 generations ago, followed by a rapid collapse: within just two generations *N*_e_ plummeted from ∼4.2 × 10⁴ to 63, and then fell to fewer than 10 individuals (contemporary *N*_e_ = 2.7) (**Fig. 2E, Supplementary Figure 14–19**; see consistent estimates by multiple independent alternative approaches in **Supplementary Note 2** and bottleneck drivers in **Supplementary Note 3**). In contrast, *P. bungeana* maintained stable *N*_e_ (mean *N*_e_ = 594 ± 153; contemporary *N*_e_ = 561.5) over the past 200 generations, while *P. gerardiana* underwent a more recent decline, starting around 8 generations ago (mean *N*_e_ = 1909 ± 776; contemporary *N*_e_ = 92.5). The genome-wide population recombination rate (*ρ* = 4*N*ₑ*r*) also showed a severe reduction in *P. squamata* (**Fig. 2F**), consistent with the recent drastic decline in *N*ₑ. Because *ρ* reflects both the recombination rate per generation (*r*) and *N*ₑ, this reduction primarily indicates a loss of recombination efficacy due to the extremely small contemporary population size.

Most Pinaceae species are highly outcrossing, with selfing occurring at appreciable rates in only a few taxa, such as *Picea chihuahuana*, a species with small, isolated populations (selfing rate = 0.92) (Williams, 2009). This suggests that demographic collapse in *P. squamata* may have similarly enforced inbreeding and/or selfing. By genotyping offspring of wild trees, we estimated the selfing rate to be 0.82 (**Fig. 2G**), which supports this hypothesis. The average inbreeding coefficient *F*_IS_ for *P. squamata* is as high as 0.42, which is significantly greater than that of the other two pine species (**Supplementary Figure 20**). Inbreeding (including selfing) in such a severely reduced population likely exposed recessive deleterious alleles to purifying selection, thereby accelerating the erosion of genetic diversity and further reducing *N*_e_ (Dussex et al., 2023; Hedrick and Garcia-Dorado, 2016).

Given these demographic reconstructions, we next examined how the extreme population decline shaped efficacy of purifying selection and accumulation of genetic load — both of which are central to understanding fitness consequences. The ratio of heterozygosity at zero-fold versus four-fold degenerate sites (*H*_0_/*H*_4_) provides a proxy for the efficacy of purifying selection, with lower values typically indicating stronger selection (Yang et al., 2018). Despite its extremely low genome-wide heterozygosity and *N*_e_, *P. squamata* exhibited significantly lower *H*₀/*H*₄ ratios than *P. bungeana* and *P. gerardiana* (**Fig. 2H**). This pattern differs from theoretical expectations that reduced *Nₑ* should elevate *H*₀/*H*₄ due to weakened purifying selection under strong genetic drift (Chen et al., 2017; Yang et al., 2018; Zhao et al., 2024). Instead, the reduced *H*₀/*H*₄ observed in *P. squamata* suggests that inbreeding may have enhanced the efficacy of purifying selection, facilitating the purging of deleterious alleles despite small population size. Windowed *π*₀/*π*₄ also revealed that half (∼ 50%) of windows in *P. squamata* fall below the values observed in the other two pines, although the large variation reflects strong drift (**Fig. 2I**). The interpretation of strong purifying selection is also consistent with the inferred distribution of fitness effects (DFE), where *P. squamata* shows a much higher fraction of strongly deleterious mutations (population selection coefficient 4*N*_e_*s* < –100) among predicted new mutations (**Fig. 2J**). Furthermore, *P. squamata* showed a markedly negative Tajima’s *D* (median = –0.66), lower than those of other pines (median = –0.06 and –0.34) (**Fig. 2K**), consistent with an excess of rare alleles under strong purifying selection but inconsistent with the expectations of a bottleneck (Tajima, 1989). These findings imply that purifying selection — likely reinforced by inbreeding — plays a key role in shaping the genome of *P. squamata*.

We next directly assessed mutational load to test whether purifying selection had driven purging of deleterious alleles and thus a reduced deleterious burden. We examined heterozygous and derived homozygous sites, and total derived alleles, for four functional categories: tolerated, moderate-effect deleterious, high-effect deleterious, and loss-of-function (LoF). Counts were normalized by the corresponding synonymous class to control for background mutation rates and SNP ascertainment bias (**Fig. 2L, Supplementary Figure 21**). *P. squamata* showed comparable levels of tolerated and deleterious mutations (with the exception of LoF mutations) in the heterozygous state compared to *P. bungeana* and *P. gerardiana* (**Supplementary Note 4**). In the homozygous state, *P. squamata* exhibited markedly elevated frequencies of tolerated and deleterious mutations compared to the other two pines, reflecting inbreeding-driven exposure of recessive deleterious alleles. Strikingly, however, loss-of-function (LoF) mutations remained at very low frequencies in the homozygous state. Together with significantly lower allele frequencies of highly deleterious mutations, these results strongly suggest efficient purging of severe-effect mutations by purifying selection (**Fig. 2L**). These patterns mirror findings in other small, inbred populations, such as snow leopards (Yang et al., 2025) and certain carnivores (Hasselgren et al., 2021), where historical bottlenecks and inbreeding have effectively removed strong deleterious load while allowing moderate-effect variants to persist. However, in *P. squamata*, the observed reduction in frequency of moderate-effect alleles relative to tolerated alleles (**Fig. 2L**) suggests that purging remains moderately effective in preventing their accumulation. Overall, these data revealed a threshold-dependent pattern of selection efficacy: strong-effect mutations are efficiently purged in *P. squamata*, while moderate-effect variants are purged less efficiently in the context of small *N*ₑ and high inbreeding. Despite this purging, the higher burden of homozygous deleterious variants — including both moderate and high-effect types— may result in constraints on long-term adaptive capacity.

### Significant genetic differentiation despite spatial proximity in *P. squamata*

The extremely small size of the *P. squamata* population provides a rare opportunity to investigate how population differentiation can arise under conditions of such small *N*_e_. Based on admixture, phylogenetic, and PCA analyses, the 33 wild individuals of *P. squamata* clustered into three geographically distinct groups: West (*n* = 18), East (*n* = 13), and North (*n* = 2) (**Fig. 3A–D, Supplementary Figure 22**). Identity-by-state (IBS) and identity-by-descent (IBD) analyses also supported this subdivision (**Supplementary Figure 23**).

**Figure 3.**
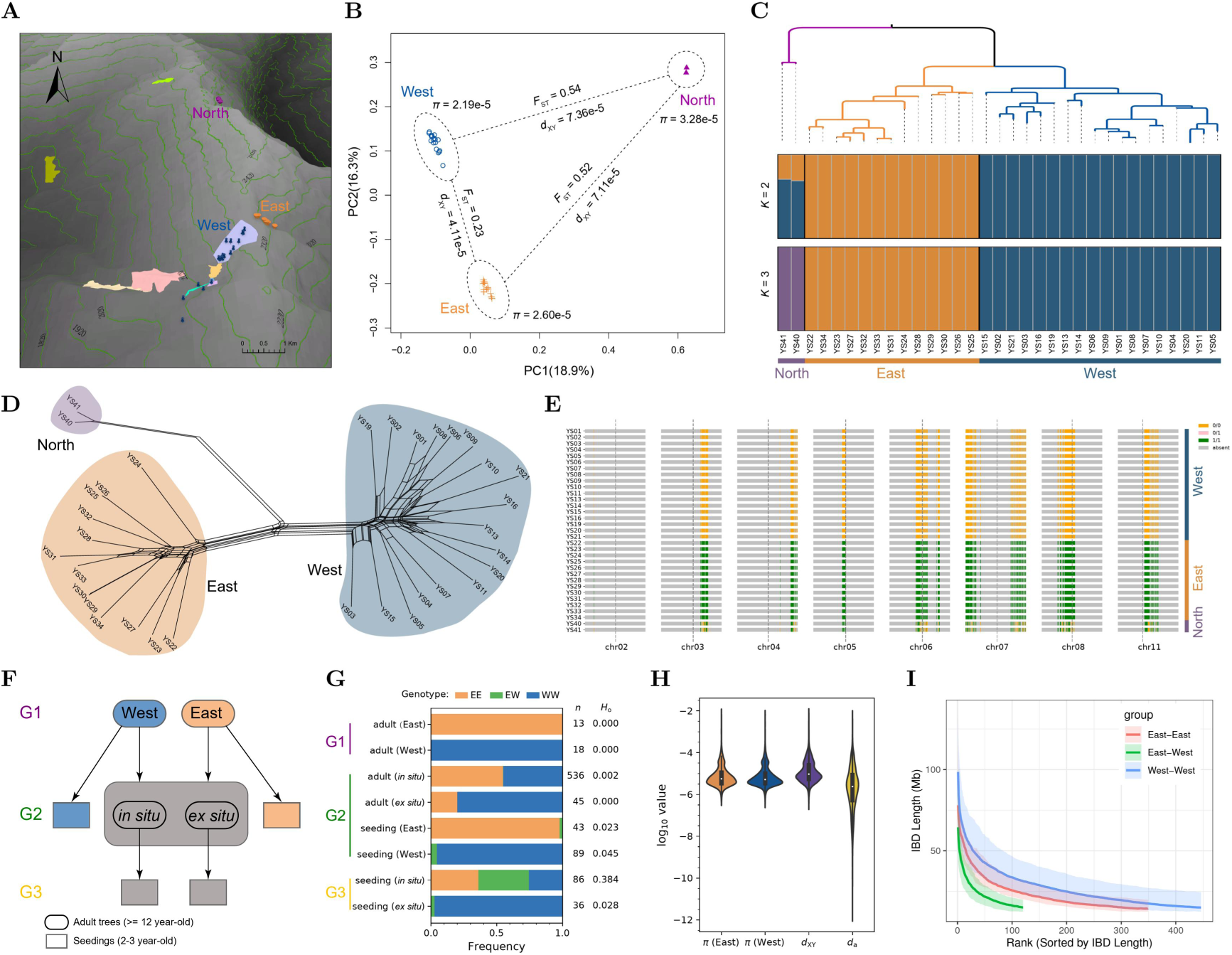
Population structure and differentiation within *P. squamata*. **A**) Geographical distribution of the three local populations. Colored points denote wild individuals. Irregular patches denote plantations from wild-collected seeds for *in situ* (population augmentation) conservation. Note: although some plantations overlaps with the West population, these trees are distant from wild individuals. East-slope plantations largely failed to reach adulthood; the North population occurs on cliffs and remains unfruitful. **B**) Principal component analysis (PCA). **C**) Phylogenetic tree and population structure (admixture). **D**) Split network. **E**) Distribution of fixed SNPs between East and West populations. Colored blocks represent genotypes (0/0, 1/1, 0/1). Dashed lines indicate centromeres. **F**) Pedigree and sampling scheme for genotyping population-specific alleles. G1 (generation 1) represents the initial wild individuals, first discovered and documented during the 1990s. From these wild sources, G2 (generation 2) individuals were established between 1995 and 2013 through artificial seeding for both *in situ* (population augmentation) and *ex situ* conservation. These individuals have since reached maturity. However, because parental origins were not systematically recorded during this early propagation phase, the G2 conservation stands consist of a genetic mixture with uncertain ancestry. To overcome these limitations and ensure traceability, our group initiated new propagation efforts in 2023 with explicit parental documentation. Seeds collected directly from the G1 wild populations yielded new G2 seedlings, while seeds harvested from the mature G2 conservation trees produced a third generation (G3). **G**) Genotype frequency of population-specific alleles in adult offsprings and seedlings. EE represents the homozygous genotype specific to the East population, WW denotes the homozygous genotype specific to the West population, and EW signifies the heterozygous genotype. *n*, number of tested individuals of each group. *H*o, the proportion of observed heterozygotes in each group. **H**) Nucleotide diversity (*π*) within populations, average number of nucleotide differences (*d*_XY_) and net nucleotide differences (*d*_a_) between East and West populations. **I**) Identity-by-descent (IBD) decay between East and West populations.

Although separated by less than 500 meters, the East and West populations of *P. squamata* showed no evidence of recent admixture or gene flow, as revealed by population structure and phylogenetic network analyses **(Fig.3A–C, Supplementary Figure 22B)**. Pairwise *F*_ST_ comparisons revealed extensive genomic differentiation, with multiple genomic islands approaching fixation (*F*_ST_ = ∼1). A total of 166,417 fixed SNPs (17,513 without missing genotypes) concentrated across seven chromosomes were identified (**Fig. 3E**). To verify whether adult offspring (generation 2, G2) that were propagated from wild-collected seeds had genetic admixture between East and West populations (G1), we further genotyped 581 G2 adult offspring at five fixed SNPs using the Kompetitive Allele Specific PCR (KASP) method, and detected only a single heterozygote individual (**Fig. 3F–G**), strongly supporting a lack of gene flow between the two populations.

Annotation of these fixed SNPs revealed that several were missense mutations in nucleotide-binding leucine-rich repeat (NLR) genes (**Supplementary Table 9**), leading us to suspect potential genic incompatibilities (Li and Weigel, 2021). However, genotyping 86 G3 seedlings from adult offspring *in situ* within mixed-genotype plantations (containing both EE and WW genotypes) revealed a high proportion of heterozygotes (38.4% EW; *P* = 0.12, χ^2^ test for Hardy-Weinberg equilibrium) (**Fig. 3F–G**), indicating no strong prezygotic barrier or early hybrid inviability. Nonetheless, postzygotic selection cannot be excluded, as fitness components (e.g., fertility, growth) in later growth were not assessed due to the long generation time of pines. In contrast, G2 seedlings from the East and West populations exhibited a low heterozygote frequency (2.3% and 4.5%, respectively), even though these wild populations are geographically close to the mixed-genotype plantations that have recently become potential pollen sources (**Fig. 3A**). This pattern likely reflects limited pollen dispersal, as the distances separating wild populations from the plantations remain considerably larger than the typical spacing among trees within the plantations.

Collectively, these results indicate that the observed genetic subdivision is unlikely to stem from intrinsic reproductive barriers. Instead, we propose that microgeographic isolation, amplified by severe demographic constraint explains the differentiation. Specifically, the ridge separating East and West populations may act as a barrier (**Fig. 3A**). Although direct estimates of pollen dispersal are unavailable for *P. squamata*, studies in other conifers have shown that pollen dispersal is highly restricted (∼50 m mean, <7% beyond 200 m) (Robledo-Arnuncio and Gil, 2005). Moreover, the combination of very small *N*_e_ and elevated inbreeding can heighten stochasticity and thus accelerate genetic drift and/or draft, rapidly fixing alleles and creating genomic islands of divergence within only a few generations.

Furthermore, the extremely small *d*_XY_ (absolute divergence) and *d*_a_ (net divergence) between East and West populations (mean *d*_XY_ = 4.28 × 10⁻⁵, median *d*_XY_ = 9.20 × 10⁻^6^; mean *d*_a_ = 1.88 × 10⁻⁵, median *d*_a_ = 1.43 × 10⁻^6^) suggest a very recent split following the diversity collapse (**Fig. 3H, Supplementary Note 2**). This is consistent with the estimated divergence time of ∼18 generations based on IBD tract lengths (**Fig. 3I, Supplementary Figure 24**), which closely followed the demographic decline estimated by the LD-based methods (**Fig. 2E**). Thus, a recent bottleneck coupled with limited dispersal and strong drift/draft likely led to the observed genetic subdivision.

### Genomic landscape of diversity and differentiation in *P. squamata*

Across the genome, population genetic metrics (*π*, *F*_ST_, *d*_XY_, Tajima’s *D*) exhibited clear bimodal distributions, consisting of one major peak and typically one minor peak (**Fig. 4**; histogram plots). This bimodal pattern indicates that, for each metric, most genomic regions cluster around one value class, while a minority around another. However, such consistency among regions is not uniform across metrics: different genomic regions can adopt distinct combinations of between-population (*F*_ST_ and *d*_XY_) and within-population (*π* and Tajima’s *D*) metrics (**Fig. 4**; Manhattan plots). Similar patterns were also generally observed for other diversity and differentiation metrics (**Supplementary Figure 25–26**). By contrast, equivalent analyses of two other pines (e.g., Fig. 2K) and other small plant populations, including *Acer yangbiense* (Ma et al., 2022), *Rhododendron* griersonianum (Ma et al., 2021), and *Malania oleifera* (Shen et al., 2024), recovered (nearly) normal distributions of these genetic metrics, consistent with theoretical expectations under neutral evolution.

**Figure 4.**
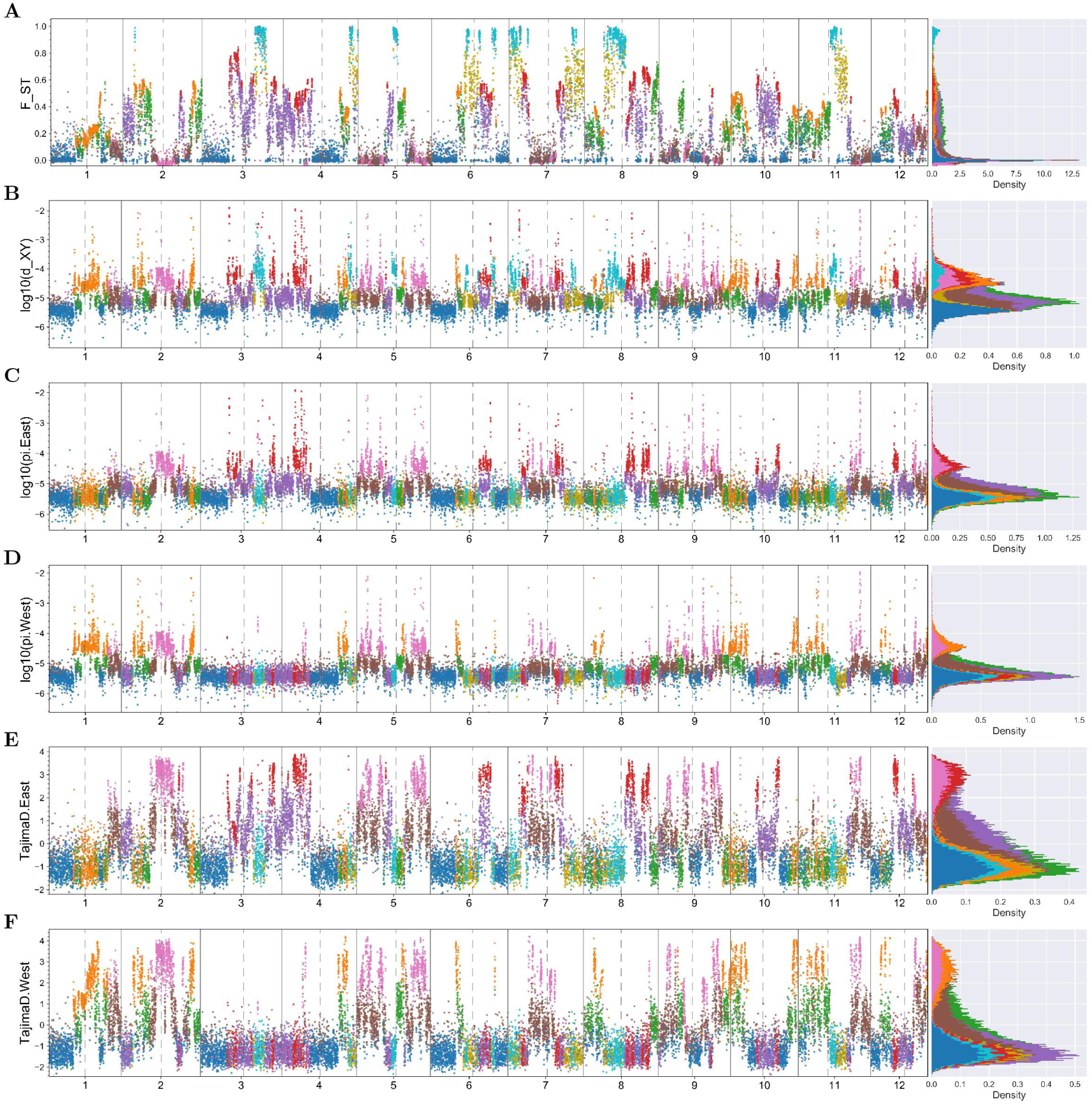
Genomic landscape of population differentiation in *P. squamata*. **A–F)** Manhattan plots and histogram plots of *F*_ST_ (**A**) and *d*_XY_ (**B**) between East and West populations, *π* of the East population (**C**) and West population (**D**), and Tajima’s *D* of the East population (**E**) and West population (**F**). Window size is 1 Mb. The dashed gray lines indicate centromere positions. Right panel showed histograms of each metrics, sharing the same Y axis with the corresponding Manhattan plots. All panels use the same cluster color scheme (see Fig. 5A).

To synthesize these heterogeneous patterns in *P. squamata*, we clustered non-overlapping 1-Mb windows with t-SNE (**Fig. 5A, Supplementary Figure 27**), which yielded nine clusters (Clusters 1–9; **Fig. 4, 5B–G, Supplementary Figure 28–34**). These clusters were characterized by consistent metric combinations (**Fig. 4, 5B–D, Supplementary Figure 28–34**). Clusters 1–5 occupy unique positions on the two-dimension site frequency spectrum (**Fig. 5F**), which we considered as primary. Clusters 6–9 mirror blends of clusters 2–5 with cluster 1, lying metrically intermediate and physically adjacent (**Fig. 4, 5B–F, Supplementary Figure 28–34**), implying origin via linkage disequilibrium (LD). At a broad scale, Clusters 1 and 2 displayed polarized values for between-population metrics (*F*_ST_ and *d*_XY_), yet similarly for within-population metrics (both low *π* and negative Tajima’s *D*); Cluster 3 showed similar *F*_ST_ values with Cluster 1, but inversely high *π* and positive Tajima’s *D*; Clusters 4 and 5 displayed opposite patterns for within-population metrics, with one population showing similar values to those of Cluster 1 and the other similar to those of Cluster 3 (**Fig. 4, 5B–F**). These relationships imply that driving forces underlying these primary clusters can operate similarly in some respects, but diverge in others.

**Figure 5.**
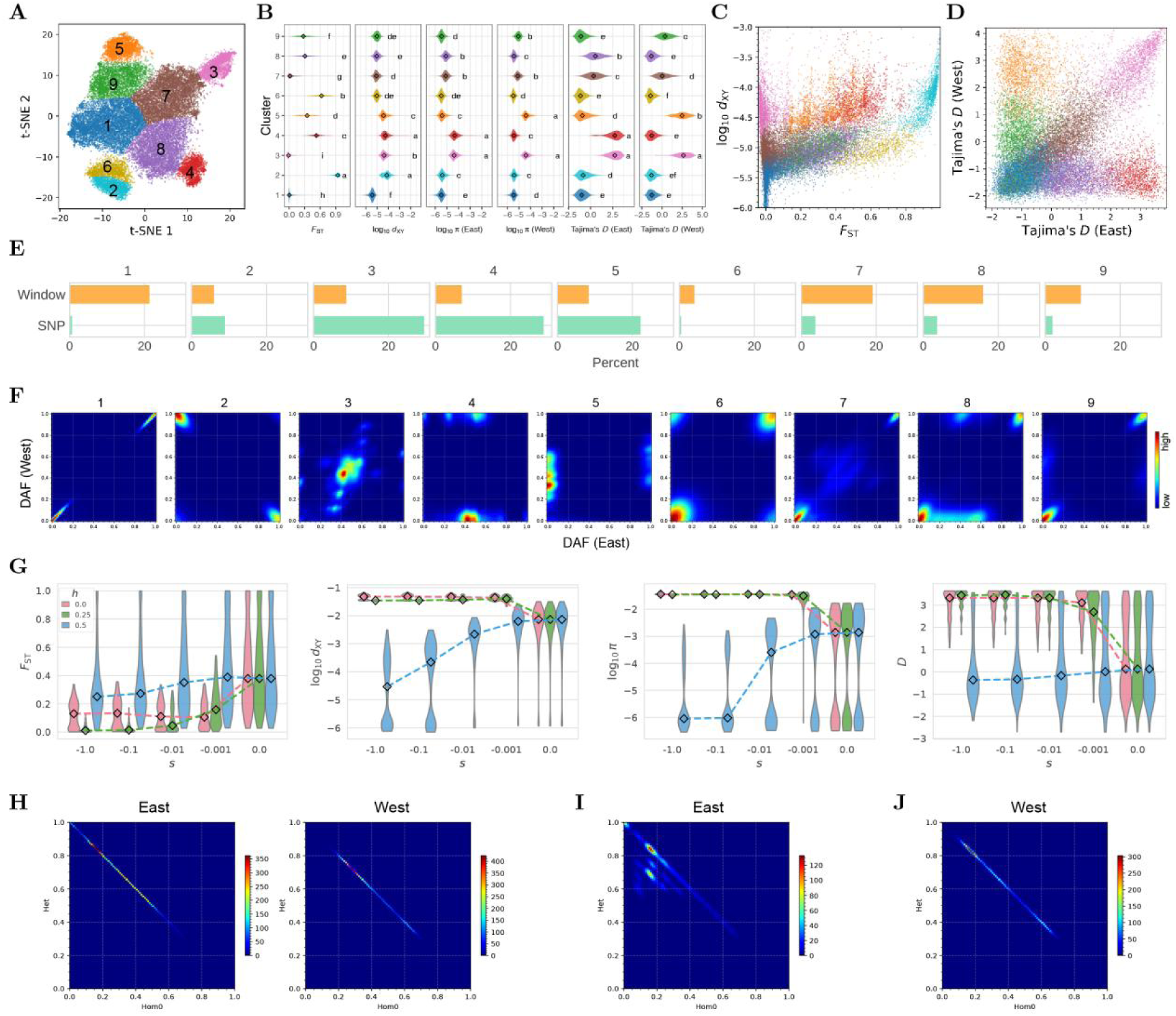
Genomic clusters of *P. squamata*. **A**) t-SNE clustering of 1-Mb genomic windows based on summary statistics, including *F*_ST_, *d*_XY_, multiple estimations of *θ* and neutrality tests. Nine clusters were identified and classified by their characteristic statistic profiles. **B**) Violin plots showing the distribution of summary statistics across clusters. **C**) Scatter plot of *F*_ST_ against *d*_XY_. **D**) Scatter plot of Tajima’s *D* of the East versus West population. **E**) Percent of the nine clusters calculated based on window and SNP counts. **F**) Unfolded Site Frequency Spectrum (SFS) between East and West populations for the nine clusters. DAF, derived allele frequency. **G**) Forward simulations confirm the differential impact of purifying selection on multiple metrics ( *F*_ST_, *d*_XY_, *π*, and Tajima’s *D*). *s*, selection coefficient; *h*, dominance coefficients. **H**) Frequency of heterozygote (Het) and one homozygote class (Hom0) among sites with excess heterozygosity (*F*_IS_ < -0.25) in both populations. The frequency of the other homozygote class (Hom1) is zero, when data points appear on the anti-diagonal. **I–J**) Frequency of heterozygote (Het) and one homozygote class (Hom0) among sites with excess heterozygosity (*F*_IS_ < -0.25) in East (**I**) and West (**J**) populations. Diamond symbols indicate median values in panels B and G.

Cluster 1 comprised the genomic majority (the major peak across multiple metrics), characterized by suppressed *d*_XY_ and near-zero *F*_ST_ between populations, and reduced *π* and Tajima’s *D* (negative) in both populations (**Fig. 4** and **5B–D**). This pattern is inconsistent with the general prediction under the scenario of neutral evolution (genetic drift dominates over natural selection across the whole genome), which would yield high *F*_ST_ with relatively intermediate *d*_XY_ and *π* (see also forward simulations in **Fig. 5G, Supplementary Figure 37–45** and **Supplementary Note 5**). It is also inconsistent with balancing selection and positive selection, as the former force would elevate *π* above the majority and the latter force would elevate *F*_ST_ but typically affect only small genomic regions (**Supplementary Figure 35–36**, see also **Supplementary Note 6**). Instead, the pervasive low *F*_ST_, low *d*_XY_, low *π*, and negative Tajima’s *D* across most of the genome points to strong purifying selection. Given the very low recombination rates in conifers (Stapley et al., 2017), background selection (BGS) likely extends the effects of purifying selection across large genomic regions. This process provides a coherent explanation for the major peak observed across multiple metrics. This view is also consistent with the observations of strong purifying selection in *P. squamata* (**Fig. 2**).

Cluster 2 corresponded to genomic islands of differentiation, exhibiting elevated relative divergence (*F*_ST_ ≈ 1) and absolute divergence (*d*_XY_) between the East and West populations (**Fig. 4A–B, 5B–C**). Such patterns can arise from three alternative scenarios: (i) divergent selection with gene flow, which would be expected to reduce *π*; (ii) inflow introgression, which would increase π; or (iii) divergent sorting of ancestral polymorphisms (Cruickshank and Hahn, 2014; Han et al., 2017). Because *π* in both populations was unchanged relative to the genomic majority (Cluster 1; **Fig. 4C–D**) and gene flow was not detected (**Fig. 3**), the first two scenarios are not supported. Instead, the elevated *d*_a_ (**Supplementary Figure 29**) suggests that these regions diverged prior to the population split, consistent with sorting of ancestral polymorphisms. Moreover, the low *π* and negative Tajima’s *D* within populations in Cluster 2 were similar to Cluster 1 (**Fig. 4, 5B–D**), indicating that BGS likely acted as the underlying force for each population. Under this scenario, BGS can retain different haplotypes in the two populations, producing near fixation differences (*F*_ST_ ≈ 1). Although neutral drift could also elevate *F*_ST_, it would be expected to increase *π* relative to BGS, which differs from the observed patterns.

Cluster 3 was characterized by high *d*_XY_ and low *F*_ST_ (near-zero) between populations, and elevated *π* and Tajima’s *D* (positive) in both populations (**Fig. 4 and 5B–D**). Although population contraction can generate positive Tajima’s *D*, it would be expected to elevate *F*_ST_ due to drift under neutrality, which differs from our observation of near-zero *F*_ST_. Instead, these patterns are consistent with balancing selection acting on sites shared by both populations. Balancing selection can maintain ancestral polymorphisms, thereby increasing *π* and *d*_XY_, while keeping *F*_ST_ low and producing positive Tajima’s *D*. However, the effectiveness of balancing selection in such small, inbred populations is questionable (Olito and Connallon, 2025), unless loci under selection confer exceptionally strong heterozygote advantage (true overdominance). Another, more plausible explanation is pseudo-overdominance (POD; or associative overdominance). Following purging of recessive deleterious homozygotes, haplotypes carrying complementary deleterious alleles can persist in heterozygous states, maintaining polymorphism across linked regions in small populations (Gilbert et al., 2020; Jin et al., 2021; Salson et al., 2025; Schou et al., 2017). We indeed observed one homozygote class absent at almost all sites with excess heterozygosity (*F*_IS_ < -0.25; **Fig. 5H**), consistent with the patterns of POD driven by purifying selection to eliminate homozygotes (Salson et al., 2025). Both POD and BGS stem from purifying selection on linked sites, but POD can dominate when selection is weak, dominance coefficients are low and recombination rate is low (Gilbert et al., 2020; Zhao and Charlesworth, 2016). Our forward simulations further support the differential impact of purifying selection on these various metrics across a range of dominance coefficients (**Fig. 5G, Supplementary Figure 37–45, Supplementary Note 5**). Specifically, the simulations demonstrated that low dominance coefficients (*h*) sharply increased the prevalence of POD, leading to the co-occurrence of elevated *π* and *d*_XY_ alongside suppressed *F*_ST_ (**Fig. 5G**), mirroring our empirical observations in Cluster 3.

Clusters 4–5: Clusters 4 and 5 exhibited contrasting within-population metrics (**Fig. 4 and 5B–D**). In Cluster 4, the East population displayed high Tajima’s *D* and *π*, whereas the West showed low values for both; in Cluster 5, this trend was reversed. These within-population metrics resemble those of Cluster 3 in the population with high values (e.g., those of Cluster 4 in East population) but resemble those of Cluster 1 in the other population with low values (**Fig. 4 and 5B–D, I–J**), suggesting POD acting within one population (resulting in high Tajima’s *D* and *π*) and BGS acting within the other population (resulting in low Tajima’s *D* and *π*). Alternatively, some of the contrasting signals could result from ongoing positive selection operating independently in each population, supported by deviations in Fay and Wu’s *H* (**Supplementary Figure 34**).

Collectively, these observations indicate that purifying selection on linked sites (i.e., BGS and POD) likely represents the primary driver shaping these genomic landscapes. We propose a unified model (**Fig. 6**) in which purifying selection is the central mechanism. Given the extremely small *N*_e_ (∼2.7), as few as two major haplotypes can be expected to persist within the population. If the same haplotype is retained in both populations after removing deleterious haplotypes by purifying selection (BGS), all metrics remain low (Scenario 1; **Fig. 6**). If different haplotypes are retained after BGS (Scenario 2; **Fig. 6**), *F*_ST_ and *d*_XY_ become high while *π* and Tajima’s *D* remain low. If complementary deleterious haplotypes are maintained (POD) in both populations (Scenario 3; **Fig. 6**) after eliminating homozygotes by purifying selection, *F*_ST_ remains low, while *d*_XY_, *π*, and Tajima’s *D* are high. If one population experiences BGS while the other experiences POD, Tajima’s *D* and *π* are high in one population but low in the other (Scenarios 4–5; **Fig. 6**). These five scenarios closely match the observed patterns in Clusters 1–5 (**Fig. 4–6**), while Clusters 6–9 likely represent linkage spillovers from Clusters 2–5. In summary, the two linked selection processes driven by purifying selection—BGS (eroding genetic diversity) and POD (maintaining diversity)—govern patterns of genetic diversity observed in P. *squamata*. Different combinations and orientations of these two linked selection regimes across populations generate heterogeneity in genetic differentiation. Concordant BGS or POD suppresses differentiation, whereas divergent BGS, or BGS in one population and POD in the other, accelerates differentiation. Under strong purifying selection and low recombination rates, these two linked selection processes can dominate genome evolution and generate highly heterogeneous genomic landscapes such as that reported here.

**Figure 6.**
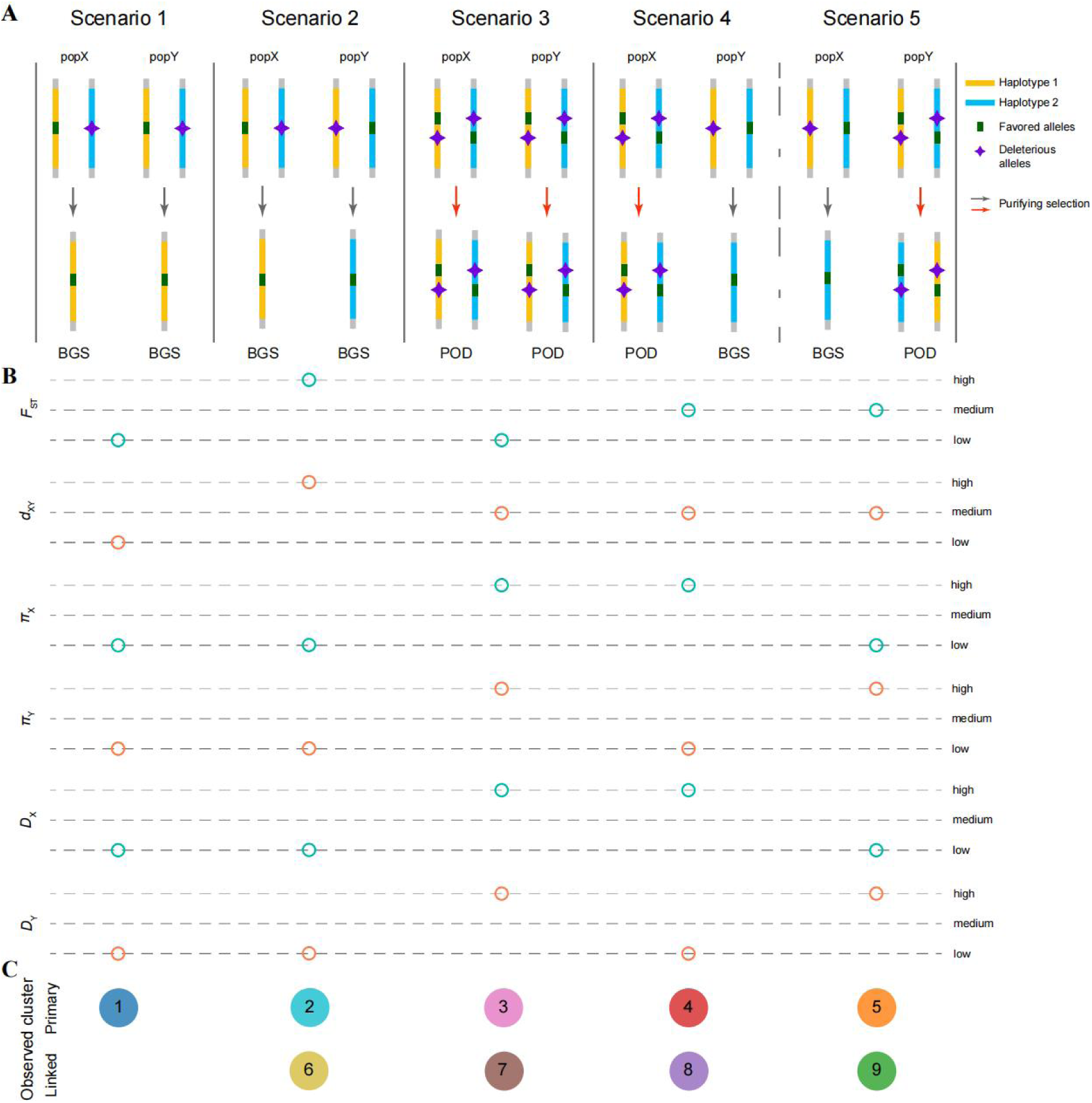
A unified ‘purifying selection-centric model’ to explain the formation of heterogeneous genomic patterns. **A**) Five linked selection scenarios driven by purifying selection in two populations (X and Y): Scenario 1 describes background selection (BGS) where the same ancestral haplotype is retained in both populations after the removal of deleterious alleles; Scenario 2 represents divergent BGS where different ancestral haplotypes are independently retained in each population; Scenario 3 illustrates pseudo-overdominance (POD), where complementary deleterious haplotypes are maintained in both populations due to the elimination of deleterious homozygotes; Scenarios 4–5 represent asymmetric modes where the two populations experience different selective outcomes, with one population characterized by BGS while the other undergoes POD. **B**) Predicted patterns of population genetic statistics (*F*_ST_, *d*_XY_, π, and Tajima’s *D*) across the five scenarios. Each statistic is categorized into three relative levels (high, medium, and low), with each scenario yielding a distinct combinatorial profile. **C**) The primary observed genomic patterns (Clusters 1–5) corresponding to the five scenarios. Clusters 6–9 represent intermediate states between the primary patterns (Clusters 2–5) and the baseline BGS pattern (Cluster 1), likely resulting from linkage disequilibrium.

## Discussion

Our data indicate that strong purifying selection has effectively purged deleterious mutations from *P. squamata* despite its extremely small *N*ₑ (approximately 2.7). Although purging has been widely documented in small populations (Dussex et al., 2023), the overall efficacy of purifying selection typically remains lower in small populations than in large ones (Feng et al., 2024; Lin et al., 2025; Yang et al., 2018). For instance, in *Ostrya rehderiana* (currently reduced to only five wild individuals), selection efficacy (indicated by *H*_0_/*H*_4_) is lower than in its widespread congeners, aligning with the expectations of the nearly neutral theory (Yang et al., 2018). In striking contrast to this pattern, *P. squamata* exhibited markedly higher purifying selection efficacy relative to its widespread congeners. Multiple lines of evidence support such genuine and effective selection (**Fig. 2H--M**). Among these, ratio-based metrics such as *π*₀/*π*₄ can be biased in populations far from equilibrium (Brandvain and Wright, 2016; Simons and Sella, 2016), but the population has persisted at small size for more than 4*N*ₑ generations, making equilibrium plausible. Furthermore, alternative metrics less sensitive to non-equilibrium — including distribution of fitness effects and derived deleterious allele frequencies (Brandvain and Wright, 2016; Simons and Sella, 2016) — yield concordant results.

Several interacting factors may contribute to the unexpectedly efficient selection observed in this system. First, the high selfing rate (approximately 0.8) enables systematic inbreeding, which can purge more efficiently than panmictic inbreeding because recessive deleterious alleles are more consistently exposed to selection in the homozygous state (Glémin, 2003). Although Glémin (2003) predicted that purging becomes ineffective below a threshold of *N*_e_*s* = –10, our forward simulations suggest that moderately deleterious mutations (mean s = −0.01) may still be effectively purged under the inferred demographic history (**Fig. 5G**), consistent with our empirical data (**Fig. 2L**) and previous simulations (Dussex et al., 2023). This discrepancy implies that the earlier theoretical work might underestimate the realized efficacy of purifying selection in small, highly inbred populations such as *P. squamata*. Second, the species’ life history may amplify per-generation selection intensity. Longevity generates overlapping generations that buffer random allelic fluctuations, while high fecundity coupled with extremely low seedling recruitment means that the vast majority of progeny fail to establish. Under such conditions, even modest fitness differences among offspring can be amplified—a dynamic conceptually similar to truncation selection, the most efficient form of directional selection (Crow and Kimura, 1979). In small, selfing maize populations, truncation selection has been demonstrated to maintain high selection efficiency despite strong genetic drift (Desbiez-Piat et al., 2021).

The efficient removal of strongly deleterious alleles may in turn have reshaped the specie’s realized mating system. Following purging of strongly deleterious alleles, inbreeding depression is expected to decline, making elevated selfing more tolerable for the progeny (Shalev et al., 2022). Unlike the pre-fertilization self-incompatibility systems common in angiosperms, conifers avoid selfing primarily through the embryo lethal system, a post-fertilization barrier driven by inbreeding depression caused by exposure of recessive lethal mutations (Williams, 2008; Williams, 2009). Given the limited pollen dispersal distance typical of pines (predominantly < 100 m), self-pollination (geitonogamy) rates are intrinsically high (Williams, 2007; Williams, 2009), yet strong inbreeding depression results in a very low proportion of detectable selfed offspring, as is common in conifers. Once deleterious mutations are purged and inbreeding depression is mitigated, however, viable selfed individuals become detectable (Busch, 2005; Qiao et al., 2010; Shalev et al., 2022). The observed selfing rate in *P. squamata* is therefore consistent with the finding that severely deleterious alleles have been substantially purged. Overall, the resulting dynamic is likely self-reinforcing: intrinsically high selfing sustains purging, and purging may help maintain the conditions that permit high selfing.

Beyond these consequences for the mating system, purifying selection at linked sites appears to have generated a markedly heterogeneous genomic landscape. Clustering analysis resolved distinct genomic modes dominated by background selection (BGS) and pseudo-overdominance (POD), with nucleotide diversity differing by an order of magnitude (**Fig. 4–5**), reflecting the dual nature of purifying selection in determining genetic diversity. Importantly, while these two distinct genomic landscapes show no clear correlation with recombination rates or centromere proximity, our simulations indicate they are largely determined by the dominance coefficients (*h*) of deleterious mutations (**Fig. 4–5, Supplementary Figure 37–45**). This also contrasts with patterns in other eukaryotes, where BGS strength typically correlates positively with recombination rate (Pouyet et al., 2018; Sella et al., 2009). Such BGS is likely the result of long-term, recurrent purifying selection, which typically exerts a more pronounced influence in large populations and has a limited impact in small populations (Charlesworth, 2012; Cutter and Payseur, 2013). However, *P. squamata* may represent a rare case where an extremely small population is significantly affected by BGS, potentially because: 1) strong purifying selection—even when acting on only a few highly deleterious sites—can extend to BGS across large genomic regions under conditions of extremely low recombination (Stapley et al., 2017); 2) over a short evolutionary timescale (approximately 20 generations), these large-scale haplotypes have not yet been repeatedly disrupted by extensive recombination.

Our study also provides broader insights into the conservation genetic theory. Traditionally, small *N*_e_ is associated with strong drift and reduced selection efficacy. Yet under high inbreeding in such small populations, we propose that the resulting enhanced purging can allow selection to outweigh drift. When combined with low recombination in conifers (Stapley et al., 2017), extensive linkage disequilibrium further amplifies this effect, extending selection (i.e., linked selection) across vast genomic regions. Under these conditions, the (nearly) neutral theory could not fully explain the observed patterns. This may be because the simplifying assumptions when applying the (nearly) neutral theory, such as random mating, non-overlapping generations, and independent loci, are systematically violated in *P. squamata*. Our purifying selection-centric framework (**Fig. 6**), which integrates the previous BGS and POD models, better explains the patterns observed in *P. squamata*. In future work, it is important to see if these patterns are replicated in other systems with very small *N*_e_ to determine if *Pinus* is exceptional, as well as to better understand the factors that enhance the efficacy of the purifying selection in very small populations.

In summary, our comprehensive population genomic analysis provides molecular evidence for strong purifying selection at the extreme edge of small *N*_e_, challenging the traditional expectations of conservation genetic theory that drift should overwhelm selection in small populations. High selfing and other traits likely enhance the efficacy of purifying selection. Successful purging, in turn, may have facilitated a turnover in the mating system from outcrossing to selfing. Strong purifying selection, combined with low recombination rates, likely plays a dominant role in shaping the highly heterogeneous genomic landscapes via linked selection (BGS and POD), further challenging the conventional view that drift governs genome evolution in small populations. These findings shed light on the evolutionary resilience of extremely small populations and provide insights into the genomic mechanisms that can potentially mitigate the risk of mutational meltdown and escape the drift-driven extinction vortex in critically endangered species.

### Limitations

Key limitations of this study relate to the reliance on assumed parameters and the need for experimental validation. First, our demographic and selection inferences assume a constant recombination map, which could biased the modelling. Second, comparisons of the fitness of seedlings derived from mixed plantations versus controlled crosses, could reveal whether the observed POD islands confer heterozygote advantage under natural conditions. Finally, achieving full resolution of the recombination landscape and megabase-scale haplotypes demands long-read sequencing of additional individuals, which was not financially feasible for these giant genomes.

## Methods

### Genome sequencing

Plant samples were collected from an individual (GL03) conserved *ex-situ* in Kunming Botanical Garden, Kunming Institute of Botany, Chinese Academy of Sciences. This tree was germinated in 2009 from wild-collected seeds, transplanted to the Kunming Botanical Garden in 2012, and first set fruit in 2021. Genomic DNA was extracted using a modified CTAB method (Doyle and Doyle, 1987). The genome was sequenced using various high-throughput sequencing technologies.

For the whole genome sequencing, genomic DNA (1 μg) was fragmented to sizes of 200–400 bp using a Covaris sonicator (Brighton, UK). These fragments were used to prepare short-read libraries according to the manufacturer’s protocols, and short-read sequencing was performed on the DNBSEQ-T7 platform (BGI lnc., Shenzhen, China) in paired-end 150 mode.

For long-read sequencing, genomic DNA was fragmented using the Megaruptor 3 shearing kit (Diagenode SA, Seraing, Belgium). Fragments less than 5 kb were selectively removed using the AMPure PB beads size selection kit (Pacbio, Menlo Park, CA, USA). The SMRTbell® prep kit 3.0 from Pacbio was used to prepare the sequencing libraries, which were then sequenced on the PacBio Revio system in high-fidelity mode. The Nanopore DNA library was prepared using SQK-LSK109 Kit (Oxford Nanopore Technologies, Oxford, UK), and sequenced on a Nanopore PromethION sequencer.

For Hi-C sequencing, we fixed young leaf material in a 2 % formaldehyde solution, with library preparation following a published protocol (van Berkum et al., 2010). In brief, the chromatin was digested with *Mbo*I, tagged with biotin-14-dCTP, and ligated. After ligation, the DNA underwent reverse cross-linking and purification, and was then sheared to 200-600 bp fragments. These biotin-labeled Hi-C fragments were enriched using streptavidin magnetic beads. After A-tailing and adapter ligation, the remaining Hi-C fragments were amplified using PCR (12–14 cycles) and sequenced on the DNBSEQ-T7 platform in PE150 mode.

Total RNA was isolated from various tissues using the R6827 Plant RNA Kit (Omega Bio-Tek, Norcross, GA, USA), according to the manufacturer’s instructions. For RNA-seq, libraries were prepared using the VAHTS Universal V6 RNA-seq Library Prep Kit. The libraries were then sequenced on the DNBSEQ-T7 platform (BGI lnc., Shenzhen, China) in paired-end 150 mode. For Iso-seq, full-length cDNA libraries were prepared for mixed samples using the cDNA-PCR Sequencing kit SQK-PCS109 (Oxford Nanopore Technologies, Oxford, UK) and sequenced on the Nanopore PromethION sequencer.

### Genome assembly

To facilitate genome assembly, *k*-mer spectra were first generated from short-read data using KMC v3.1.1 (Kokot et al., 2017). Genome size and heterozygosity were subsequently estimated from these k-mer profiles through findGSE v0.1.0 (Sun et al., 2017) and GenomeScope 2.0 (Ranallo-Benavidez et al., 2020). Initial contig assembly employed Hifiasm v0.19.5 (Cheng et al., 2024), which constructed a unitig graph and produced primary contigs by integrating both HiFi and ONT reads. To achieve chromosome-scale scaffolding, Hi-C reads were aligned against the contig assembly with Juicer v1.5.6 (Durand et al., 2016a). The resulting contact maps were processed through the 3D-DNA pipeline v180922 (Dudchenko et al., 2017) using ’--early-exit -m haploid -r 0’ parameters. Chromosome boundaries and assembly misjoins were then manually inspected and corrected within Juicebox v1.11.08 (Durand et al., 2016b), yielding a fully chromosome-scale assembly.

To further enhance assembly contiguity and resolve collapsed large-scale tandem duplications, unitig graph sequences were aligned to the draft assembly with minimap2 (Li, 2018). Gaps were subsequently filled and collapsed repeat regions resolved through manual inspection using Bandage v0.9.0 (Wick et al., 2015). Organellar genomes (chloroplast and mitochondrion) were independently assembled by GetOrganelle v1.7.5 (Jin et al., 2020) followed by manual refinement in Bandage v0.9.0 (Wick et al., 2015). The nuclear assembly underwent an additional polishing step with Nextpolish2 v0.1.0 (Hu et al., 2024), leveraging both HiFi and short-read data to correct remaining base errors. Finally, redundant haplotigs, ribosomal DNA arrays, and organelle-derived contaminant sequences were filtered using Redundans v0.13c (Pryszcz and Toni, 2016) under ’--identity 0.98 --overlap 0.8’ settings, supplemented by manual curation.

Assembly completeness and accuracy were evaluated through two complementary approaches. Benchmarking Universal Single-Copy Orthologs (BUSCO) scores were computed with miniBUSCO (Huang et al., 2023) against the Embryophyta odb10 reference set (Kriventseva et al., 2018), while base-level accuracy was assessed via k-mer spectrum analysis using Merqury v1.3 (Rhie et al., 2020) with high-quality short reads as the truth set.

### Genome annotation

Transposable element (TE) annotation began with constructing a de novo TE library from 1 Gb of randomly sampled genomic sequence using EDTA v1.9.9 (Ou et al., 2019) in sensitive mode (’--sensitive 1’). Genome-wide repeat masking was then performed by applying this custom library with RepeatMasker v4.0.7 (Tarailo-Graovac and Chen, 2009). Telomeric repeat arrays were located by scanning the assembly for the canonical plant telomere motif TTTAGGG.

Protein-coding gene annotation integrated three complementary lines of evidence: homology, transcriptomics, and ab initio prediction. For homology support, a comprehensive dataset of 315,505 non-redundant protein sequences was assembled from ten phylogenetically diverse plant species: *Amborella trichopoda* (Käfer et al., 2021)*, Cycas panzhihuaensis* (Liu et al., 2022), *Ginkgo biloba* (Liu et al., 2021), *Gnetum montanum* (Wan et al., 2021), *Metasequoia glyptostroboides* (Fu et al., 2023), *Pinus lambertiana* (Crepeau et al., 2017), *Pinus tabuliformis* (Niu et al., 2022), *Taxus chinensis* (Xiong et al., 2021), *Torreya grandis* (Lou et al., 2023), and *Welwitschia mirabilis* (Wan et al., 2021). Transcript evidence was derived from RNA-seq reads aligned with hisat2 (Kim et al., 2015) and Iso-seq reads mapped with minimap2 v2.24 (Li, 2018), followed by transcript assembly using StringTie v2.1.5 (Pertea et al., 2015) and Scallop v0.10.5 (Shao and Kingsford, 2017). These transcript assemblies informed initial gene structure models through PASA v2.4.1 (Haas et al., 2003). For ab initio prediction, AUGUSTUS v3.4.0 (Stanke et al., 2008) parameters were iteratively trained over five rounds using full-length gene models derived from PASA output, and SNAP (Korf, 2004) was trained on the same reference set. All evidence streams were combined within the MAKER2 v2.31.9 pipeline (Holt and Yandell, 2011), which jointly considered *ab initio* predictions (AUGUSTUS and SNAP), transcript alignments, and protein homology. A consensus gene set was subsequently generated by EvidenceModeler v1.1.1 (Haas et al., 2008), merging the independent outputs of PASA and MAKER2. TE-encoded protein domains, detected by TEsorter v1.4.1 (Zhang et al., 2022), were masked during the EVM integration step to prevent spurious annotations. Low-confidence gene models were filtered by aligning predicted protein sequences against transcript and homology evidence with BLAT (Kent, 2002); models were retained only when exhibiting >80% coverage and >98% identity to transcripts, or >80% coverage and >35% identity to homologous proteins. Gene structures were further refined by incorporating UTR boundaries and alternative splicing isoforms via PASA. Models with fewer than 50 amino acids, missing initiation or termination codons, internal stop codons, or ambiguous bases were discarded. The final annotation quality was benchmarked using BUSCO v5.3.2 (Simão et al., 2015) against the embryophyta odb10 lineage dataset.

Non-coding RNA genes were systematically catalogued across three categories. Transfer RNAs (tRNAs) were detected by tRNAScan-SE v1.3.1 (Lowe and Eddy, 1997); ribosomal RNA (rRNA) loci were identified with Barrnap v0.9 (https://github.com/tseemann/barrnap), retaining only full-length annotations after removing partial hits. Additional non-coding RNA families were discovered through homology searching against the Rfam database using RfamScan v14.2 (Nawrocki et al., 2015).

Functional characterization of the predicted proteome employed three independent strategies. First, eggNOG-mapper v2.0.0 (Huerta-Cepas et al., 2017) assigned orthologous groups by querying the eggNOG database restricted to Viridiplantae, enabling transfer of Gene Ontology (GO) terms and KEGG pathway annotations. Second, DIAMOND v0.9.24 (Buchfink et al., 2015) was run with ’--evalue 1e-5 --max-target-seqs 5’ to search against the Swiss-Prot, TrEMBL, NR, and TAIR10 protein databases. Third, protein domains and functional motifs were catalogued using InterProScan v5.27 (Jones et al., 2014), which interrogated the PRINTS, Pfam, SMART, PANTHER, and CDD databases consolidated within the InterPro framework.

### Chromatin immunoprecipitation sequencing (ChIP-seq)

The CENH3 protein sequences of *Pinus* spp. (Plackova et al., 2024) were used as references to identify homologs in *Pinus squamata* using BLAT v36 (Kent, 2002). A polypeptide “ARSKTVPARKKSGSGNAASC” was designed as an antigen, based on the identified CENH3 variants. The peptide is synthesized, emulsifed with an adjuvant, and injected into rabbits for immunization. After 7–10 weeks, blood samples were collected, and serum was harvested. Antibodies were purified via peptide-specific affinity chromatography. The specificity and reactivity against the target antigen were validated using Enzyme Linked Immunosorbent Assay (ELISA), and those against CENH3 in the plant samples were validated using Western blots. Fresh leaves were crosslinked with 1.25% formaldehyde, followed by glycine quenching, to preserve protein-DNA complexes. Chromatin DNA was mechanically sheared into fragments (200--1500 bp) using sonication. The chromatin is immunoprecipitated overnight with the CENH3-specifc antibodies, followed by precipitation of the antibody-antigen complexes with additional Salmon Sperm DNA/Protein A Agarose. Following multiple washes to remove non-specific bindings, the complexes were decrosslinked, and DNA was extracted for sequencing library construction. The libraries were sequenced using Illumina NovaSeq 6000 platform. Reads were aligned to the genome sequences using BWA-MEM v0.7.17 (Li, 2013). The CENH3 enrichment was quantifed using bamCompare from the DeepTools package (Ramírez et al., 2016) (‘--operation log2 --ignoreDuplicates --minMappingQuality 30 --scaleFactorsMethod None --normalizeUsing CPM’).

### Centromere identification

Centromere positions were initially identified from the Hi-C interaction heatmap, generated using Juicer v1.5.6 (Durand et al., 2016a) with Hi-C data. Inter-chromosomal signals around the centromere positions are characterized in the Hi-C heatmap of chromatin with Rabl configuration, and peri-centromere regions were identified in this way (Marie-Nelly et al., 2014). Tandem repeats (TR) in the peri-centromere regions were identified using TRF (Benson, 1999) and TRASH (Wlodzimierz et al., 2023), and the putative centromere positions were defined based on the commonly enriched regions of a 148 bp motif. The centromere positions were finally confirmed using the ChIP-seq data.

### Genome resequencing and SNP calling

We sampled wild individuals of *Pinus squamata* (n = 33) from Yunnan Province, *P*. *bungeana* (n = 10) from Shanxi Province, and *P*. *gerardiana* (n = 9) from Xizang Province, respectively, in China. Genomic DNA (1 μg) was fragmented to sizes ranging from 200 to 400 bp using a Covaris sonicator (Brighton, UK). These fragments were used to prepare short-read libraries according to the manufacturer’s protocols, and short-read sequencing was performed on the DNBSEQ-T7 platform (BGI lnc., Shenzhen, China) in paired-end 150 mode. The NGS data of *Pinus albicaulis* (Neale et al., 2024) were downloaded from SRA to serve as an outgroup.

Raw reads were filtered using Fastp v0.19.3 (Chen et al., 2018). Paired-end clean reads were aligned to the reference genome using BWA-MEM v0.7.17 (Li, 2013). SAMtools v1.17 (Danecek et al., 2021) was used to sort binary alignment map (BAM) format files and remove duplicate reads. BCFtools v1.17 (Danecek et al., 2021) was employed to genotype all sites, including variants and non-variants (parameters: ‘-q 5’). We then employed vcftools v0.1.15 (Danecek et al., 2011) to filter sites with the following criteria: (1) sites with coverage depth above 2.5 * average coverage depth were discarded after investigating the coverage distribution; (2) sites with more than two alleles or with an average mapping quality quality (MQ) less than 40. (3) MNPs (multiple nucleotide polymorphisms) and insertion/deletions (InDels) were excluded. (4) genotypes with a depth below 4× or a genotype quality score <20 were redefined as missing; (5) sites with a missing rate >20% were removed. This yielded 16,644,074,599 sites (dataset 1), including 1,521,523,280 SNPs and 15,122,551,319 non-variants. To further reduce reference bias, sites within TE regions or with heterozygosity greater than 0.7 in any species were excluded, yielding 325,277,611 SNPs and 4,518,776,513 non-variants (dataset 2). Note that, in the downstream analyses, dataset 2 was used as basis for three-species comparison to reduce reference bias, while dataset 1 was used as basis for analyses within *P. squamata* to retain more sites. The genotypes of *P. squamata* were extracted from dataset 1, resulting in 4,305,505 SNPs (dataset 3; vcftools parameters: --max-missing 0.8 --mac 1). We further extracted 1,639,084 non-singleton SNPs with minor allele count greater than 2 (dataset 4; vcftools parameter: --mac 3) and 313,269 putative independent SNPs (dataset 5; vcftools parameter: --thin 2000).

### Inbreeding

Based on dataset 2, runs of homozygosity (ROH) were estimated by bcftools (‘roh -G30’) (Narasimhan et al., 2016). Only ROH longer than 100 kb were kept. We calculated the fraction of runs of homozygosity (*F*_ROH_), which is equal to the total length of ROH divided by genome effective length. Heterozygosity (*H*) of each sample was also counted from the this dataset.

### Estimation of demographic history

Recent demographic history (within the last 200 generations) was inferred from patterns of linkage disequilibrium (LD) among SNPs using GONE (Santiago et al., 2020). The recombination rate was set to 0.12 cM/Mb according to an updated genetic map of *Pinus lambertiana* (3600 cM / 30 Gb) (Weiss et al., 2020). There were too many SNPs for *Pinus bungeana* and *Pinus gerardiana* to run GONE, so we randomly subsampled 20% SNPs for both species, according to the recommendation by the software author. More methods for estimating demographic history and contemporary *N*_e_ can be found in Supplementary Methods 1–2.

### Selection and Recombination inference

The ancestral genomic sequences were reconstructed using an empirical Bayes method implemented in IQ-TREE 1.6.5 (Nguyen et al., 2015) (see more details in Supplementary Method 3) . Folded site frequency spectrum (SFS) was estimated using realSFS (Korneliussen et al., 2014). DFE were modeled using FastDFE with 1000 bootstraps (Sendrowski and Bataillon, 2024). Fourfold degenerate (4d) sites were considered as neutral, and zerofold degenerate (0d) sites were considered as selected. Parametrization was conducted for mixture of a gamma and exponential distribution.

Signatures of positive selection were detected with IHH12, iHS, nSL, XP-EHH, and XP-nSL statistics using selscan (Szpiech and Hernandez, 2014). CLR (Degiorgio et al., 2015) and XP-CLR (https://github.com/hardingnj/xpclr) methods were also used to identify signals of positive selection. Balancing selection was detected using betascan (Siewert and Voight, 2017).

Population recombination rate (*ρ*) was estimated using FastEPRR (Gao et al., 2016).

### Detection of deleterious mutations

Deleterious mutations were identified using the Sorting Intolerant From Tolerant (SIFT) algorithm (Vaser et al., 2016). The ancestral genomic sequences were used to build the SIFT database to reduce reference bias (Ma et al., 2022). The TrEMBL plant database was used to search for orthologous genes, and the SIFT scores were calculated based on the degree of conservation among loci. Based on dataset 2 (which included low-frequency variants), SNPs in the coding regions were categorized as synonymous, deleterious (SIFT score < 0.05), tolerated (SIFT score ≥ 0.05), or loss-of-function (start loss, stop gain, and stop loss) using SIFT4G (Vaser et al., 2016). Deleterious mutations were further subdivided into two categories: highly deleterious (SIFT score = 0) and moderately deleterious (SIFT score > 0). The low confidence sites and “NA” sites were excluded. The total number of derived alleles is based on counting each heterozygous genotype once and each homozygous-derived genotype twice (Yang et al., 2018). The count of each category was standardized by the corresponding number of synonymous sites in the same individual (Wang et al., 2024).

### Population structure

Population structure was inferred using ADMIXTURE v1.3.0 (Alexander and Novembre, 2009). Principal component analysis (PCA) was performed using GCTA v1.94.1 (Yang et al., 2011). A maximum-likelihood phylogenetic tree was reconstructed using IQ-TREE 1.6.5 (Nguyen et al., 2015). SplitsTree v6.1.16 was used to reconstruct phylogenetic networks (Huson and Bryant, 2024). Identity by state (IBS) was calculated by PLINK v1.9.0 (Purcell et al., 2007). These analyses were based on the LD-pruned SNPs (dataset 5).

### Split time estimation

Identity-by-descent (IBD) blocks were detected using RefineIBD v4.1 (window=100000 overlap=10000 ibdtrim=40 ibdlod=3) (Browning and Browning, 2013). IBD blocks with length less than 1.5 cM were discarded. Recent split time between two populations was estimated based on IBD length distributions (exponentially distributed) using the formula (Browning, 2008):

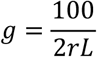

where *g* is the number of generations after split, *r* is recombination rate (0.12 cM/Mb here), and *L* is the mean length of IBD tracts (23.6 Mb here).

### Genetic diversity and differentiation estimation

Pairwise fixation index (*F*_ST_), absolute divergence (*d*_XY_), and nucleotide diversity (π) were estimated using pixy v1.2.7 (Korunes and Samuk, 2021). Estimations of *θ*, including pairwise diversity (*π* or *θ*_P_) (Nei and Li, 1979), Watterson’s *θ* (*θ*_W_) (Watterson, 1975), *θ*_H_ (Fay and Wu, 2000), *θ*_F_ (Fu and Li, 1993), *θ*_L_ (Zeng et al., 2006), and neutrality tests, including Tajima’s *D* (Tajima, 1989), Fay and Wu’s *H* (Fay and Wu, 2000), Fu and Li’s *D* and *F* (Fu and Li, 1993), and Zeng’s *E* (Zeng et al., 2006), were conducted using ANGSD v0.921 (Korneliussen et al., 2014). ANGSD was also used to calculate *F*_ST_ to confirm the results of pixy. These analyses were based on 1-Mb windows and the datasets comprising non-variants: dataset 1 for comparison among the *P. squamata* populations, or dataset 2 for comparison among the three species. The net nucleotide differences between two populations, *d*_a_, were calculated by formula (Cruickshank and Hahn, 2014):

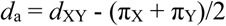

The values of *d*_XY_, *d*_a_ and *θ*, were log10-transformed due to the low value and potential log-normal distributions.

Under neutral theory, two isolated populations expects:

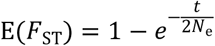

where *t* is the split generations for the populations. Assuming *t* = 18 and *N*_e_ = 2.7, we expect *F*_ST_ = 0.96.

### Clustering and classification of genomic windows

Based on the above summary statistics—including *F*_ST_, *d*_XY_, five estimators of *θ*, and four neutrality tests—we clustered 1-Mb non-overlapping winows using t-Distributed Stochastic Neighbor Embedding (t-SNE) implemented in Rtsne package (https://rdocumentation.org/packages/Rtsne/versions/0.17). The number of windows was 28,467. After investigating the distribution, we empirically and roughly classified the nine clusters by cutoffs as following.

Cluster 1: *F*_ST_ <= 0.1 & log_10_ *d*_XY_ < -5.3;

Cluster 2: *F*_ST_ >= 0.8;

Cluster 3: *F*_ST_ <= 0.1 & log_10_ *d*_XY_ >= -4.75;

Cluster 4: *F*_ST_ > 0.1 & *F*_ST_ < 0.8 & log_10_ *d*_XY_ >= -4.75 & Tajima’s *D*.West <= Tajima’s *D*.East;

Cluster 5: *F*_ST_ > 0.1 & *F*_ST_ < 0.8 & log_10_ *d*_XY_ >= -4.75 & Tajima’s *D*.West > Tajima’s *D*.East;

Cluster 6: *F*_ST_ >= 0.6 & log_10_ *d*_XY_ >= -4.75;

Cluster 7: *F*_ST_ <= 0.1 & log_10_ *d*_XY_ < -4.75 & log_10_ *d*_XY_ >= -5.3;

Cluster 8: *F*_ST_ > 0.1 & *F*_ST_ < 0.6 & log_10_ *d*_XY_ < -4.75 & Tajima’s *D*.West <= Tajima’s *D*.East; Cluster 9: *F*_ST_ > 0.1 & *F*_ST_ < 0.6 & log_10_ *d*_XY_ < -4.75 & Tajima’s *D*.West > Tajima’s *D*.East.

These clusters were subsequently refined by manual inspection and reassigned using Support Vector Classifier (SVC) implemented in sklearn (https://scikit-learn.org/). For better visualization and to reduce noises, we removed singletons with classification differing from both their adjacent windows (24,049 windows retained).

### SNP genotyping using KASP

Five diagnostic SNPs were genotyped via Kompetitive Allele Specific PCR (KASP) (**Supplementary Table 10**). Genomic DNA was extracted from samples and diluted to 5–10 ng/µL. KASP assays were performed in 10 µL reactions containing 5 µL of 2× KASP v4 Flu-Arms Mix, 0.5 µL of primer mix (two allele-specific forward primers and one common reverse primer), and 4.5 µL of DNA template. Thermal cycling was conducted on a Bio-Rad CFX96 system with an initial denaturation at 95°C for 10 min, followed by 10 cycles of 95°C for 15 s and 61°C (decreasing by 0.6°C per cycle) for 45 s, and then 36 cycles of 95°C for 15 s and 55°C for 30 s. Endpoint fluorescence was measured using FAM and HEX filters, and genotypes were determined based on cluster separation in a scatter plot, with homozygous samples showing single-color signals and heterozygous samples displaying dual signals.

### Genotyping using RNA-seq

RNA-seq was performed for 122 individuals. Reads were filtered using fastp v0.19.3 (Chen et al., 2018), and then mapped to the reference genome using HISAT2 v2.1.0 (Kim et al., 2015). SNPs were then genotyped using BCFtools v1.17 (Danecek et al., 2021). SNPs were filtered using a relatively relaxed criterion by vcftools v0.1.15 (Danecek et al., 2011) (‘--minGQ 10 --minDP 4 --min-alleles 2 --max-alleles 2 --remove-indels --max-missing 0.5’), yielding 100,267 SNPs of high quality. The dataset was used to confirm the KASP results and to infer selfing rate.

### Identification of selfing progeny

A total of 73 G2 individuals from the RNA-seq dataset were used to infer selfing, by comparing their genotypes with those of the G1 West and East individuals. For each SNP of a G2 individual, we identified whether >= 1 and 2 of the alleles matched the alleles of a given G1 individual, which are the expectations for either parent and selfing parent, respectively. The rates of these two conditions (denoted as *p*_1_ and *p*_2_) were calculated. If both *p*_1_ and *p*_2_ > 0.99, it was considered as selfing. Our data showed clear clustering for selfing, outcrossing within West or East populations, and outcrossing between West and East populations (Fig. 2G).

### Forward simulations

We performed forward-time, individual-based simulations to investigate the genomic processes of BGS and POD using SLiM 5.0 (Haller and Messer, 2023). The per-bp mutation rate (*μ*) in all simulations was 3.5e-8 (Zhao et al., 2024). Individuals consisted of one chromosome of 5Mb. Each chromosome was subject to only one recombination rate (*r*) 0.12 cM/Mb. The simulations modeled a gamma distribution for deleterious fitness effects with mean selection coefficient (*s*) equal to (−0.001, -0.01, -0.1, -1) and shape parameter 1. Mutations were only either neutral or deleterious, with the proportion of deleterious mutations fixed to be 1% (*P*_del_) of the total mutations occurring at any given point in time. Across replicates we varied the dominance of deleterious mutations (*h* = 0.5, 0.25, 0; additive, partially recessive, and fully recessive, respectively). Our simulated demographic history mirrored the inferred one: *T* = 20 generations ago a bottleneck reduced the population to *N* = 10 individuals; 15 generations ago it split into two populations, approximating the post-bottleneck *N*_e_ we estimated. After the bottleneck we set the selfing rate (*α*) to 0.5. To quickly approach the expected polymorphism for *N*_ANC_ = 1e5 (*π =* 0.014) before the bottleneck, we initially simulated an epoch for 10*N*_ANC_ generations with reduced *N*_ANC_ by a factor of 100 and increased *μ* by 100 times. The simulations were independently repeated 1000 times.

We further explored the parameter space by varying mutation rate *μ* (1e-9, 5e-9, 1e-8, 5e-8 and 1e-7), recombination rate *r* (0.01, 0.1, 1, 10 and 100 cM/Mb), selfing rate *α* (0, 0.2, 0.5, 0.8, 1), population size *N* (10, 20, 50, 100, 1000), bottleneck time *T* (10, 20, 50, 100, 1000), proportion of deleterious mutations *P*_del_ (0.01%, 0.1%, 1%, 10%), shape of gamma distribution of DFE (0.5, 1, 5, 10), ancestral population size *N*_ANC_ (1e3, 1e4, 1e5, 1e6) and lag time (Δ*T*) between bottleneck and split (5, 15, 45, 85, 985).

## Statistical analysis

For two-sample comparisons, we used the Wilcoxon rank-sum test. For multiple-sample comparisons, we used analysis of variance (ANOVA) with Tukey’s HSD (Honestly Significant Difference) post-hoc test, with Bonferroni correction applied for multiple comparisons.

## Data Availability

All the raw sequencing reads are deposited in Genome Sequence Archive (GSA) under Bioproject accession number PRJCA014196. The genome assembly and annotations are deposited in Figshare (DOI: 10.6084/m9.figshare.26103571).

[All the data will be released prior to publication.]

## Acknowledgments

We thank the staff of Yaoshan Mountain National Nature Reserve and colleagues at Kunming Institute of Botany, Chinese Academy of Sciences for help in sampling (with special thanks to Gui-Fen Luo, Xue-Lian Qin and Ji-Gui Zhang). We thank Profs. Martin Lascoux, Xiao-Ru Wang, Jian-Feng Mao, Li He, Martin Lysak, Michael Lynch, Yun-Peng Zhao, Qin Qiao, Jing-Li Zhang, Bin Tian, Hong Ma, De-Yuan Hong, Hai Ren, Yuan-Ming Zhang, Qing-Feng Wang, Hong-Wen Huang, Da-Yong Zhang, Qing-Yin Zeng, and Yong-Ping Yang for helpful discussions and/or critical comments.

## Author contributions

Y.P.M., W.Z., R.G.Z., X.Q.W., and W.B.S. conceived and designed the study. H.S., R.F.L., D.T.L., Z.J.L., H.Y., Y.Q.W., X.F.L., H.Y.S., M.J.Z., K.H.J., and Y.P.M. sampled the plants and performed experiments. R.G.Z., H.L., R.F.L., and W.Z. analyzed the data. R.G.Z., H.L., R.F.L., and H.S. prepared figures and tables. R.G.Z. and W.Z. drafted the manuscript; Y.P.M., R.G.Z., L.R., R.M., X.Q.W., H.S., W.B.S., H.L., J.C., and W.Z. revised the manuscript; all authors approved the final manuscript.

