## supplemental information for "Purifying selection purges harmful variants in the rarest pine"

### Supplementary Information

#### Supplementary Notes

##### Supplementary Note 1. Divergence among three pine species

Principal component analysis (PCA) separated *P. squamata*, *P. gerardiana*, and *P. bungeana* into three distinct clusters along PC1, with *P. squamata* forming a tight cluster and the other two species exhibiting broader within-species variation (**Supplementary Figure 13A**). Consistent with this pattern, Admixture analysis supported species-level separation at K = 3, while K = 4 revealed finer-scale substructure (**Supplementary Figure 13B**). Genetic differentiation among the species was high (mean $F_{ST}$ = 0.90 and mean $d_{XY}$ =0.015), consistent with a previous study based on RNA-seq data[1]. A split network analysis further confirmed the deep divergence of *P. squamata* from the other species (**Supplementary Figure 13C**).

##### Supplementary Note 2. Contemporary $N_e$ and historical demographic dynamics

To confirm the contemporary and recent fluctuations in effective population size ($N_e$), we performed demographic inferences using updated LD-based methods currentNe2 and GONE2, which consistently corroborated the GONE results. Our analyses revealed an extremely low contemporary $N_e$ (in the single digits) for the studied pine species, following a severe population bottleneck that occurred approximately 20 generations ago (**Supplementary Figure 15A--B**). To account for the potential confounding effects of population structure, we adopted two strategies: (i) implementing the -x parameter in the analysis and (ii) performing independent $N_e$ estimations for each sub-population. Both approaches yielded results (**Supplementary Figure 15C--E**) that were highly congruent with the species-wide analysis (i.e., treating all samples as a single population for each species), suggesting that population structure had a negligible confounding effect.

While population-specific estimates indicated slight temporal discrepancies in the onset of the bottleneck between the East and West populations (**Supplementary Figure 15D--E**), these differences do not necessarily imply independent bottleneck events. Instead, such variation likely arises from stochastic differences between the datasets or differential biases in tracing associated with post-split historical trajectories. Crucially, if the two populations had undergone independent bottlenecks following their divergence, a high level of ancestral polymorphism at the time of the split would be expected to result in elevated $d_{XY}$ values. However, the observed low $d_{XY}$ in our data contradicts this hypothesis, supporting a shared demographic bottleneck or more complex post-divergence interactions.

All coalescent-based approaches, including Stairway Plot, PSMC, fastsimcoal2, and DILS, also identified a recent population bottleneck (**Supplementary Figure 16--19**). However, the coalescent-based methods subsequently suggested a dramatic population expansion, a signal that diverges from the LD-based results and directly contradicts the critically low contemporary census population size ($N_c$). While this expansion is consistent with a genome-wide excess of rare alleles

(**Fig. 2K**), such an overrepresentation likely stems from linked selection triggered by intense purifying selection rather than true demographic growth. Under the assumption of neutrality, these methods can misinterpret the rare-allele excess as an expansion artifact.

Conversely, historical $N_e$ estimated from LD-based methods is largely unaffected by natural selection[2]. In addition, the GONE software assumes random mating; thus, population structure and selfing in *P. squamata* may introduce bias into $N_e$ estimates[3,4]. However, our results accounting for population structure remained robust, consistently inferring a recent severe bottleneck (**Supplementary Figure 15C–E**). Moreover, methods such as currentNe2 also account for mating system[5]. Therefore, biases from these LD-based methods are likely limited.

Accordingly, for historical $N_e$ inference, we did not adopt coalescent-based results, but instead relied on LD-based estimates from GONE, indicating that the bottleneck in *P. squamata* began approximately 20 generations ago, without subsequent population recovery (**Fig. 2E**).

The timing of these demographic events (bottleneck and split time) also varied, with coalescent-based estimates (dozens to hundreds of generations; **Supplementary Figure 16--19**) being times older than those derived from LD and IBD-based analyses (~20 generations). This temporal discrepancy may arise because mutation-based dating lacks sensitivity over short timescales where mutation accumulation is minimal. Furthermore, the presence of differentiated genomic islands—likely formed by the differential sorting of ancestral polymorphisms—can inflate split-time estimates since the origin of these polymorphisms predates the actual divergence. Nonetheless, IBD-based estimates may also be inflated by pseudo-overdominance (POD), which truncates IBD segments.

Ultimately, despite these inherent uncertainties in temporal dating, the combination of extremely low $N_c$, reduced genetic diversity, and low $d_{XY}$ collectively reinforces our conclusion: the species underwent a recent, severe bottleneck, followed by population split, with no subsequent recovery. This scenario, primarily inferred by LD and IBD-based methods (see main text), provides a more biologically plausible explanation for the observed genomic patterns than the models suggested by coalescent-based approaches.

### Supplementary Note 3. Population bottleneck timing and anthropogenic context

Our data reveal a sharp population bottleneck approximately 20 generations ago. The generation time for *P. squamata* likely lies between 50 and 15 years[6,7]. This gives a date range for the bottleneck of between ~1,000 and ~300 years ago. This period began with intensified economic activity in Southwest China and ended with a period marked by historical records of logging, mining, and agriculture in the region. These lines of evidence suggest that human activities likely played a key role in driving this collapse. Following the initial decline, inbreeding depression may have further exacerbated population collapse in the short term, but also enabled intense purifying selection to eliminate strongly deleterious mutations, thereby reducing the burden of strongly deleterious alleles. In the long term, however, the realized load — due to homozygous exposure of moderately deleterious variants — would be expected to increase. This dual dynamic may collectively lower fitness and adaptive potential[8], which could ultimately constrain its recovery despite the fact that the most severe mutations have already been purged.

#### Supplementary Note 4. Purging in *P. gerardiana*

Interestingly, *P. gerardiana*, despite its recent demographic decline and reduced $N_e$, exhibited relatively high burdens of moderate and highly deleterious alleles, particularly in the homozygous state (**Fig. 2E and 2L)**. This indicates that purging in *P. gerardiana* has likely been less effective than in *P. squamata*, potentially due to its more recent or less severe population bottleneck and lower levels of inbreeding. This contrast underscores the importance of both demographic timing and mating system in shaping purging efficacy and the retention of genetic load.

Compared with *P. bungeana*, *P. gerardiana* exhibits the opposite frequency pattern for highly deleterious genotypes: heterozygous genotypes are less frequent, whereas homozygous genotypes are more frequent (**Fig. 2L)**. This pattern reflects conversion from masked load to realised load, consistent with the recent population decline in *P. gerardiana*, suggesting an urgent need for conservation intervention to mitigate genetic erosion and the accumulation of deleterious mutations in the homozygous state.

#### Supplementary Note 5. Forward simulations

Based on our integrated demographic inferences, we performed forward simulations using SLiM to evaluate the impact of linked selection (**Fig. 5G**). Our simulations demonstrate that introducing a specific fraction of deleterious mutations ($s < 0$) can readily generate distinct genomic patterns: background selection (BGS) and pseudo-overdominance (POD). The emergence of these patterns is primarily governed by the dominance coefficient ($h$) of the deleterious mutations. A high dominance coefficient ($h$ = 0.5) tends to produce BGS, characterized by reduced nucleotide diversity ($\pi$) and inter-species divergence ($d_{XY}$) relative to neutral settings ($s$ = 0), alongside relatively lower $F_{ST}$ and Tajima's $D$. Conversely, a low dominance coefficient ($h$ = 0 or 0.25) yields the POD pattern, marked by significantly elevated $\pi$ and $d_{XY}$, low $F_{ST}$, and high Tajima's $D$. These distinct patterns remain robust across a wide range of parameter settings (**Supplementary Figure 37--45**).

Among the evaluated statistics, $d_{XY}$ serves as the most diagnostic indicator for distinguishing these evolutionary modes (POD, BGS and neutrality). Except for minor exceptions, $d_{XY}$ robustly segregates into three discrete tiers with order-of-magnitude differences: high (POD, $h$ = 0 or 0.25), intermediate (neutral), and low (BGS, $h$ = 0.5). While $\pi$ is also affected, it is more susceptible to confounding factors; for instance, as split time increases, neutral $\pi$ may converge toward BGS levels, whereas $\pi$ of POD remains consistently high. Furthermore, while POD exhibits clear deviations from neutrality in $F_{ST}$ and Tajima's $D$, BGS often overlaps with neutral expectations, making it difficult to identify using these metrics alone. This highlights $d_{XY}$ as the core diagnostic metric for identifying linked selection in empirical data.

Our empirical observations closely align with these simulated patterns. Cluster 1, which we putatively identified as dominated by BGS, exhibits the lowest $d_{XY}$ in our dataset, consistent with the $h$ = 0.5 simulation results. Similarly, Cluster 3 matches the simulated POD mode across all indices, including near-zero $F_{ST}$ and high $\pi$, $d_{XY}$, and Tajima's $D$. However, clusters 2, 4, and 5 likely involve divergent selection; since our simulations utilized constant parameters, they do not reproduce the specific differentiation patterns observed in these clusters, such as the high $F_{ST}$ and

$d_{XY}$ seen in Cluster 2. Additionally, while the simulations successfully reproduce the observed trends, numerical discrepancies in π and $d_{XY}$ (particularly the order-of-magnitude difference in Cluster 3) suggest that our simplified demographic model may not fully capture the absolute values observed in the empirical data.

### Supplementary Note 6. Natural selection at linked sites

Due to linkage disequilibrium, natural selection—including purifying, positive, and balancing selection—inevitably generates linked selection. There are three main forms of linked selection: background selection (BGS), pseudo-overdominance (POD), and selective sweeps. The first two are driven by purifying selection, whereas selective sweeps are driven by positive selection[9]. Selective sweeps often produce similar signatures to background selection, such as reduced π and Tajima's *D*. However, selective sweeps are likely rare in *P. squamata* for several reasons: (1) beneficial mutations are far fewer in number than deleterious and neutral variants[10]; (2) selection coefficients of beneficial mutations are generally weak, in contrast to the abundance of lethal deleterious mutations with $s = -1$[10], and strong genetic drift caused by extremely small $N_e$ further weakens the efficacy of positive selection (requiring $s \approx 0.1$ to achieve $4N_es \approx 1$); (3) intense purifying selection also interferes with positive selection, potentially masking or overriding its effects[10]; (4) limited new mutations and minimal environmental change during the short post-bottleneck period (~20 generations) restrict opportunities for adaptive selection. As a consequence, reduced efficacy of positive selection may limit local adaptation and compromise the evolutionary potential to respond to future climate change.

Consistent with this inference, within-population selective sweep statistics (nSL, CLR, etc.) are approximately normally distributed, with few outliers indicative of positive selection (**Supplementary Figure 35C–H and 36C–D**). By contrast, between-population statistics including XP-EHH and XP-CLR tend to identify Clusters 4 and 5 as showing signals of positive selection in the West and East populations, respectively (**Supplementary Figure 35A–B and 36A–B**). This likely represents a false inference driven by contrasting diversity patterns. For example, Cluster 4 exhibits higher π in the East population but lower π in the West population, which mimics a selective sweep in the West population that reduces diversity and increases homozygosity relative to the East. However, this selective sweep scenario conflicts with other genomic signatures, whereas strong background selection over short time scales better explains these patterns.

Ongoing selective sweeps occurring independently within each population may nevertheless be hidden among signals of balancing selection or pseudo-overdominance, supported by negative Fay & Wu's *H* values in high-π regions (**Supplementary Figure 26A–B**).

Additionally, signals detected by betascan are nearly normally distributed, with few outliers indicative of balancing selection and no clear association with the nine clusters (**Supplementary Figure 36E–F**). This further supports our interpretation that genomic regions with elevated within-population π (Clusters 3, 4, and 5) are dominated by pseudo-overdominance rather than balancing selection.

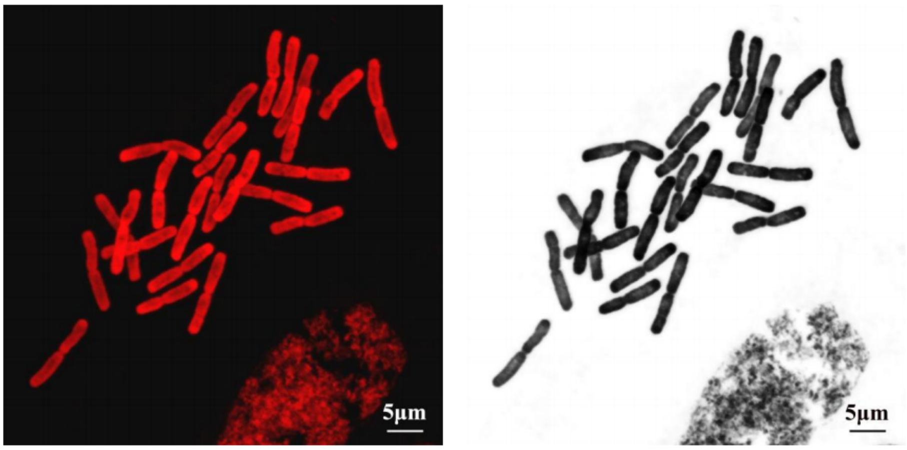


**Supplementary Figure 1. Chromosome morphology at metaphase of mitosis under the light microscope.**

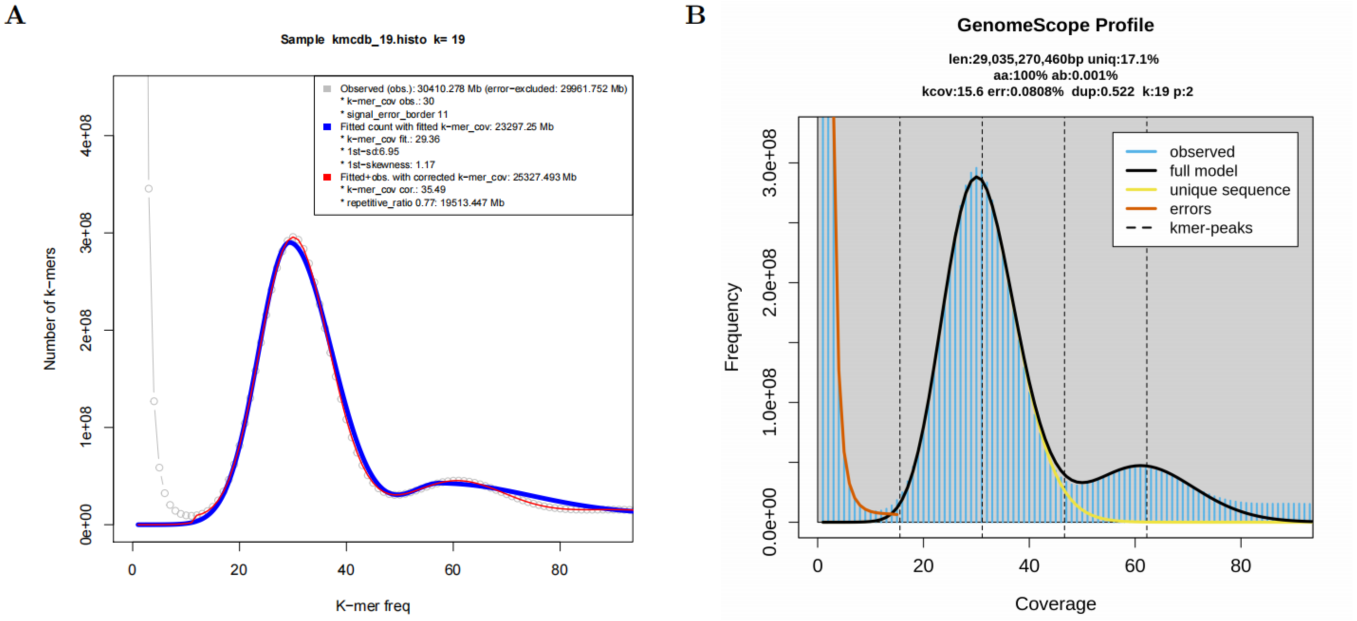


**Supplementary Figure 2.** ***K*-mer profiles to estimate genome size and heterozygosity rate using findGSE (A) and GenomeScope2 (B).** The *k*-mer size is 19 bp.

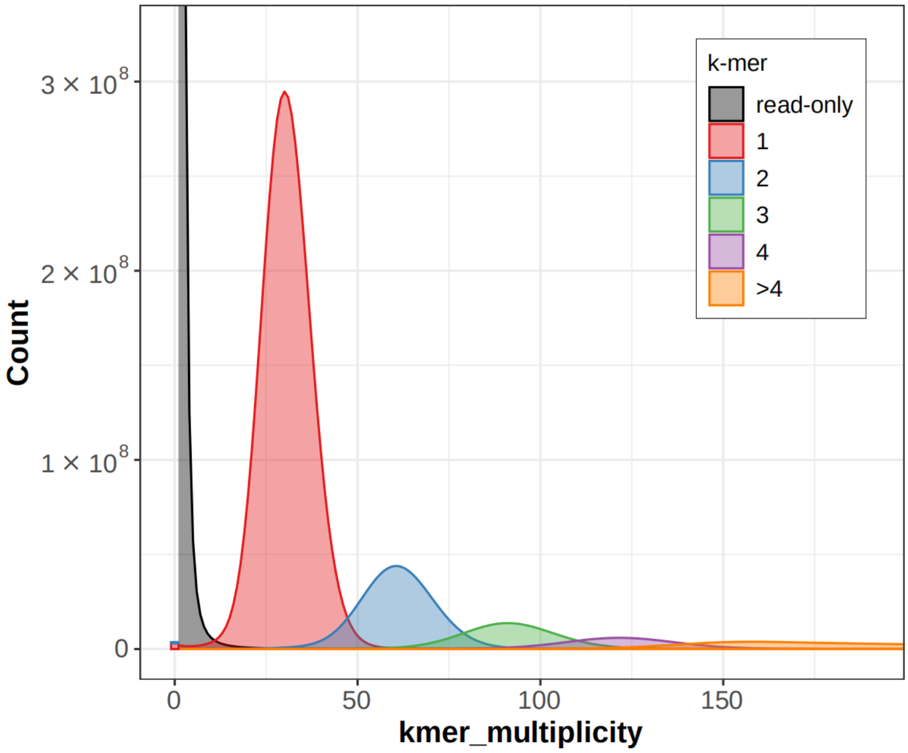


**Supplementary Figure 3. *K*-mer comparison between genome assembly and short reads.**

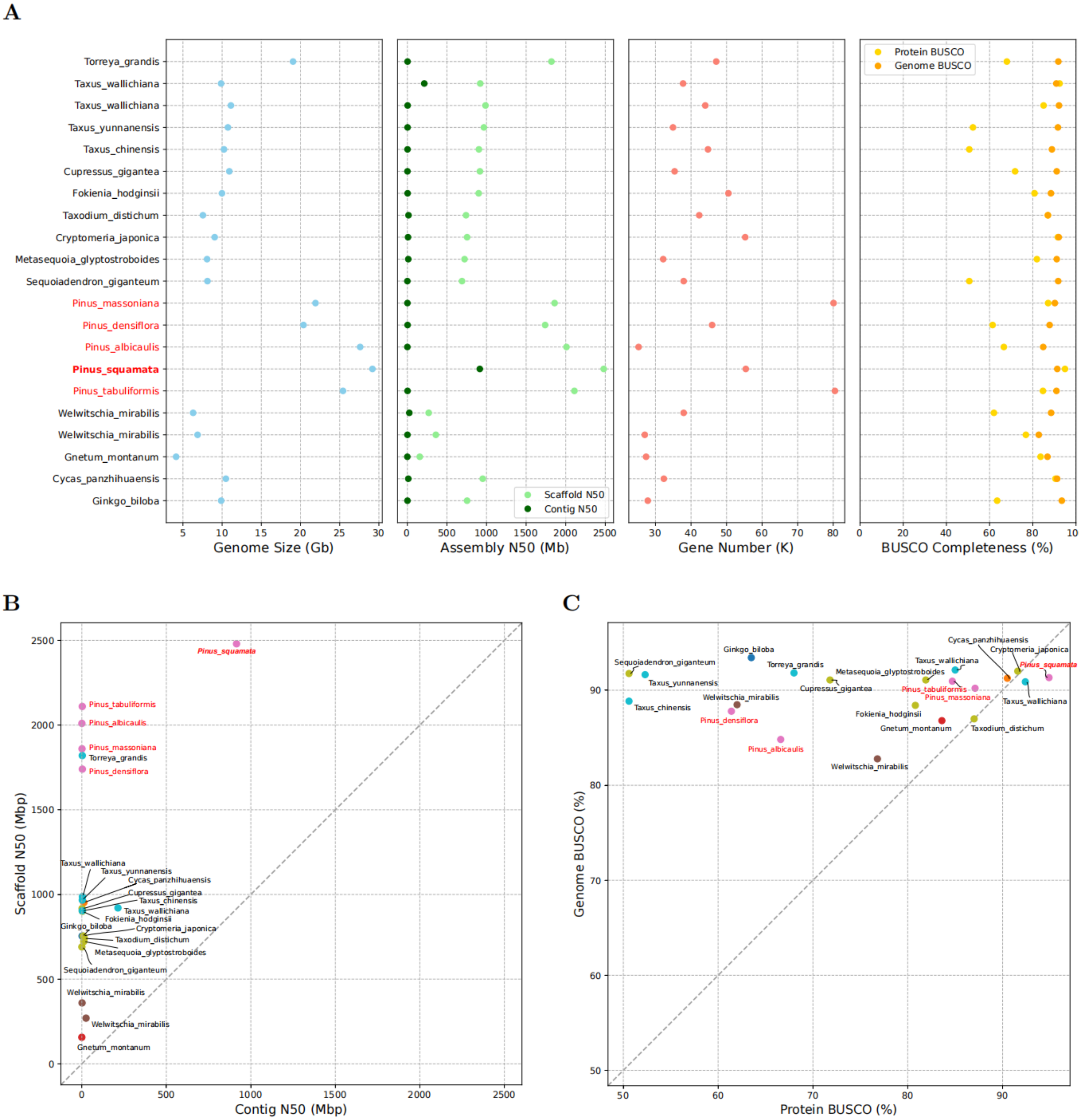


**Supplementary Figure 4. Comparison of genome assembly and gene annotation among *P. squamata* and published chromosome-level gymnosperm genomes. A)** Comparison of genome size, assembly N50, gene number, and BUSCO completeness. **B)** Comparison between contig N50 and scaffold N50; assemblies where the contig N50 approaches the scaffold N50 indicate higher continuity. **C)** Comparison between genome BUSCO and protein BUSCO; a protein BUSCO score close to or exceeding the genome BUSCO score suggests superior annotation completeness.

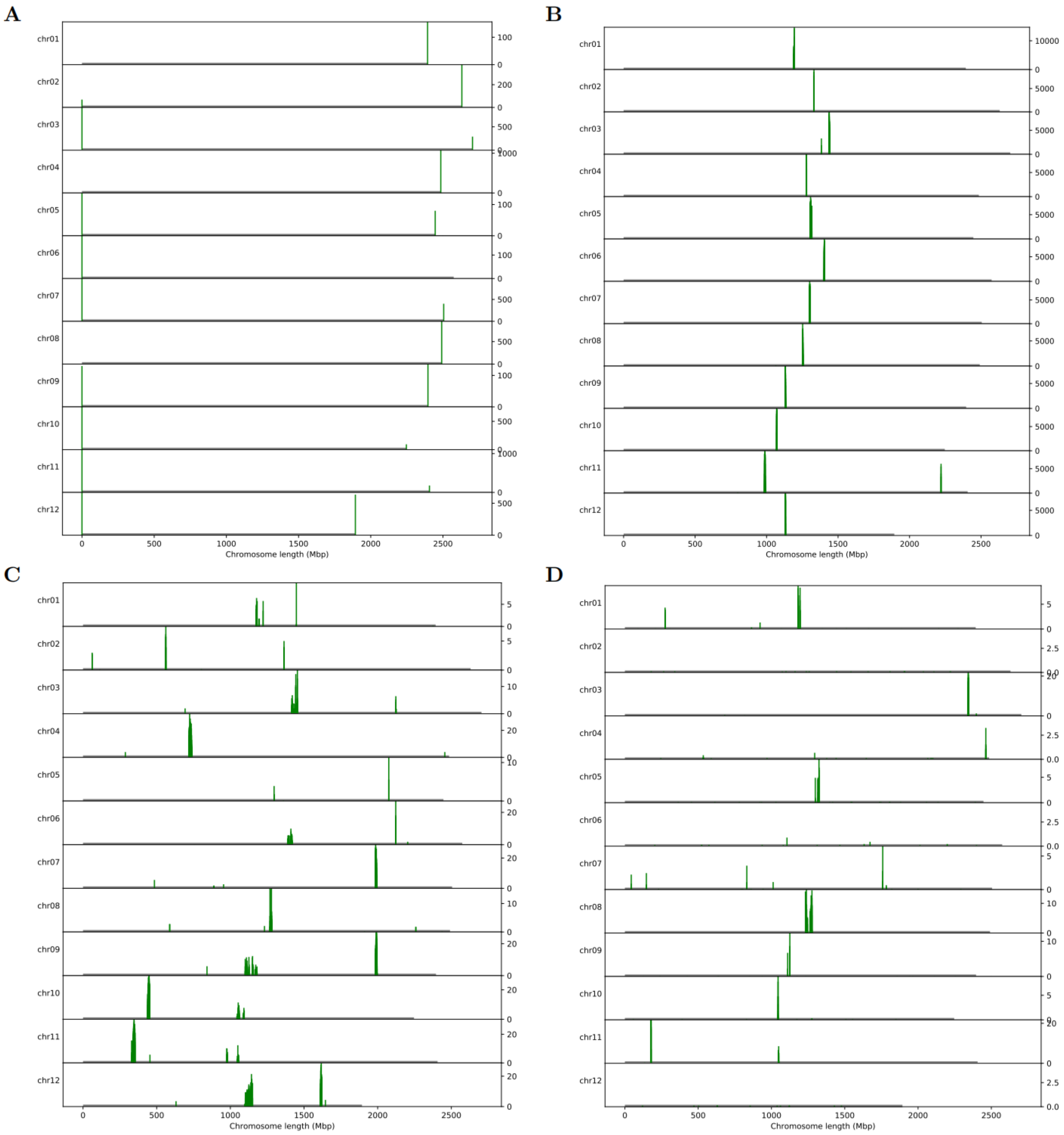


**Supplementary Figure 5. Locations of telomere repeat (TTTAGGG) (A), putative centromere tandem repeat (148 bp) (B), 45S rDNA (C) and 5S rDNA (D) sequences.** The y axes indicate the number of these sequences in 20 kb sliding windows.

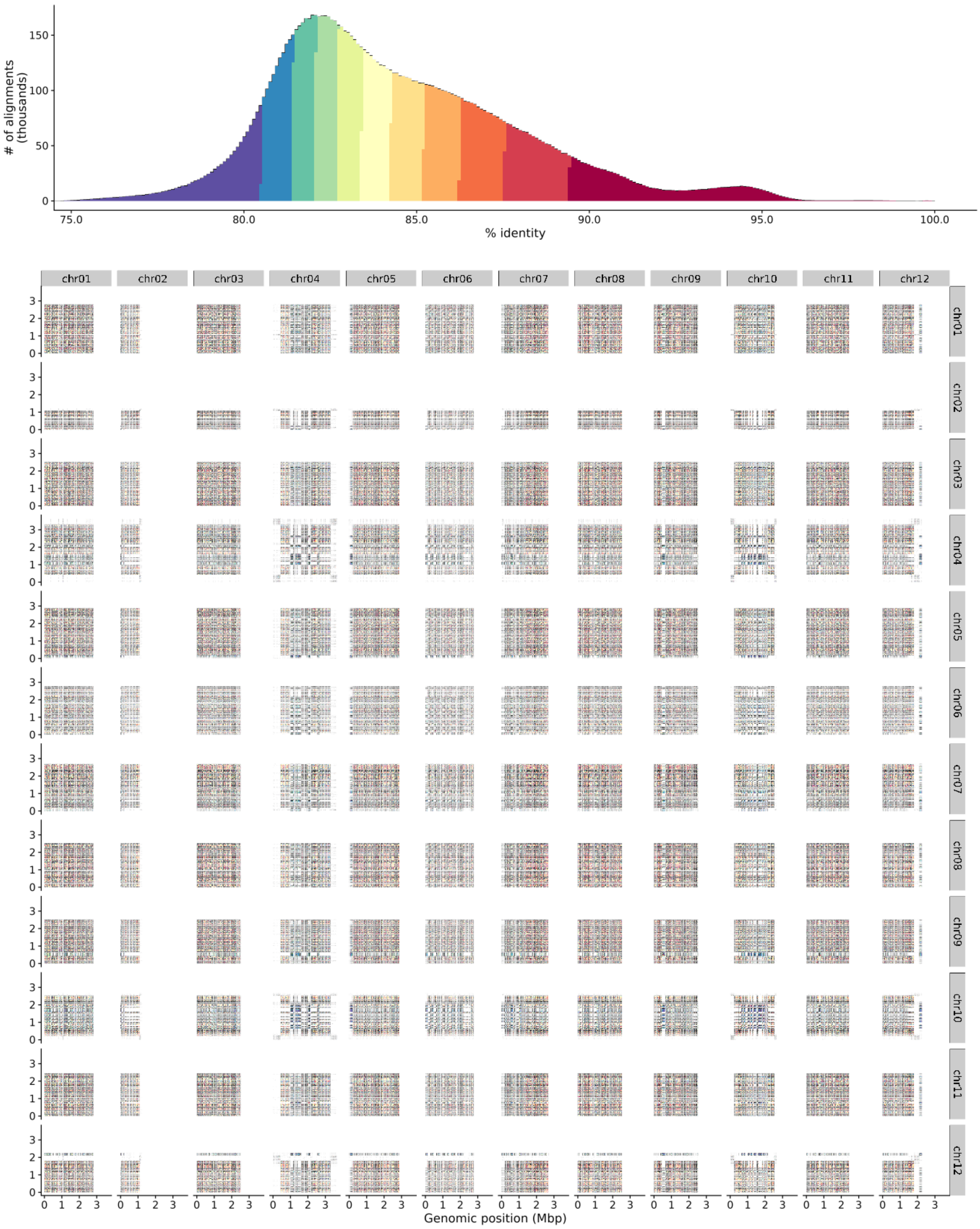


**Supplementary Figure 6. Heatmaps showing the sequence identities across centromeres.**

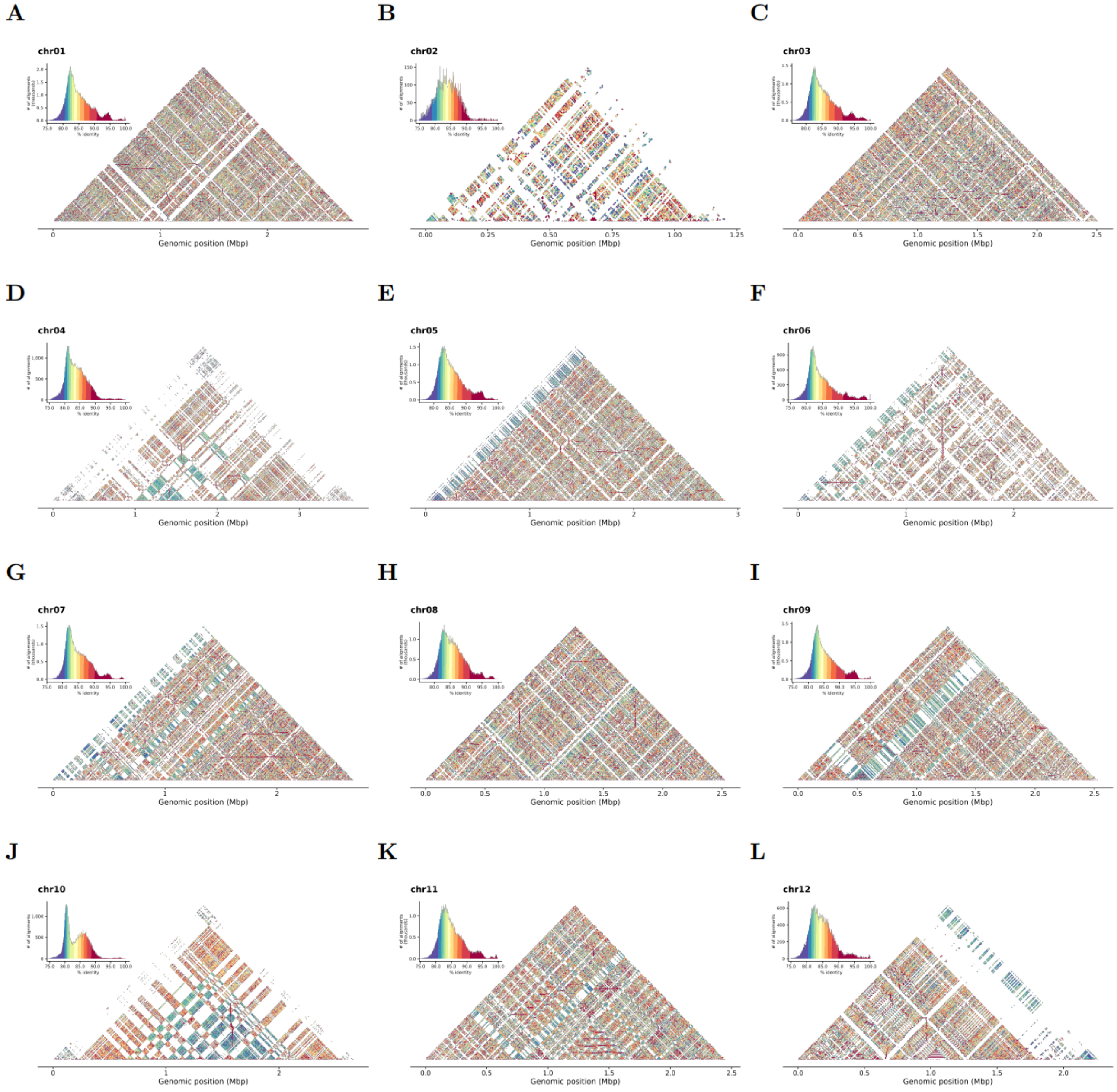


**Supplementary Figure 7. Heatmaps showing the sequence identities within each centromere.**

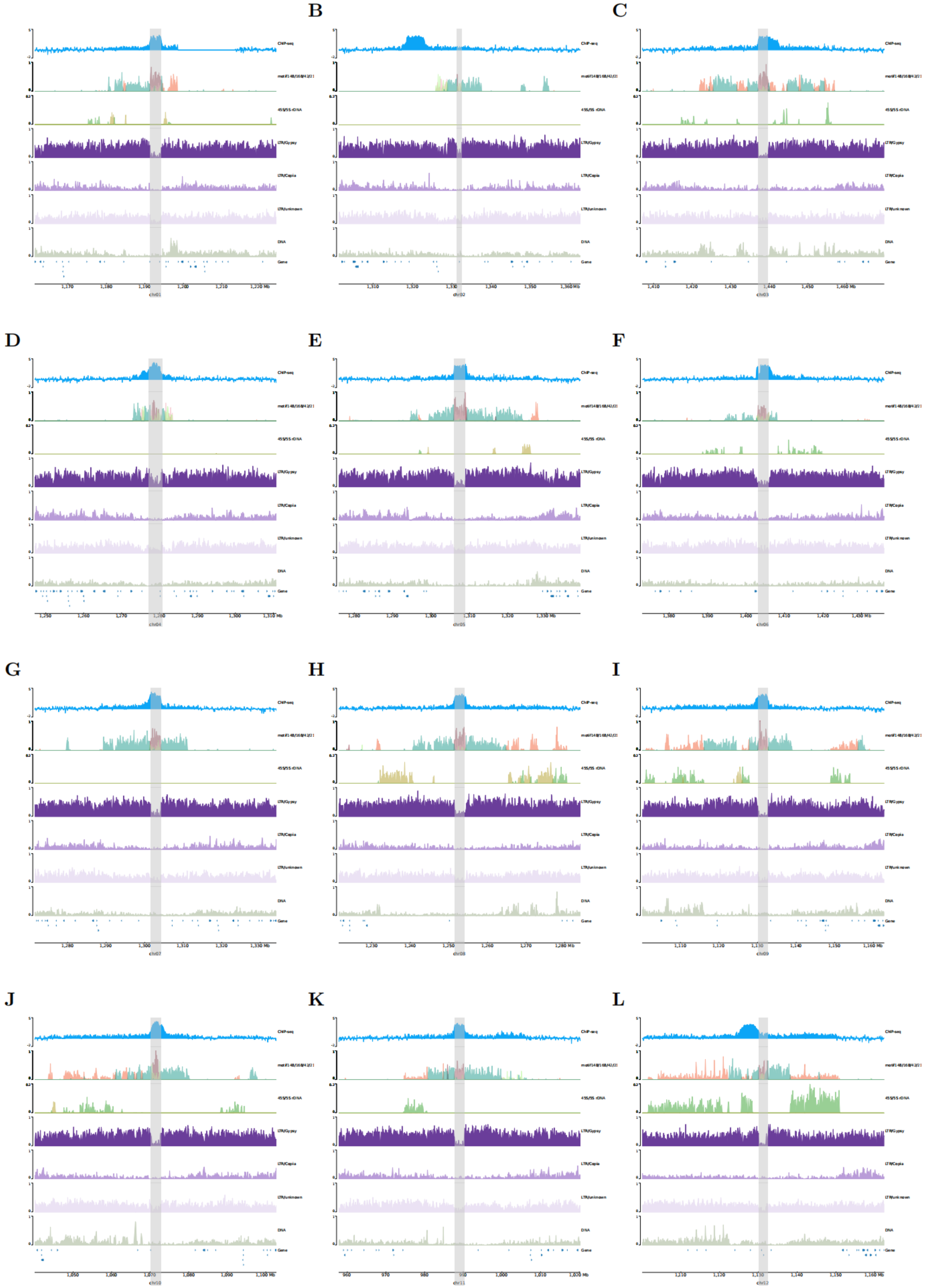


**Supplementary Figure 8. Characteristics of each centromere, including CENH3 ChIP-seq, tandem repeats, TEs and rDNAs.**

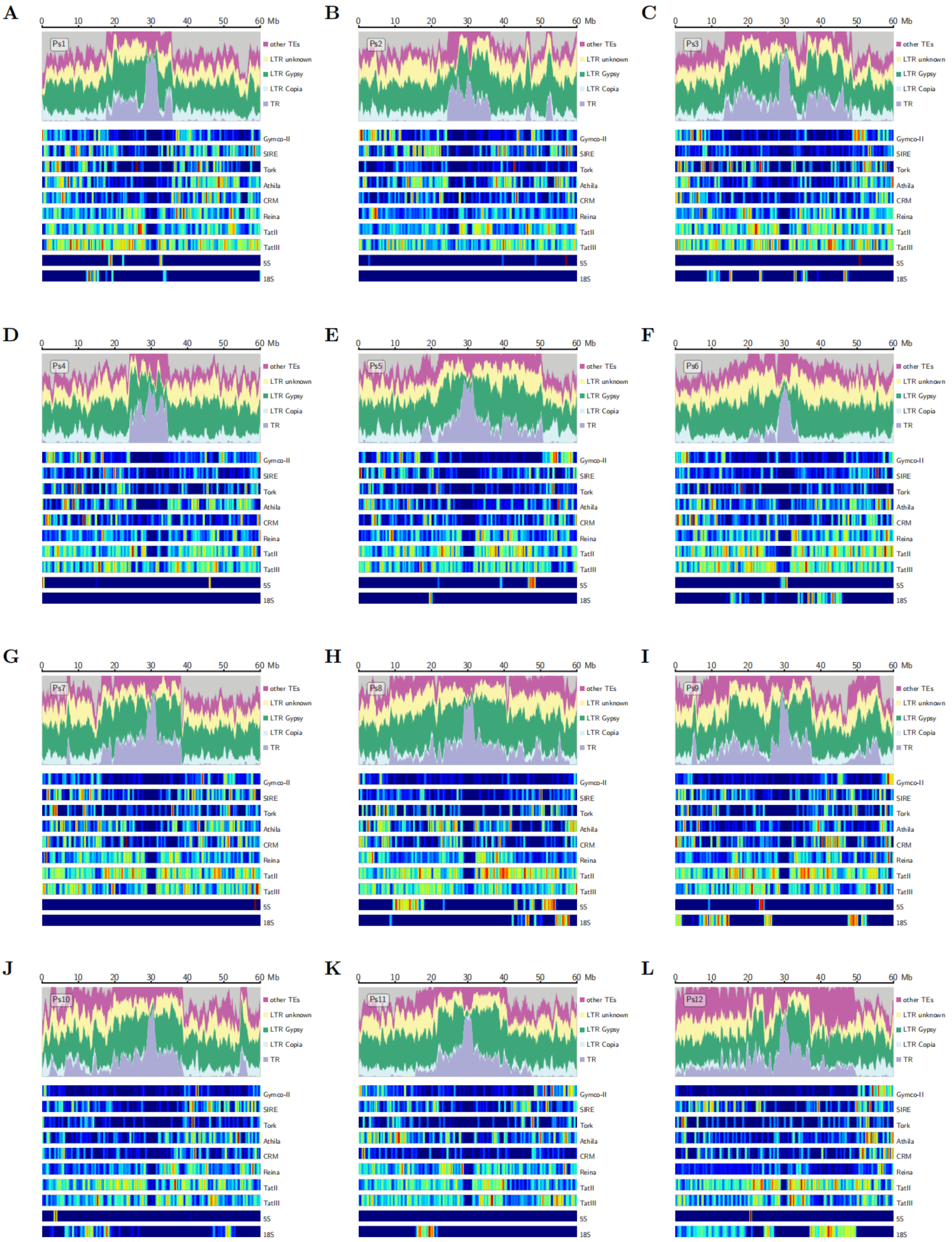


**Supplementary Figure 9. Heatmaps showing LTR-RT families of each centromere.**

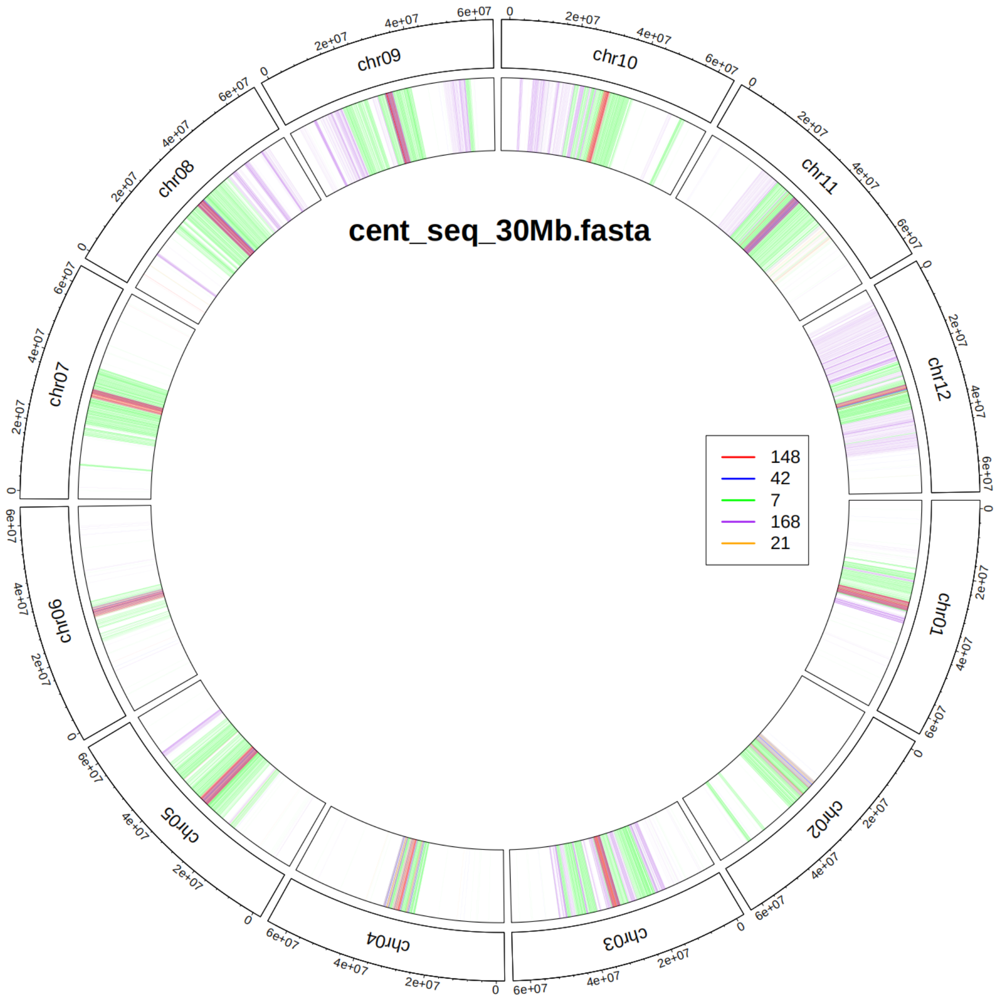


**Supplementary Figure 10. The tandem repeats in the peri-centromeric regions.** The colored lines indicate the tandem repeats with different motif lengths.

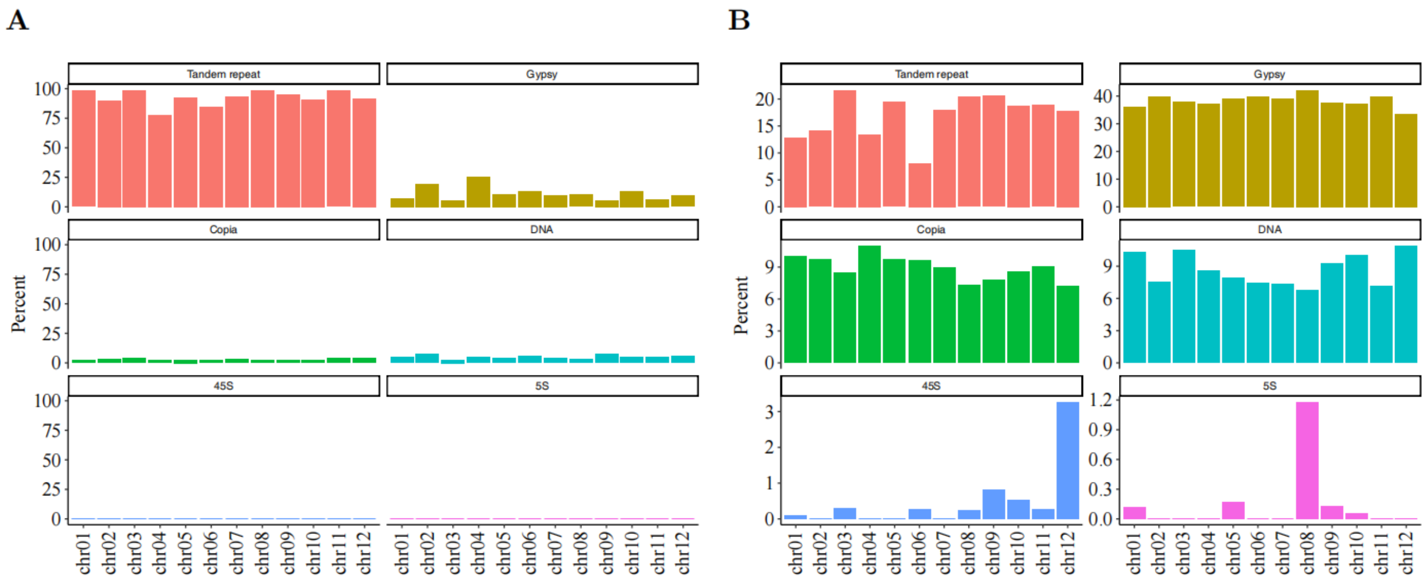


**Supplementary Figure 11. The composition of the centromeric (A) and peri-centromeric (B) regions.**

**A**

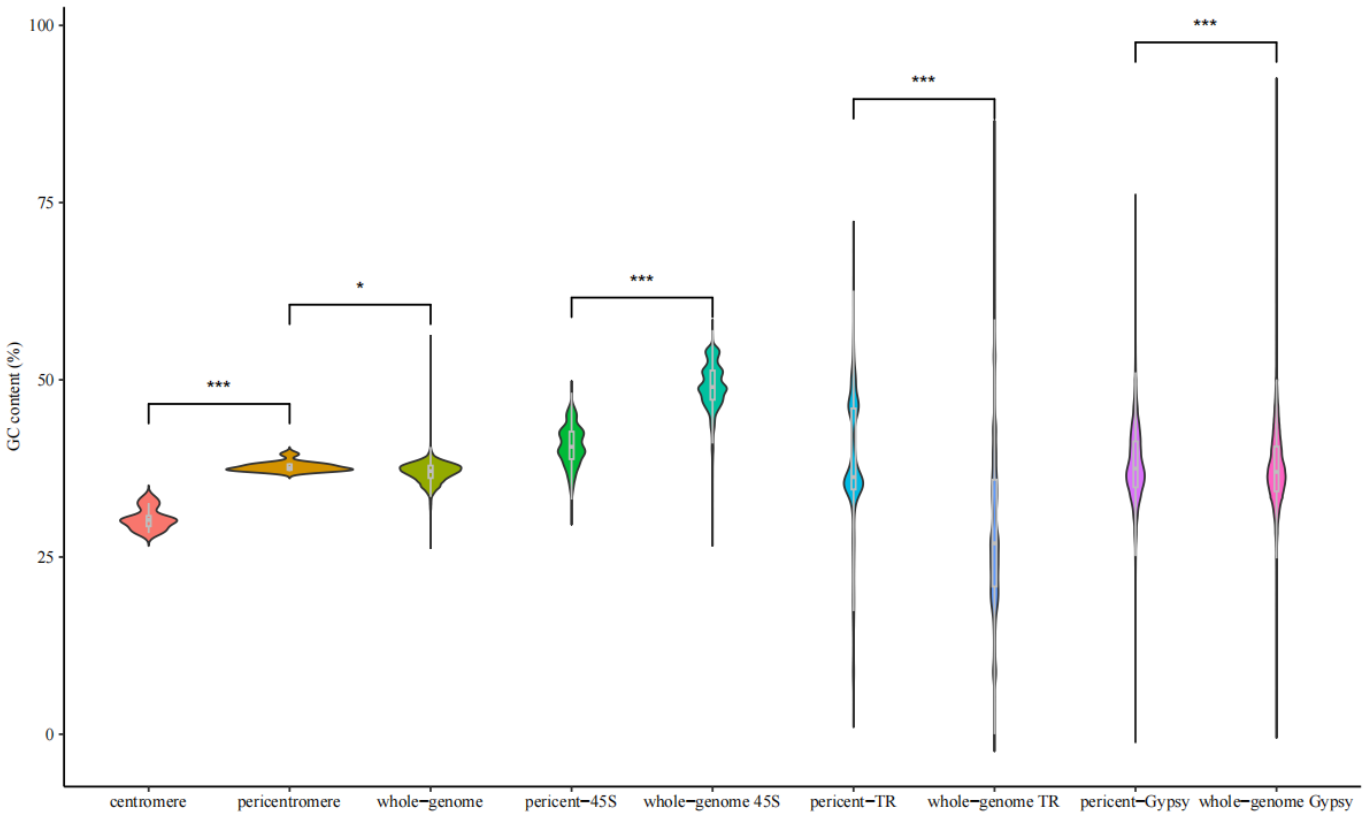


**B**

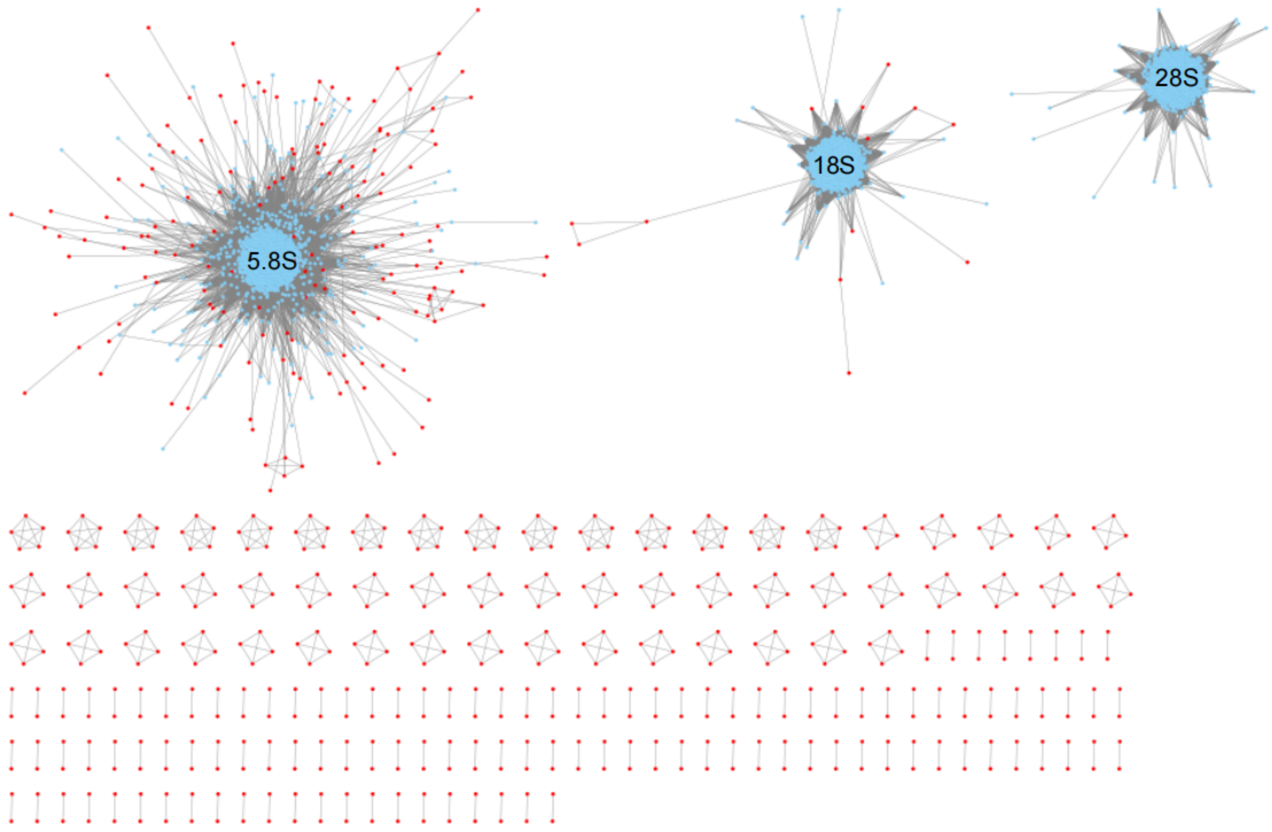


**Supplementary Figure 12. Degeneration of rDNAs within peri-centromeres. A)** GC content of different regions and elements within (peri-)centromeres and other regions (whole genome). **B)** Clustering of 45S rDNAs based on sequence identities. The red nodes indicate rDNAs within peri-centromeres, and the blue nodes indicate rDNAs outside peri-centromeres.

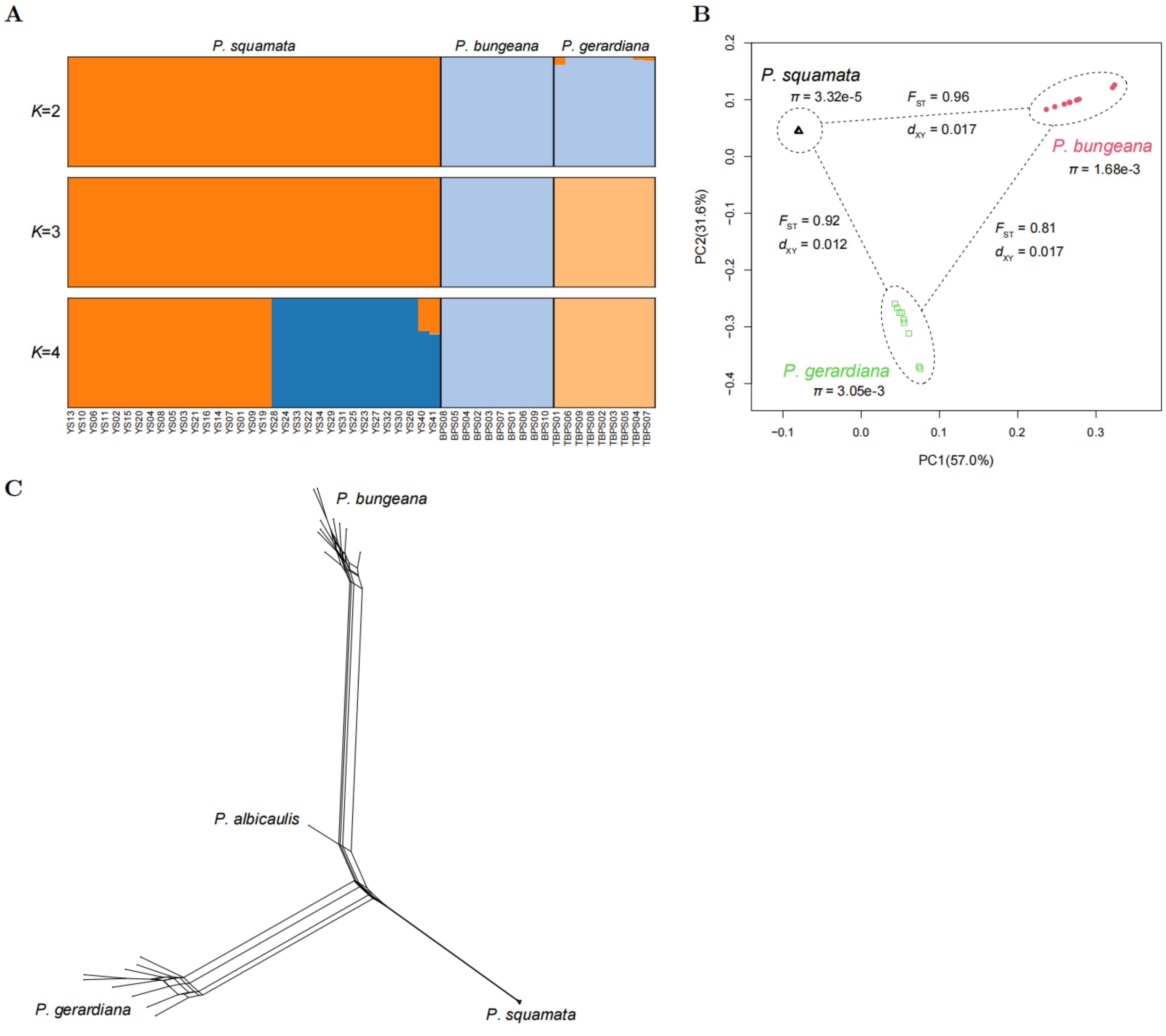


**Supplementary Figure 13. Population structure and differentiation among the three pines. A)** Population structure inferred by ADMIXTURE ($K = 2$ and $K = 3$) based on unlinked SNPs. **B)** Principal Component Analysis (PCA) of the first two principal components (PC1 and PC2). **C)** Phylogenetic network reconstructed using the Neighbor-Net algorithm in SplitsTree.

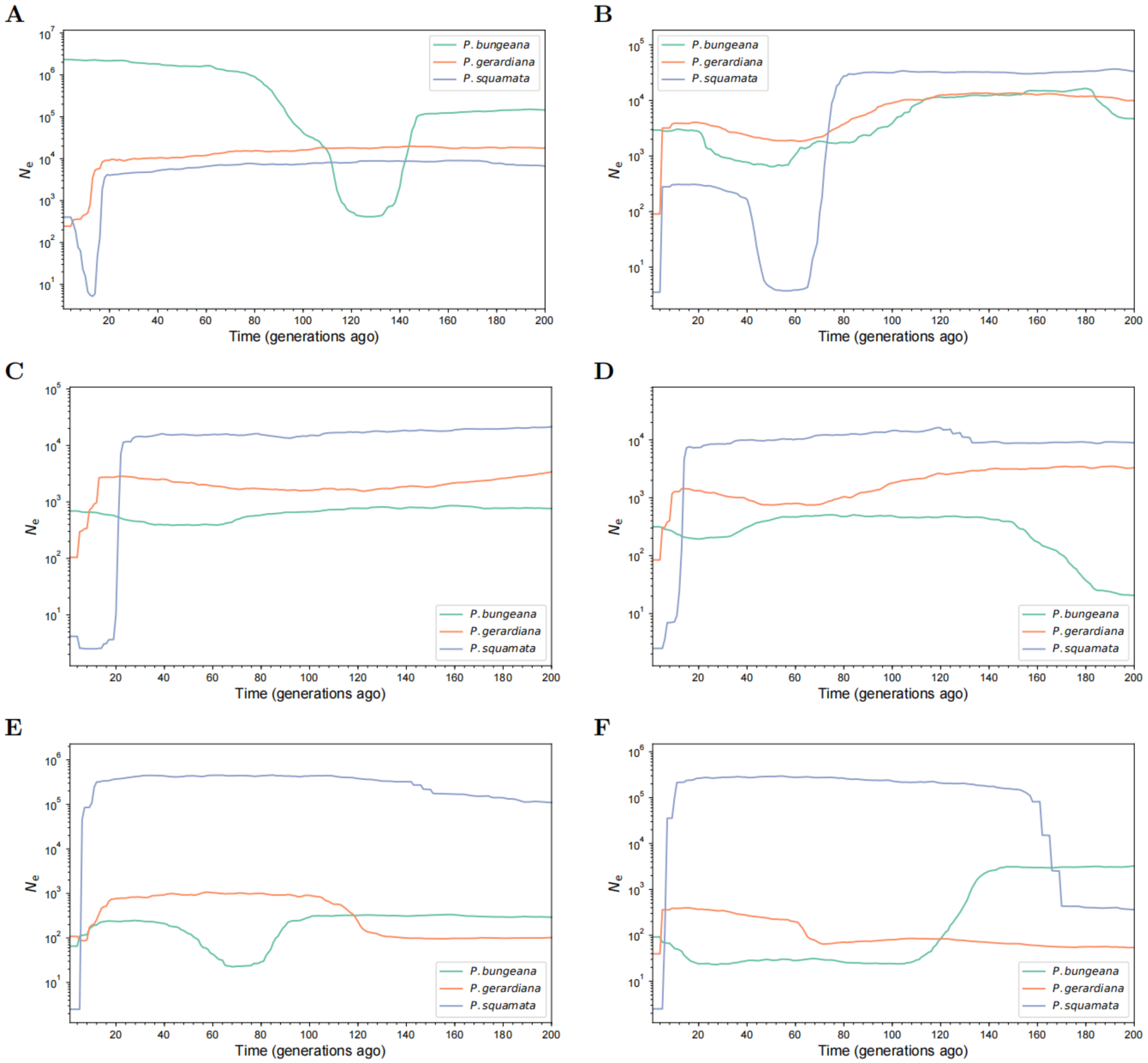


**Supplementary Figure 14. Recent demographic history estimated by GONE with different settings for recombination rate. A–F)** Recombination rate settings: 0.01 cM/Mb (**A**), 0.05 cM/Mb (**B**), 0.1 cM/Mb (**C**), 0.2 cM/Mb (**D**), 0.5 cM/Mb (**E**), and 1 cM/Mb (**F**). As the assumed recombination rate increases, the estimated onset of the bottleneck shifts to a more recent time, which is consistent with theoretical expectations.

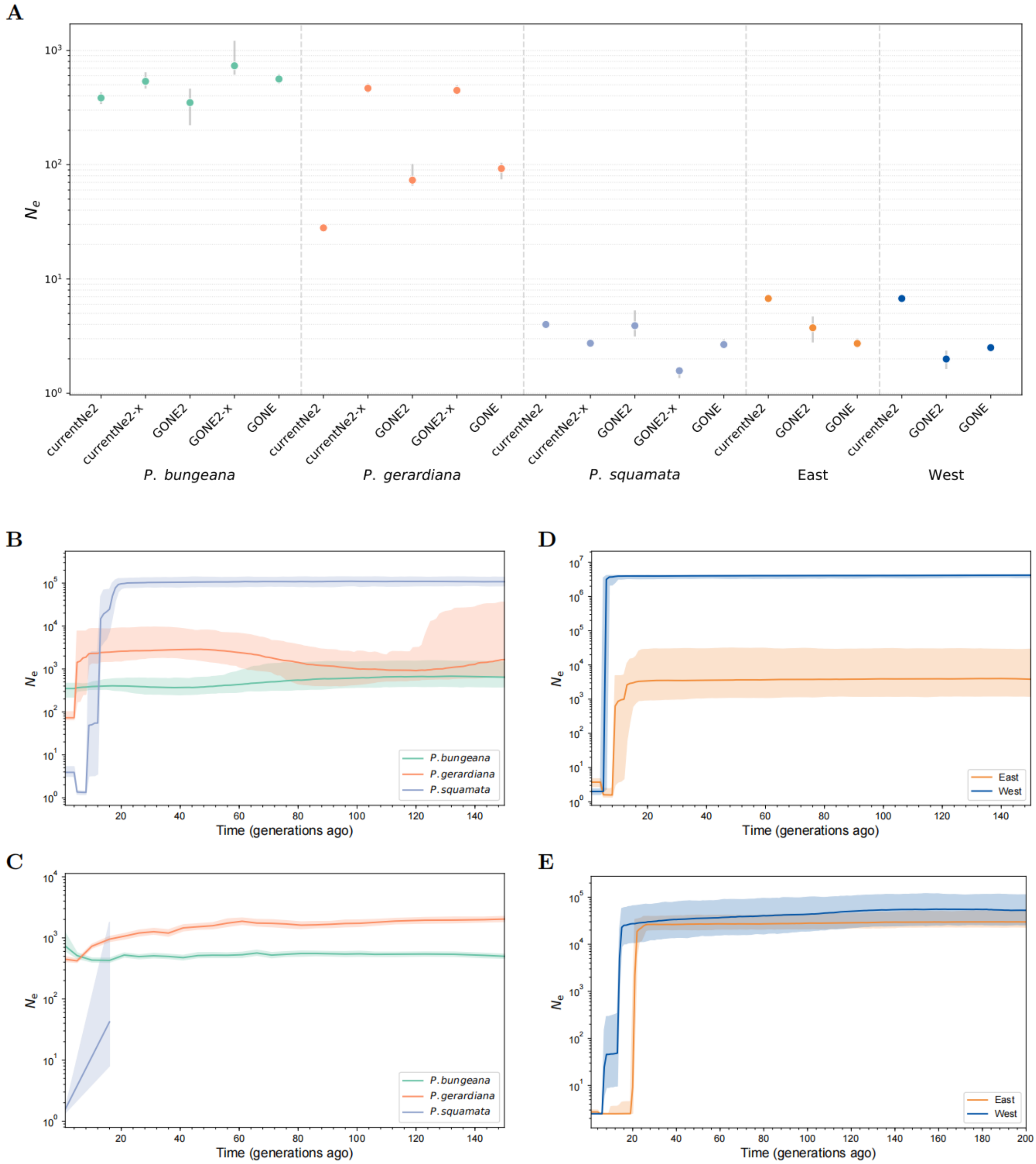


**Supplementary Figure 15. Contemporary $N_e$ and historical demographic dynamics inferred by LD-based methods. A)** Comparison of contemporary $N_e$ estimates derived from currentNe2, GONE2, and GONE. Dots represent median values, and vertical gray lines indicate the 95% confidence intervals (CI). **B–C)** Recent demographic trajectories inferred using GONE2, illustrating the impact of accounting for population structure without (B) and with (C) the -x option (-x is designed to account for population structure). Note that for the -x analysis (C), the demographic inference only traces back to 16 generations ago for *P. squamata*. **D–E)** Independent demographic reconstructions for the East and West populations performed using GONE2 (D) and GONE (E), respectively. In panels B–E, solid lines represent median estimates and shaded areas denote the 95% CI.

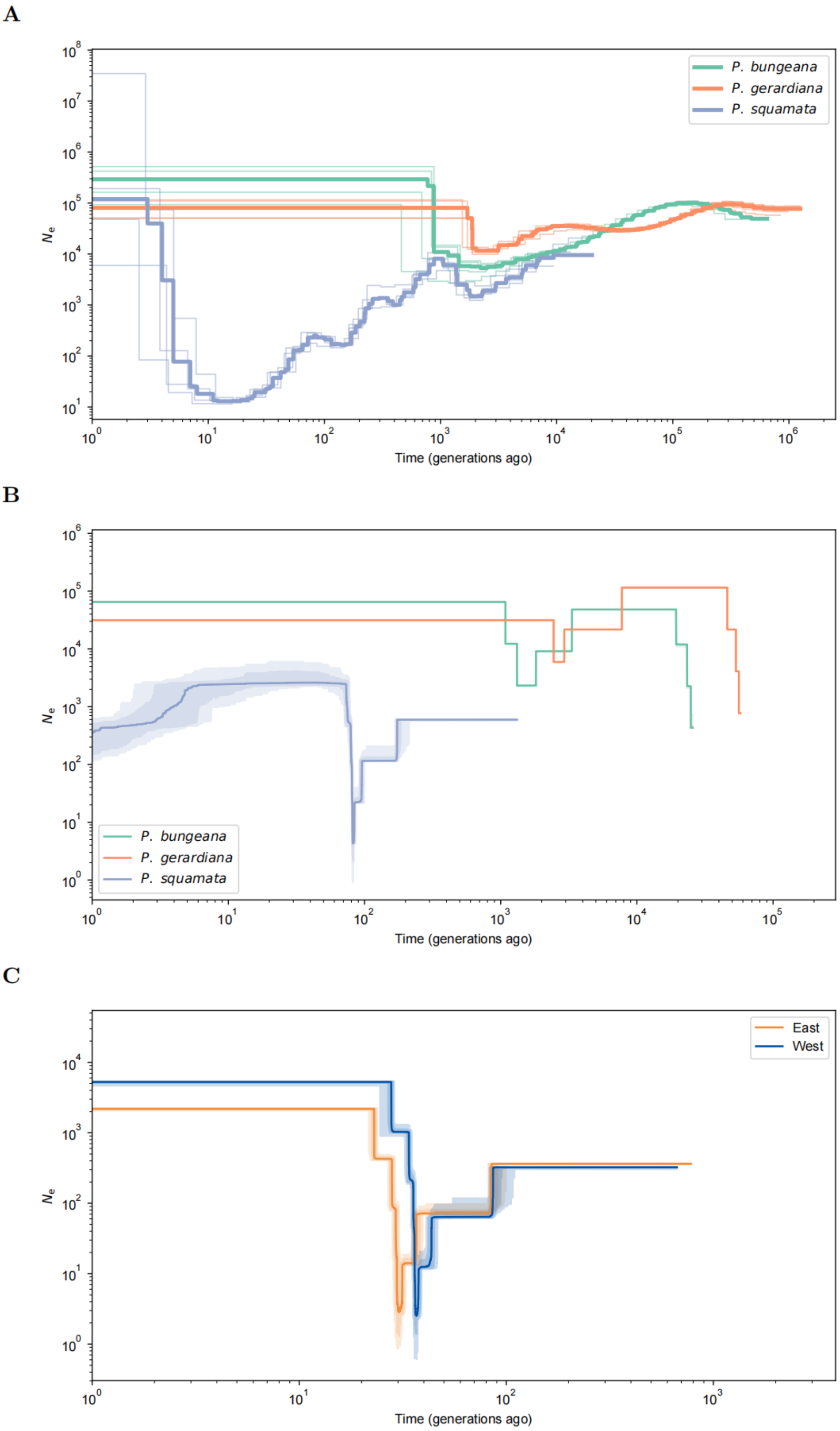


**Supplementary Figure 16. Demographic histories estimated by PSMC and Stairway Plot. A)** PSMC results for the three pine species. For each species, the four individuals with the highest sequencing depth were analyzed independently, yielding highly consistent trajectories. Estimates for *P. squamata* included two individuals each from the East and West populations, which also showed high consistency. Translucent thin lines represent individual estimates, while bold lines indicate the median across the four individuals per species. **B–C)** Stairway Plot results based on (B) species-level and (C) East vs. West population-level analyses. Solid lines represent the median estimates, with light and dark shaded areas denoting the 75% and 95% confidence intervals (CI), respectively.

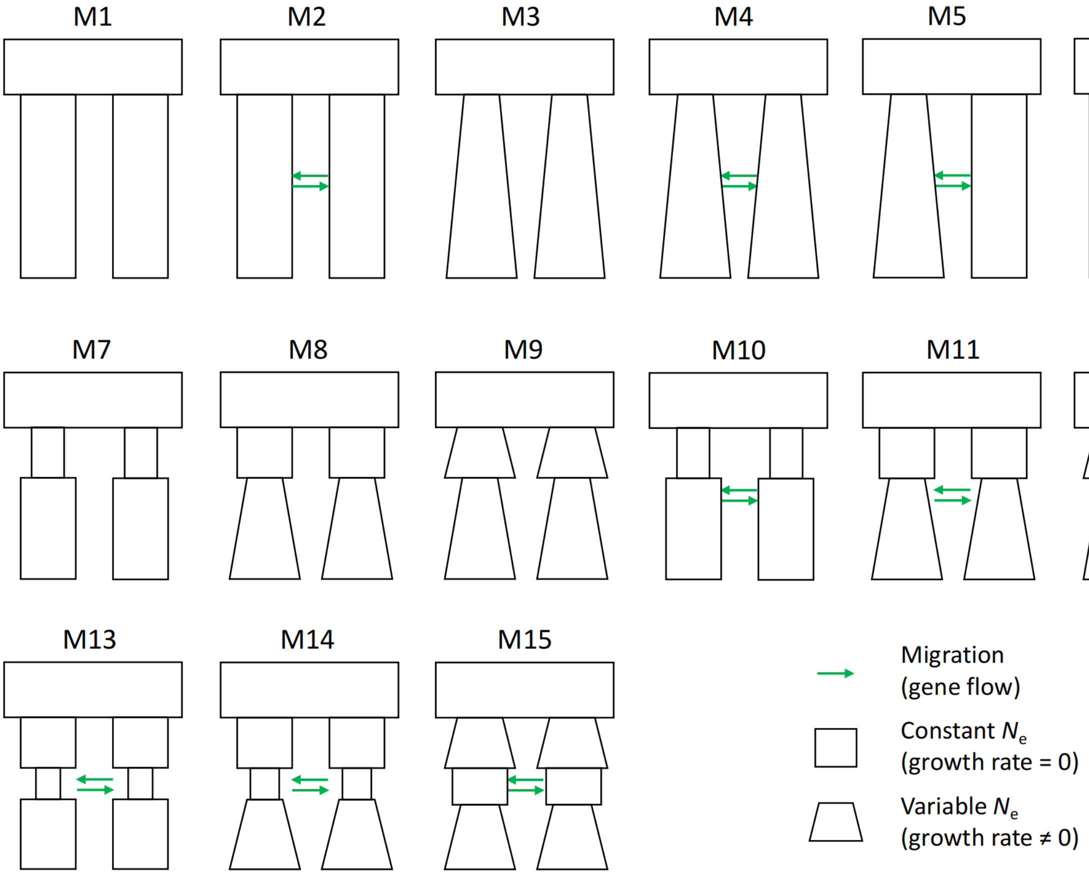


**Supplementary Figure 17. Demographic models tested using fastsimcoal2.** The tested models incorporate varying population growth rates (zero vs. non-zero) and migration rates (zero vs. $> 0$). Models were structured into one to three distinct demographic epochs after split. Migration rates were treated as global parameters, meaning the same migration rate was shared across all demographic time periods within a given model.

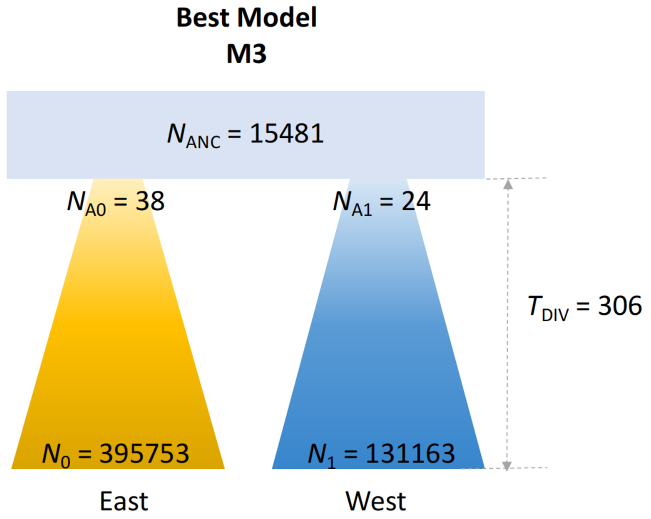


**Supplementary Figure 18. The best-fit demographic model inferred by fastsimcoal2.** The optimal model reveals an extremely low effective population size ($N_e$) following the population split, representing a severe bottleneck event, followed by a dramatic population expansion in both populations. No migration was detected between the two populations in this best-fit scenario.

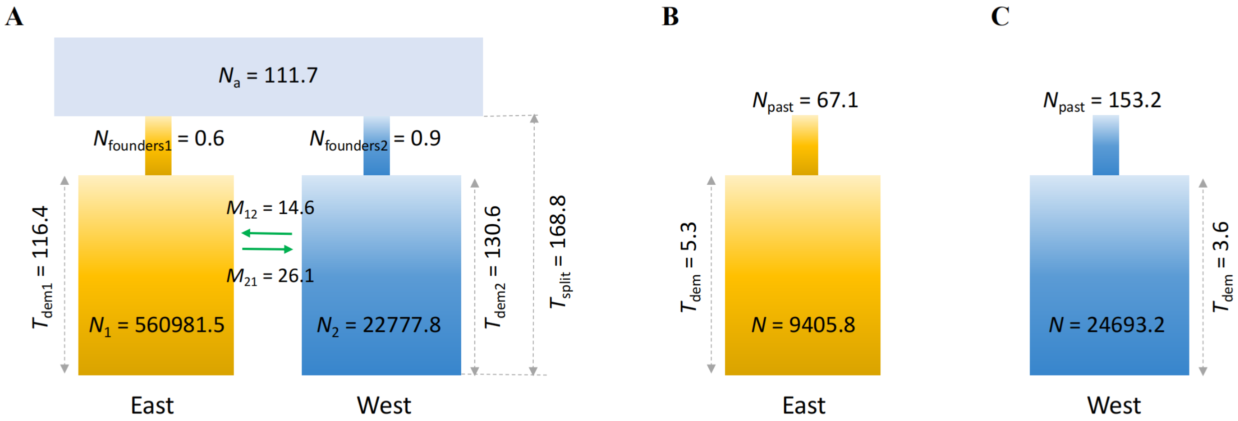


**Supplementary Figure 19. The best-fit demographic model inferred by DILS. A)** The Isolation-with-Migration (IM) model. This model illustrates a severe bottleneck following the initial isolation, with the effective population size ($N_e$) of the founding individuals for both populations being less than 1, followed by a dramatic population expansion. Ongoing gene flow was detected between the two populations, with the migration rate denoted as $M = 4N_e m$ (where $m$ represents the migration rate per generation). **B–C)** Independent demographic histories estimated for the East (B) and West (C) populations, respectively. Both analyses similarly reveal a severe bottleneck followed by a rapid population expansion within the last few generations.

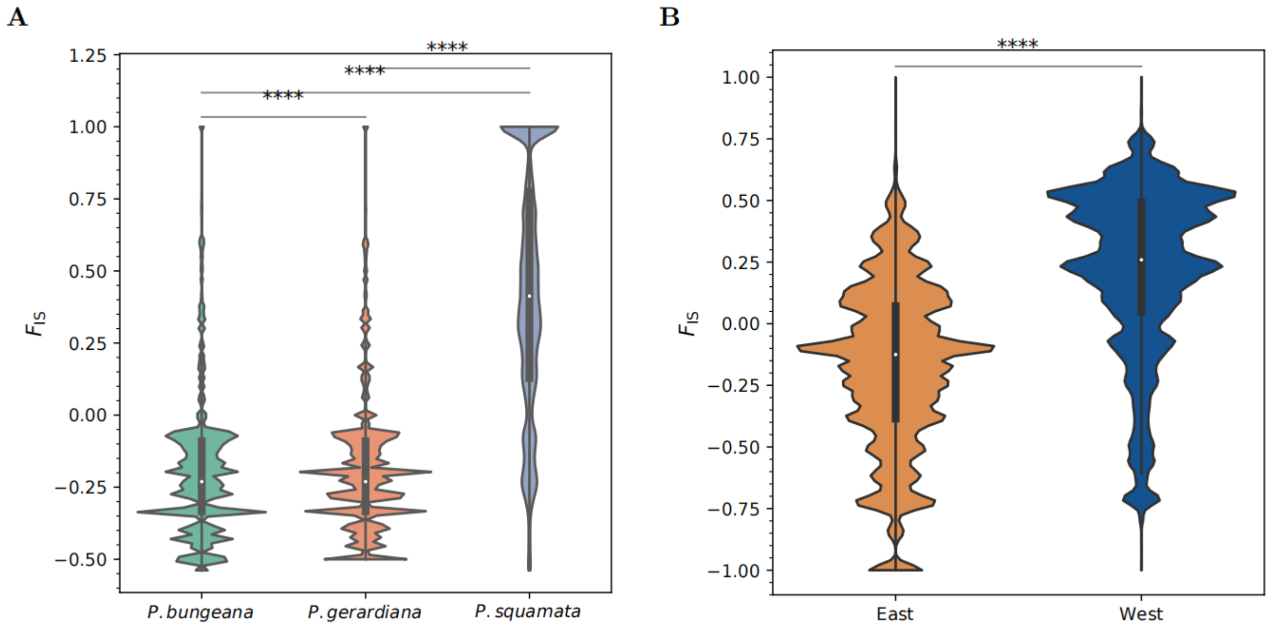


**Supplementary Figure 20. Estimates of the inbreeding coefficient ($F_{IS}$) based on SNPs (MAF > 0.05). A)** $F_{IS}$ values estimated at the species level. **B)** $F_{IS}$ values estimated separately for the East and West populations to account for and mitigate the potential confounding effects of population structure.

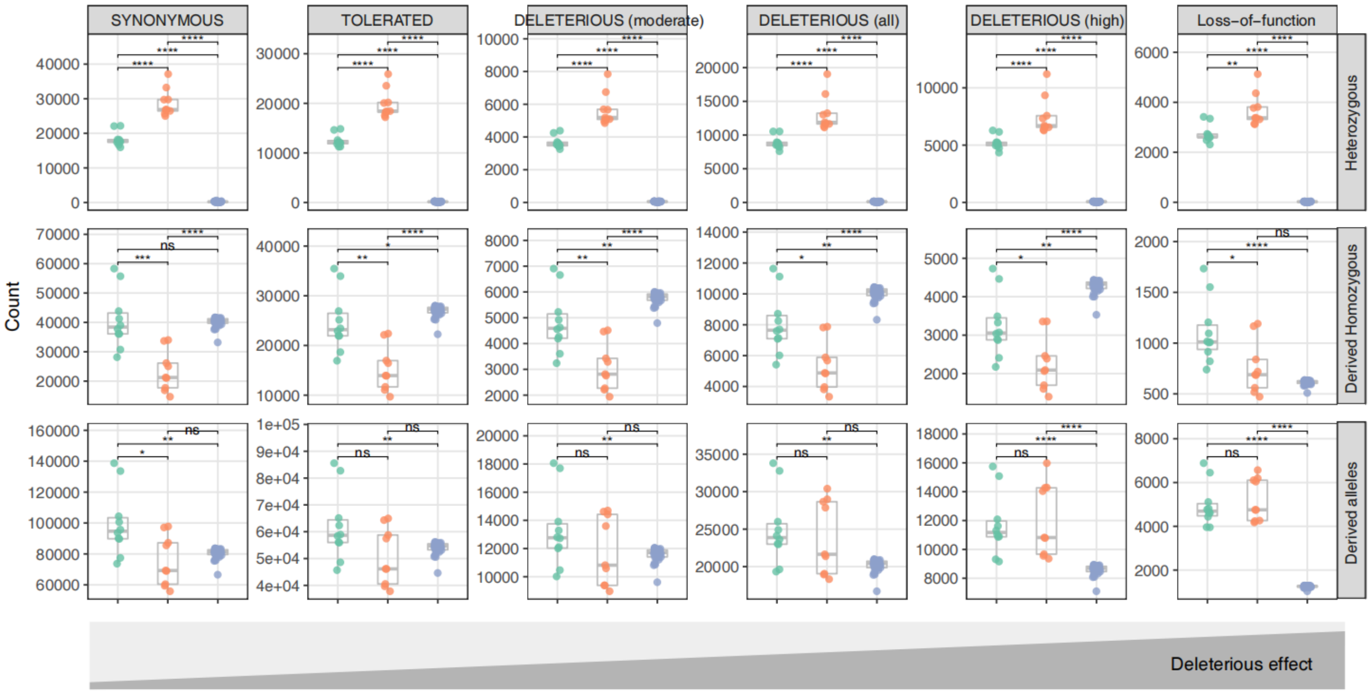


**Supplementary Figure 21. Count of synonymous and non-synonymous mutations with different deleterious effects in the three pines.** Statistical test: *, $P < 0.05$; ****, $P < 1e-4$, Wilcoxon rank-sum test.

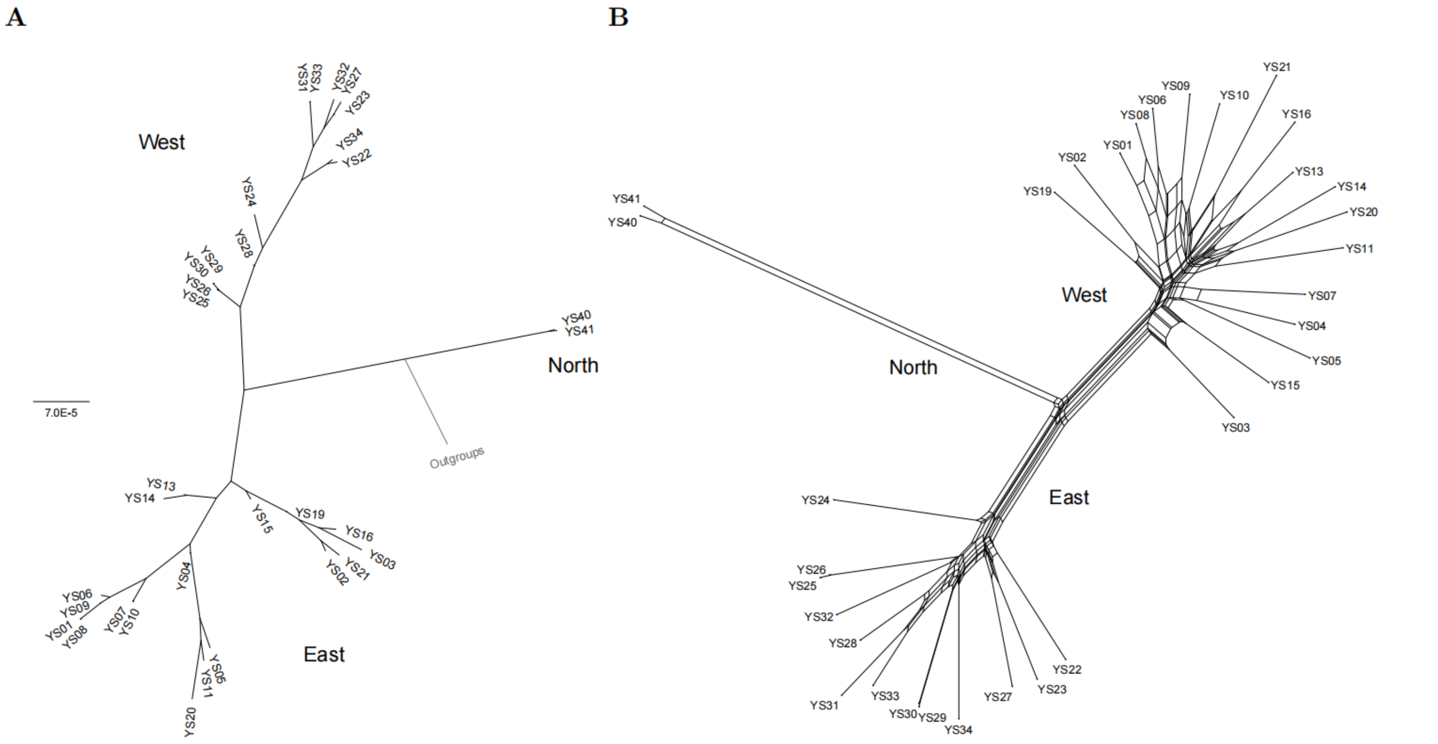


**Supplementary Figure 22. Phylogenetic analysis for *P. squamata*. A)** Maximum Likelihood (ML) tree, showing the phylogenetic relationships and hierarchical divergence among populations. **B)** Split network analysis, used to identify reticulate evolutionary signals or potential gene flow among populations.

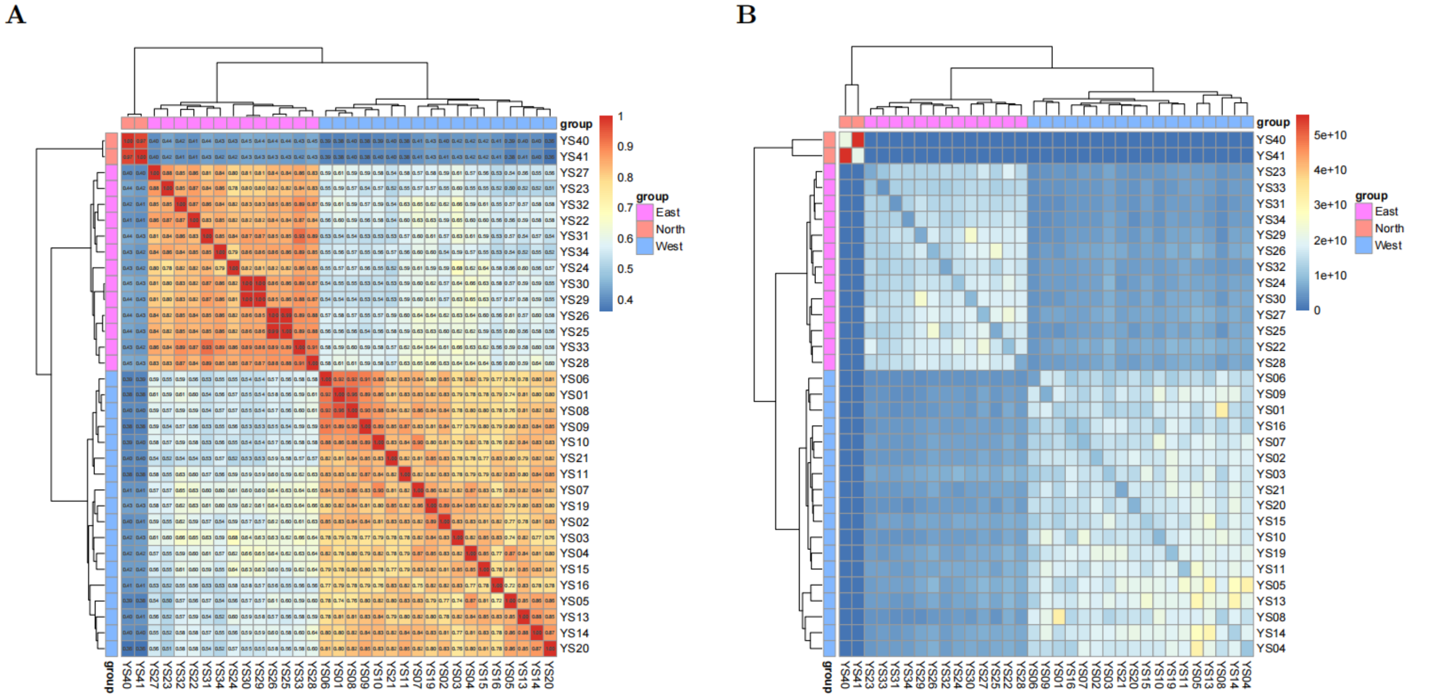


**Supplementary Figure 23. IBS (A) and IBD (B) clustering for *P. squamata*.**

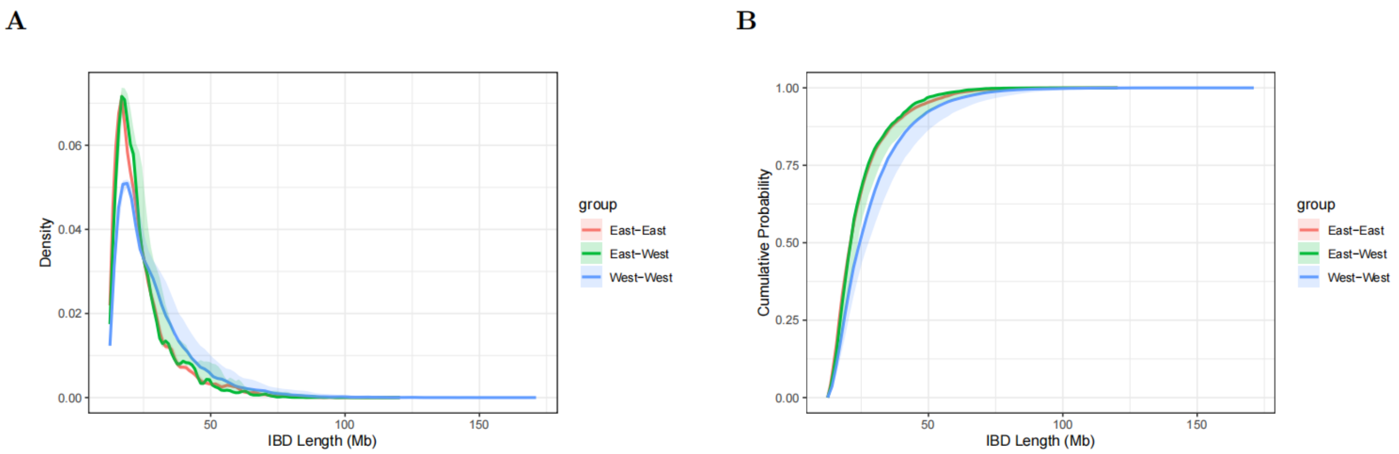


**Supplementary Figure 24. Distribution of IBD segment lengths. A)** Probability density distribution of IBD lengths. **B)** Cumulative density distribution of IBD lengths.

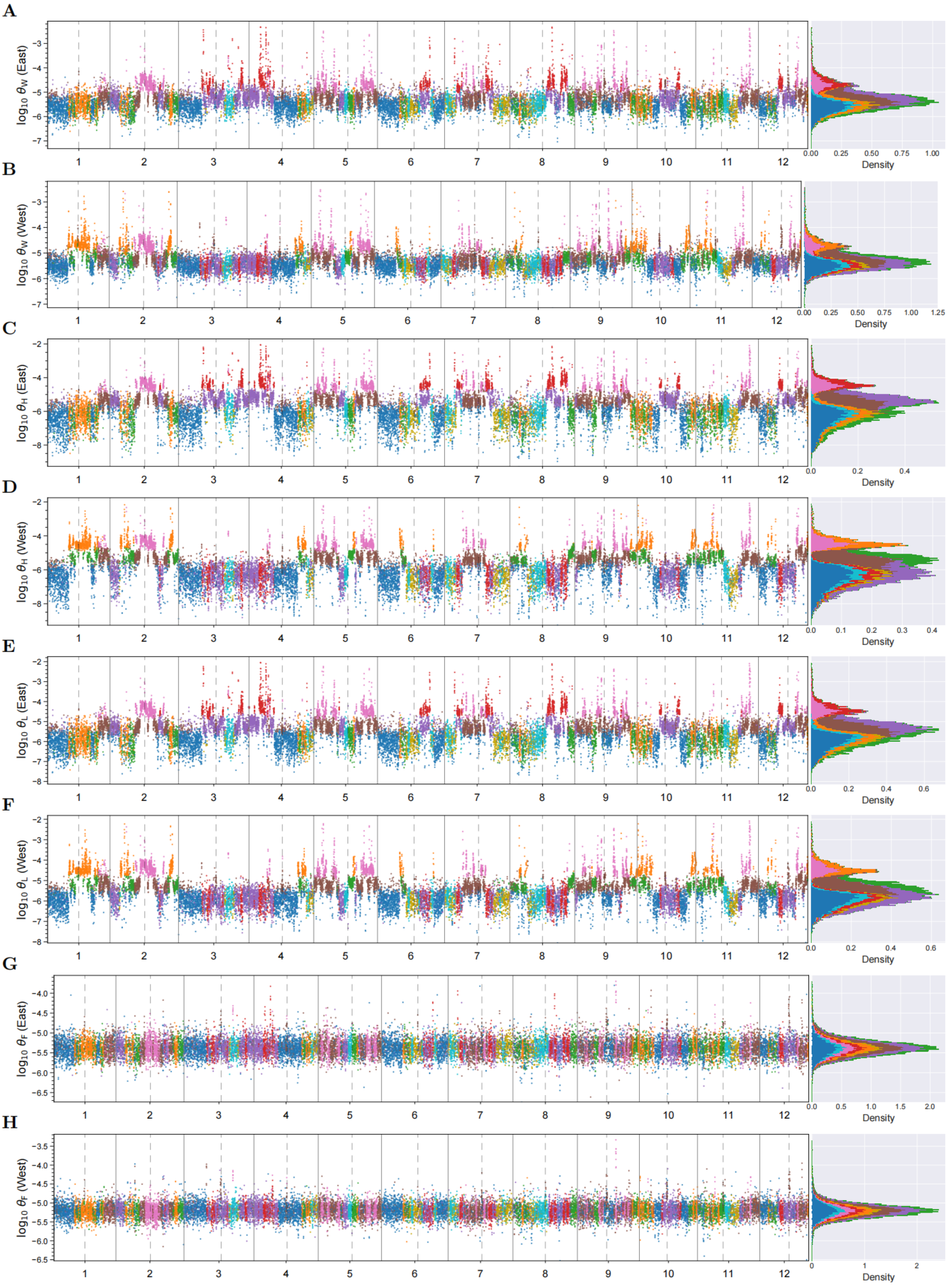


**Supplementary Figure 25. Manhattan plots of multiple estimations of $\theta$ by ANGSD. A–B)** Watterson's $\theta$ ($\theta_W$), where all mutations are weighted equally. **C–D)** $\theta_H$, which gives higher weight to high-frequency derived mutations. **E–F)** $\theta_L$, which also assigns higher weight to high-frequency derived mutations. **G–H)** $\theta_F$, representing the frequency of derived singletons.

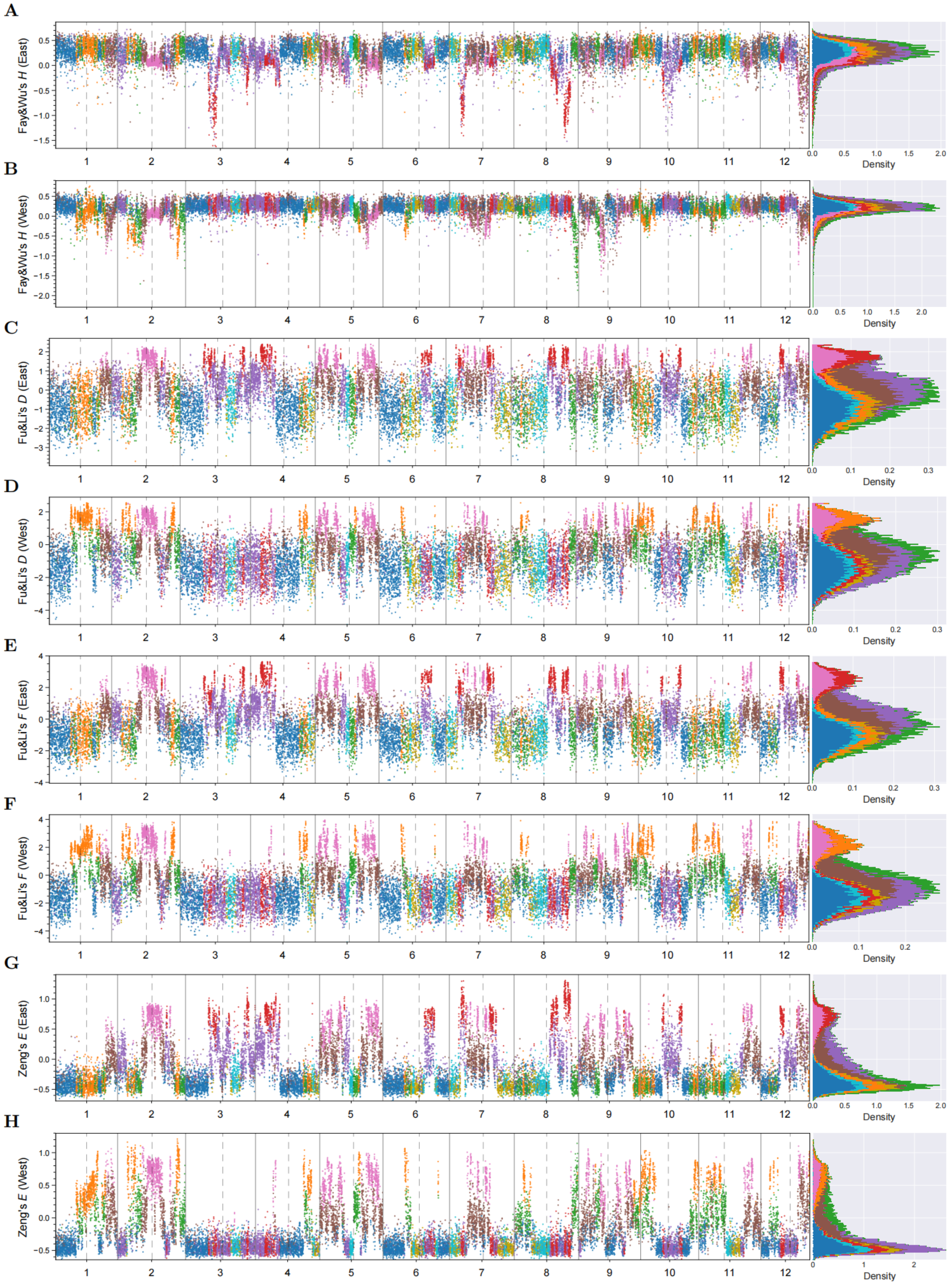


**Supplementary Figure 26. Manhattan plots of multiple neutrality tests by ANGSD. A–B)** Fay and Wu's $H$ (standardized $\theta_\pi - \theta_\mathrm{H}$), comparing intermediate- vs. high-frequency variants. **C–D)** Fu and Li's $D$ (standardized $\theta_\pi - \theta_\mathrm{F}$), comparing intermediate- vs. extremely low-frequency variants. **E–F)** Fu and Li's $F$ (standardized $\theta_\mathrm{W} - \theta_\mathrm{F}$), comparing low vs. extremely low-frequency variants. **G–H)** Zeng's $E$ (standardized $\theta_\mathrm{L} - \theta_\mathrm{W}$), comparing high- vs. low-frequency variants.

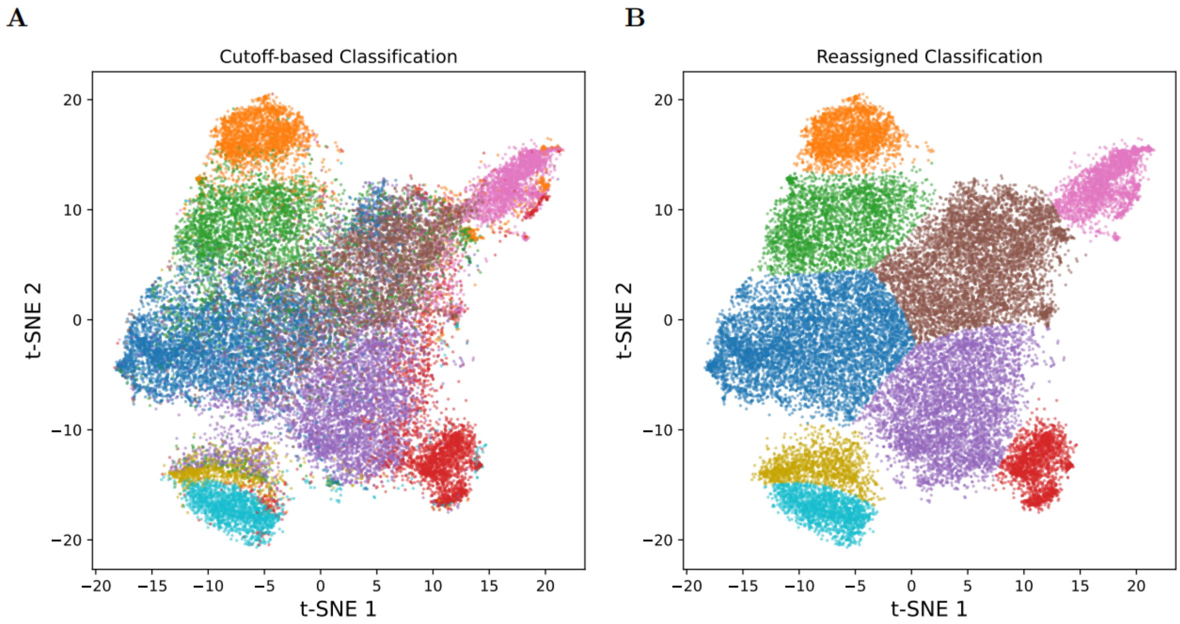


**Supplementary Figure 27. Classification of window-based genomic clusters. A)** Classification based on hard cutoffs on summary statistics. **B)** Classification after manual adjustment and SVC re-classification.

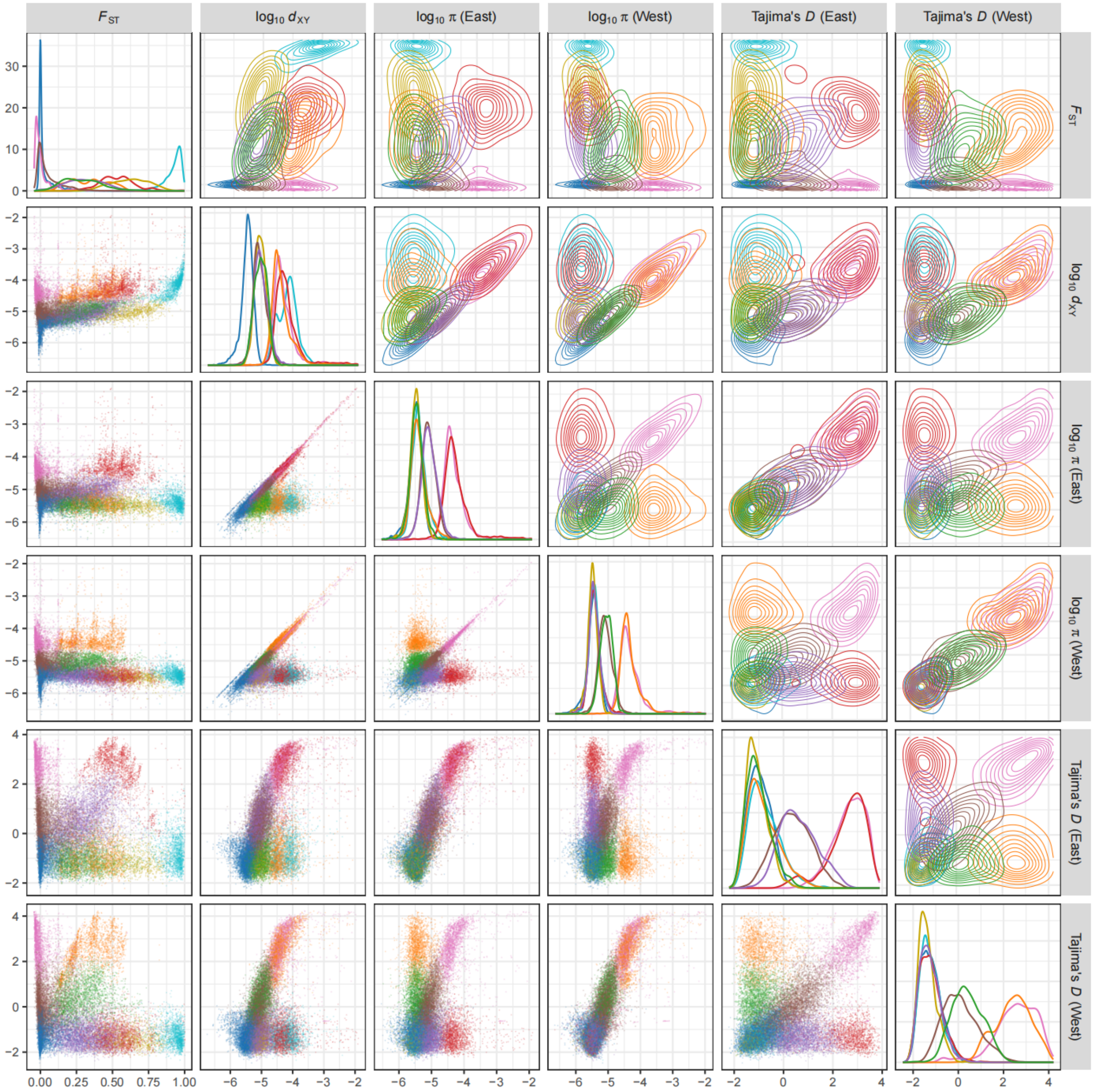


**Supplementary Figure 28. 2D distribution of $F_{ST}$ and $d_{XY}$ between East and West populations, and $\pi$ and Tajima's $D$ of East population and West population.**

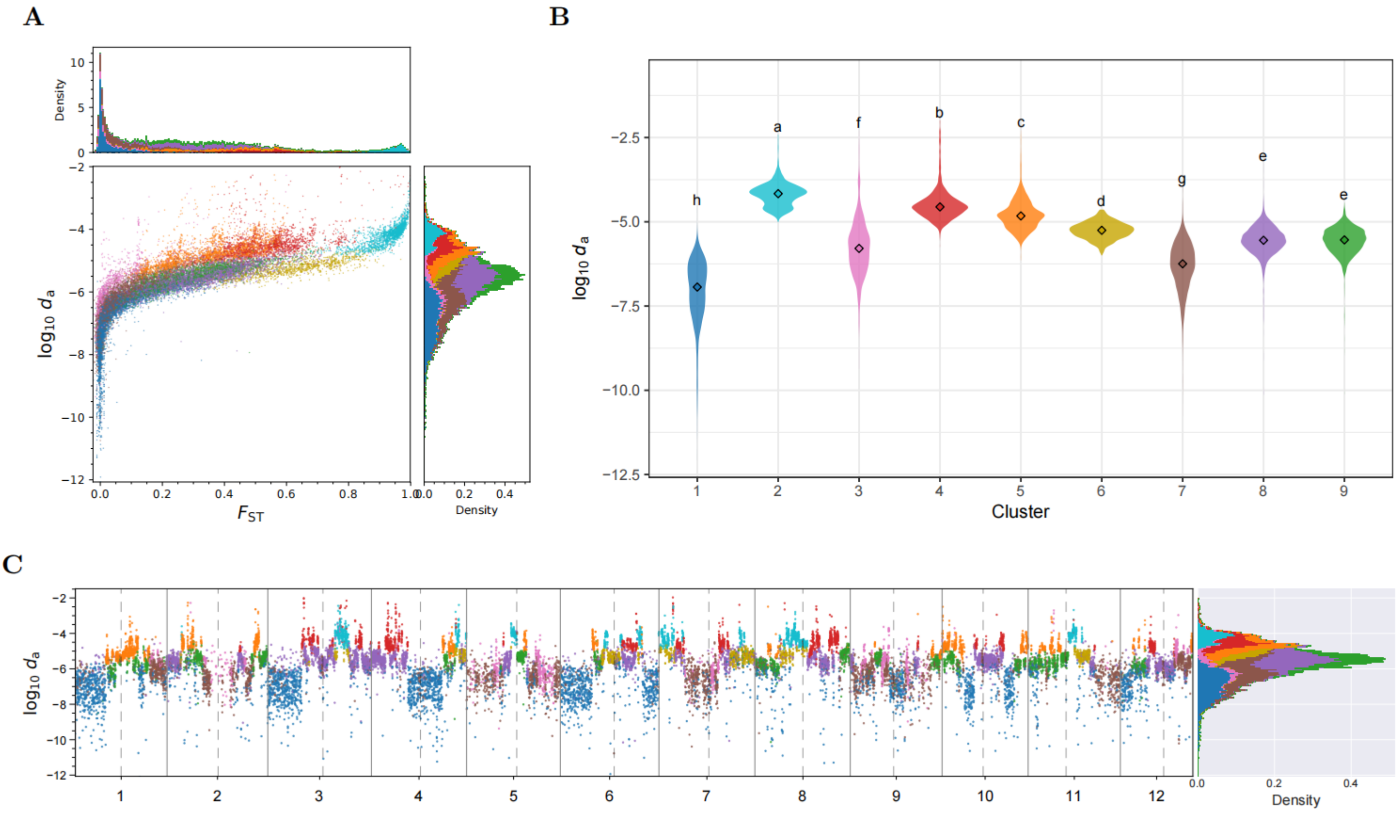


**Supplementary Figure 29. Distribution of net nucleotide differences ($d_a$).** **A)** 2D distribution of $d_a$ and $F_{ST}$. **B)** $d_a$ distribution across nine clusters. **C)** Genome-wide distribution of $d_a$. All panels are color-coded by cluster.

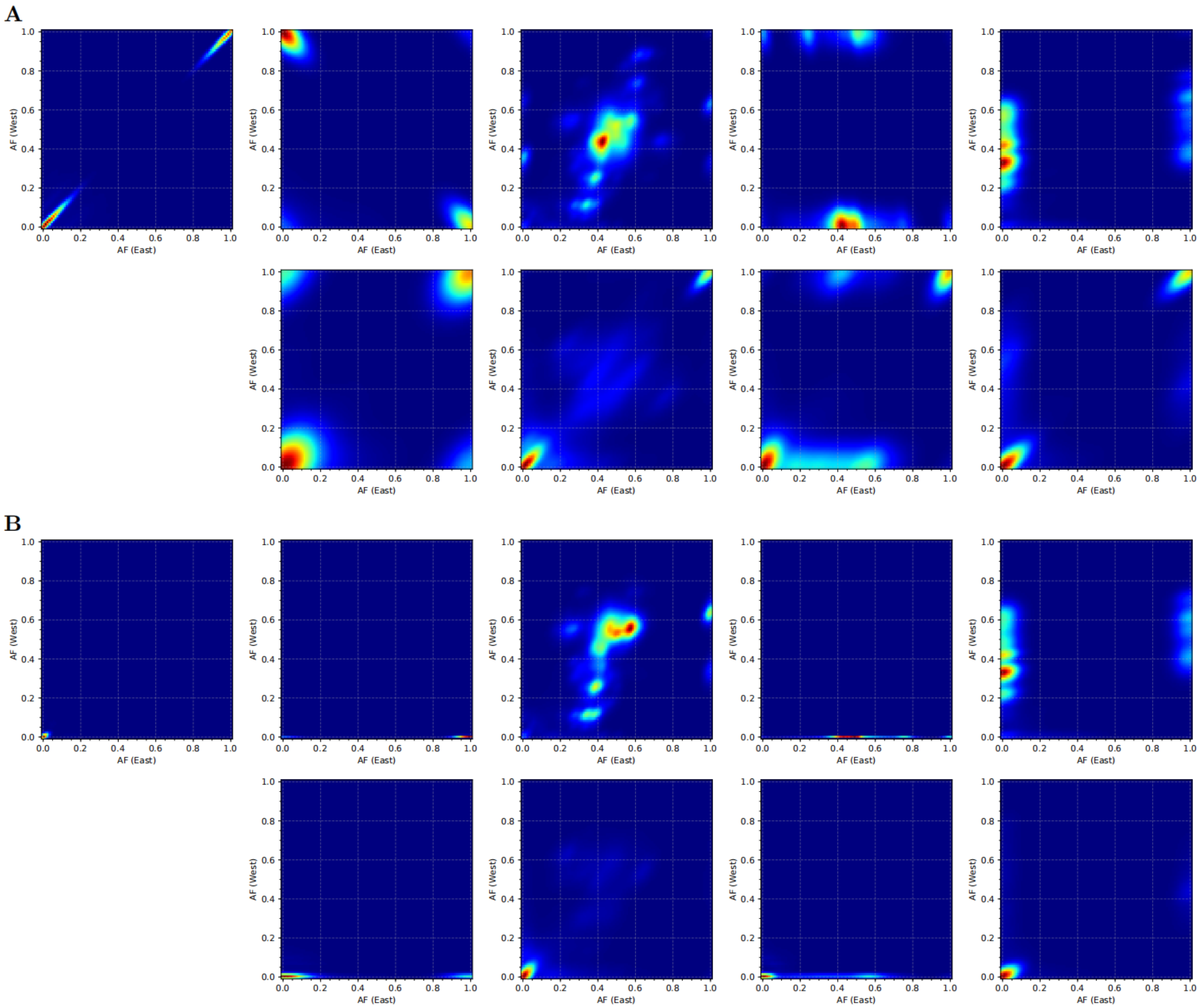


**Supplementary Figure 30. Allele frequency spectrum (AFS) between East and West populations for the nine clusters.** **A)** Based on polarized SNPs. **B)** Based on unpolarized SNPs. For both panels, clusters 1–5 are arranged from left to right in the top row, and clusters 6–9 in the bottom row.

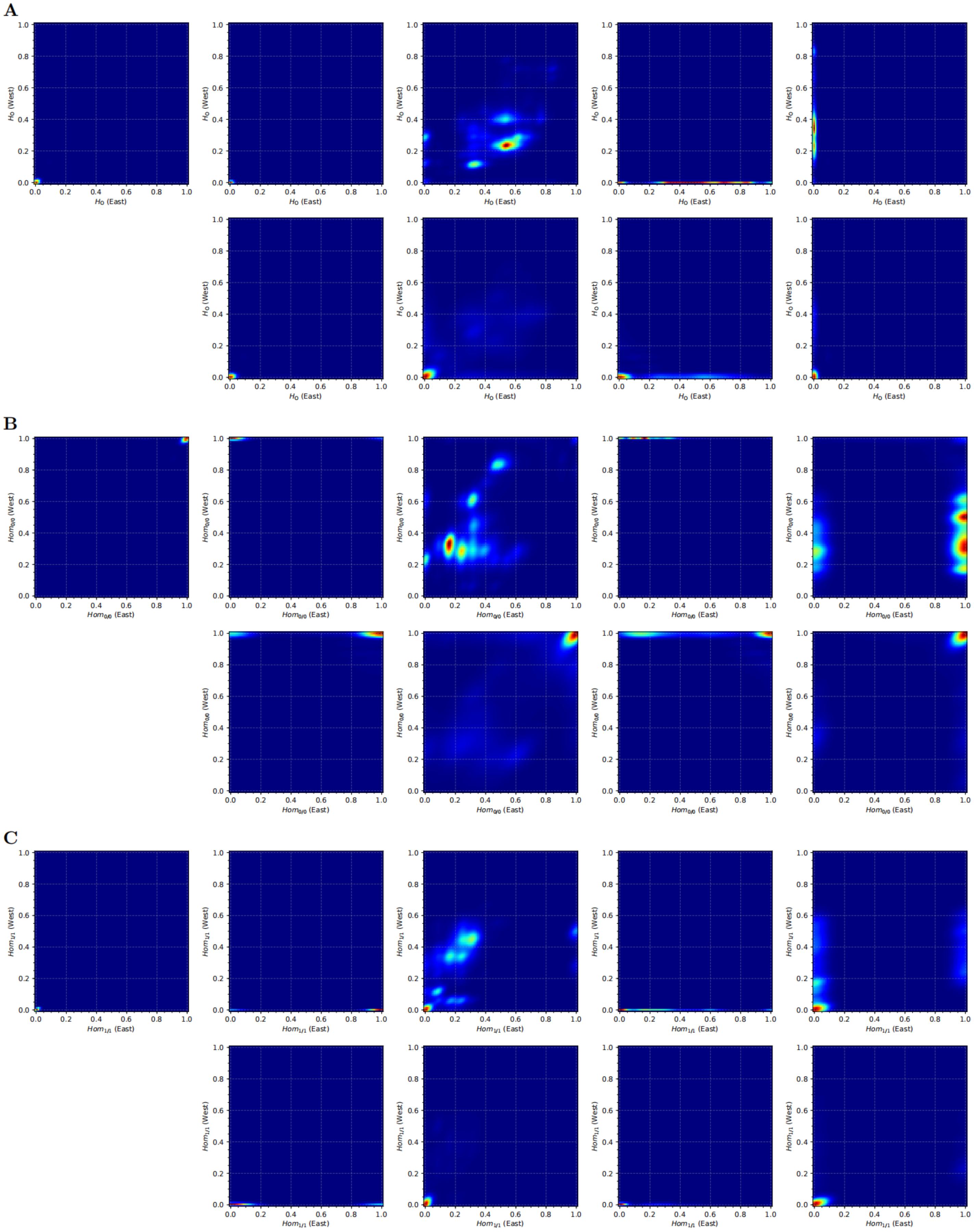


**Supplementary Figure 31. Heterozygosity and Homozygosity frequency spectrum between East and West populations for the nine clusters. A)** Observed heterozygosity ($H_O$). **B)** One type of homozygosity (genotype 0/0). **C)** The other type of homozygosity (genotype 1/1). For all panels, clusters 1–5 are arranged from left to right in the top row, and clusters 6–9 in the bottom row.

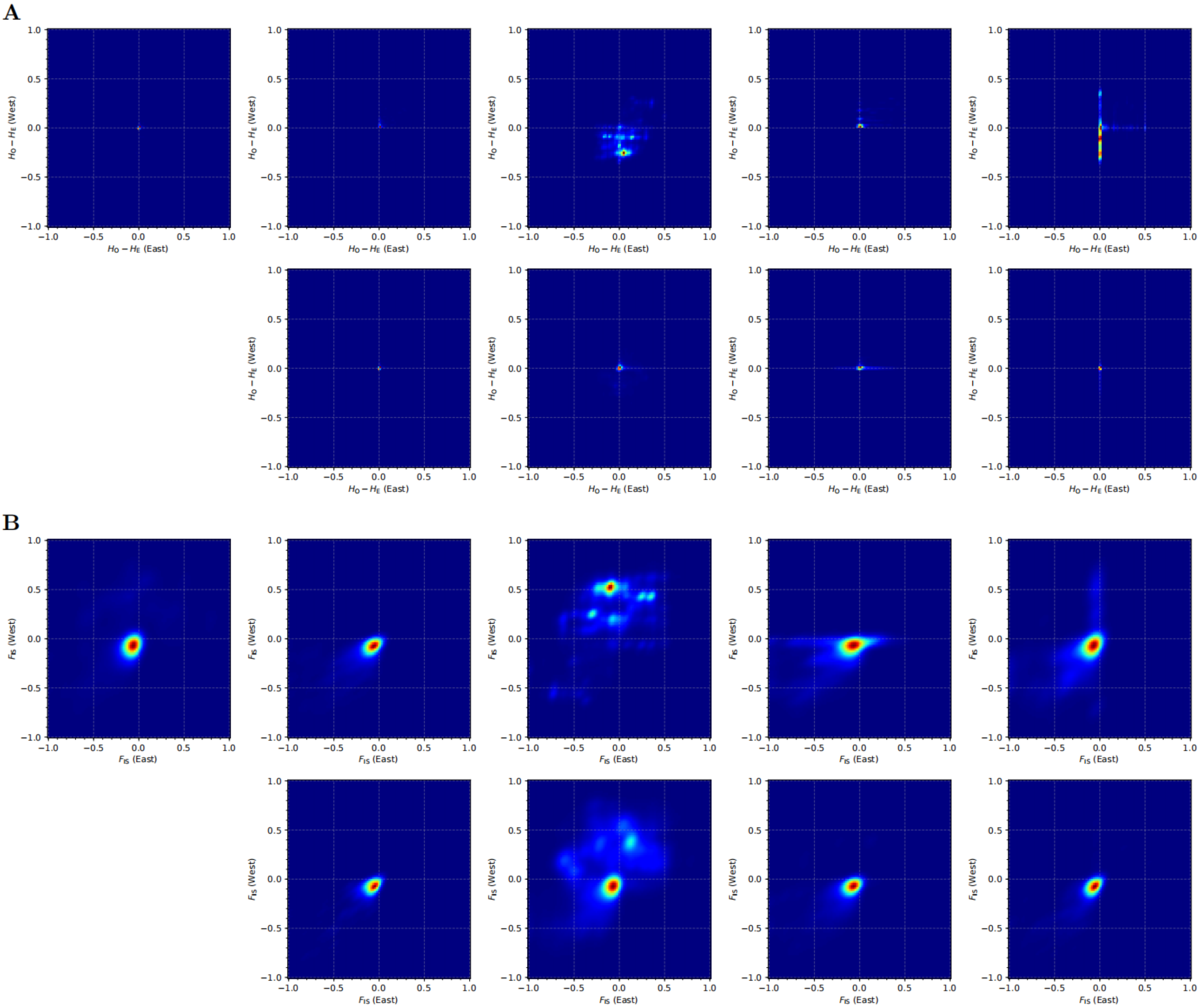


**Supplementary Figure 32. Inbreeding coefficient spectrum between East and West populations for the nine clusters.** **A)** Difference between observed heterozygosity ($H_\mathrm{O}$) and expected heterozygosity ($H_\mathrm{E}$). **B)** Inbreeding coefficient $F_\mathrm{IS} = (H_\mathrm{E} - H_\mathrm{O})/H_\mathrm{E}$. For both panels, clusters 1–5 are arranged from left to right in the top row, and clusters 6–9 in the bottom row.

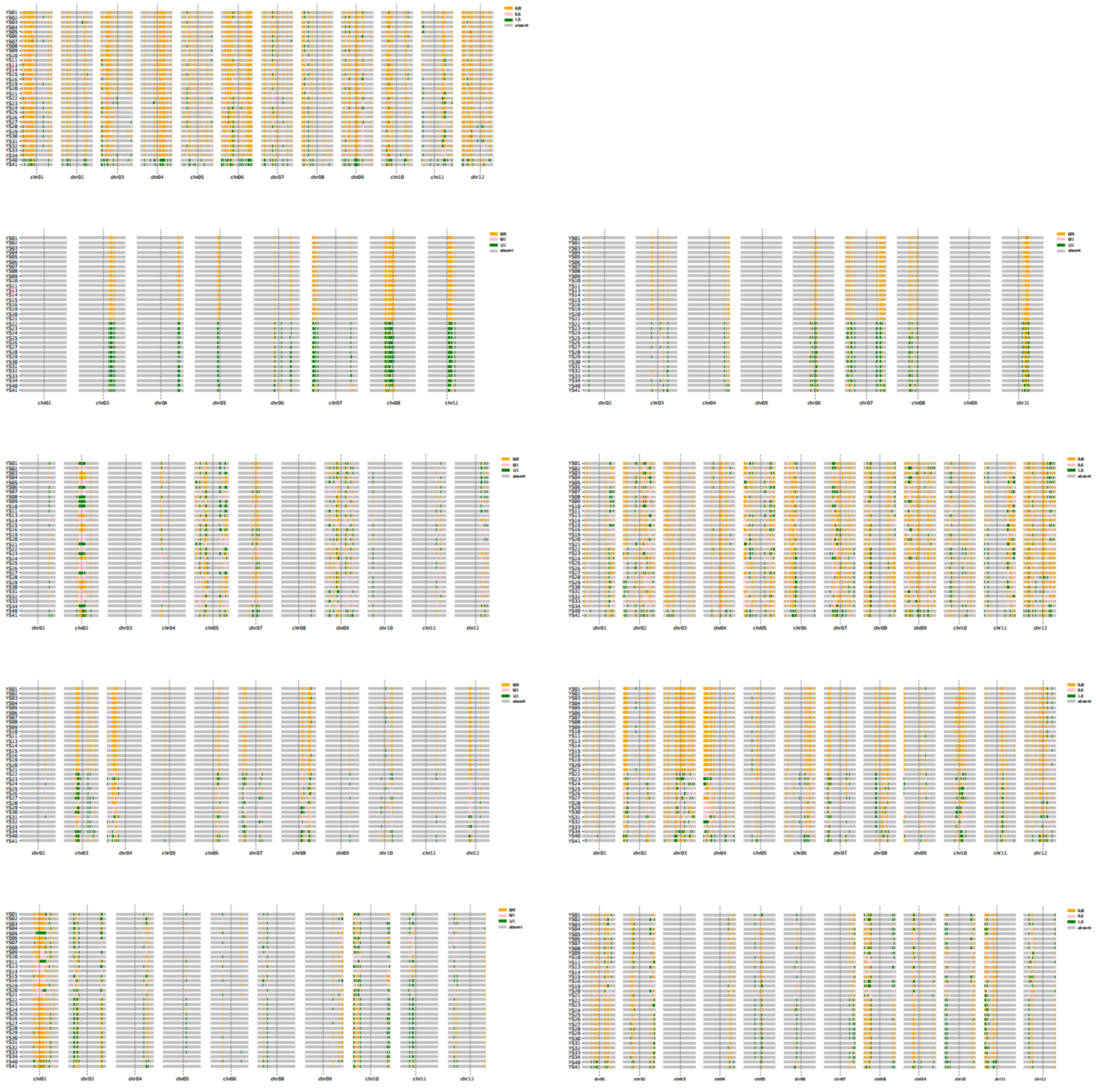

**Supplementary Figure 33. Genotypes across chromosomes of the nine clusters.** Left panels (from top to bottom): clusters 1–5; Right panels: clusters 6–9.

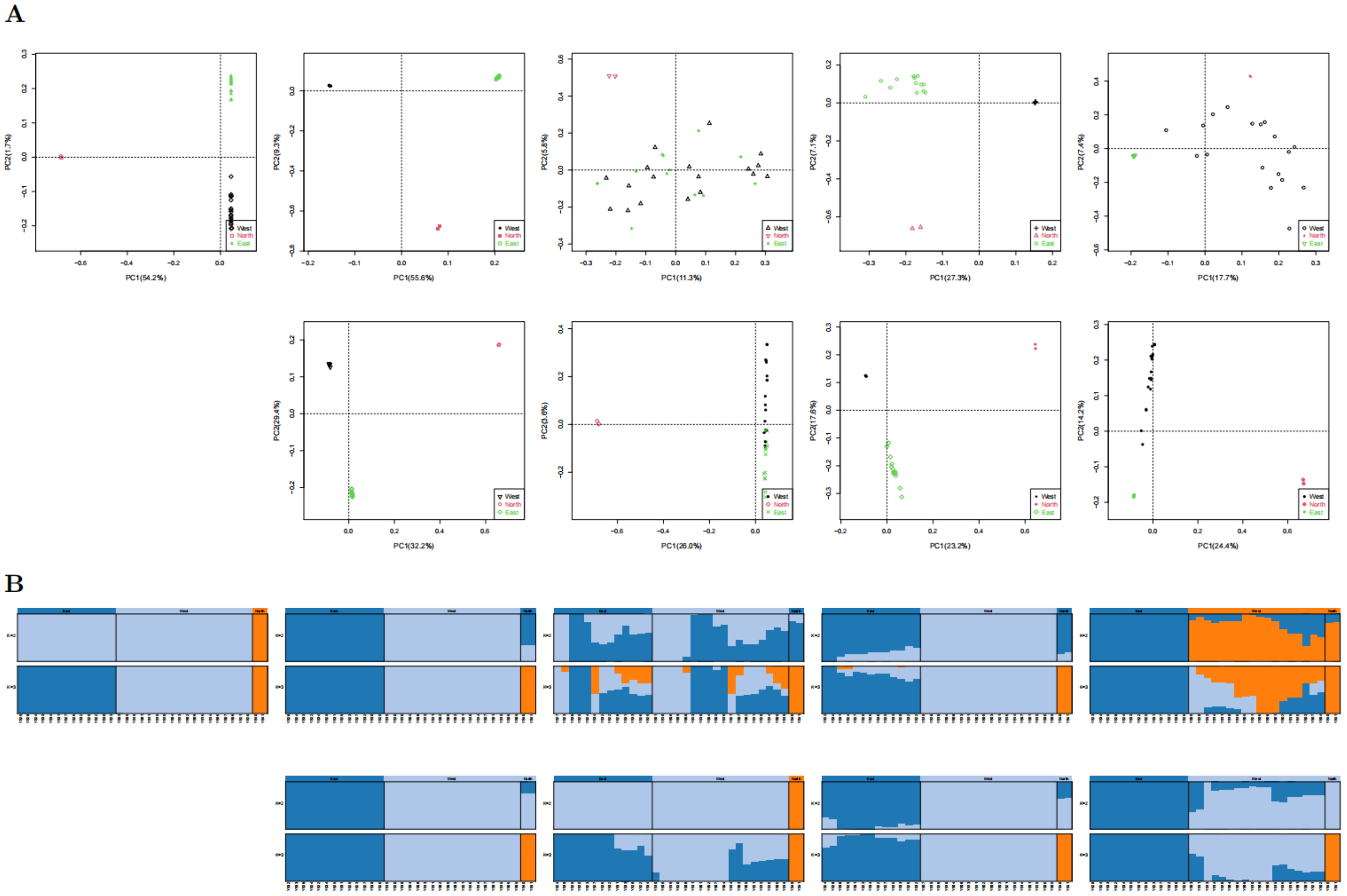


**Supplementary Figure 34. Principal Component Analysis (PCA) and genetic admixture analysis for the nine clusters.** **A)** PCA plot showing genetic clustering based on the first two principal components. **B)** Genetic ancestry proportions inferred by ADMIXTURE. For both panels, clusters 1–5 are arranged from left to right in the top row, and clusters 6–9 in the bottom row.

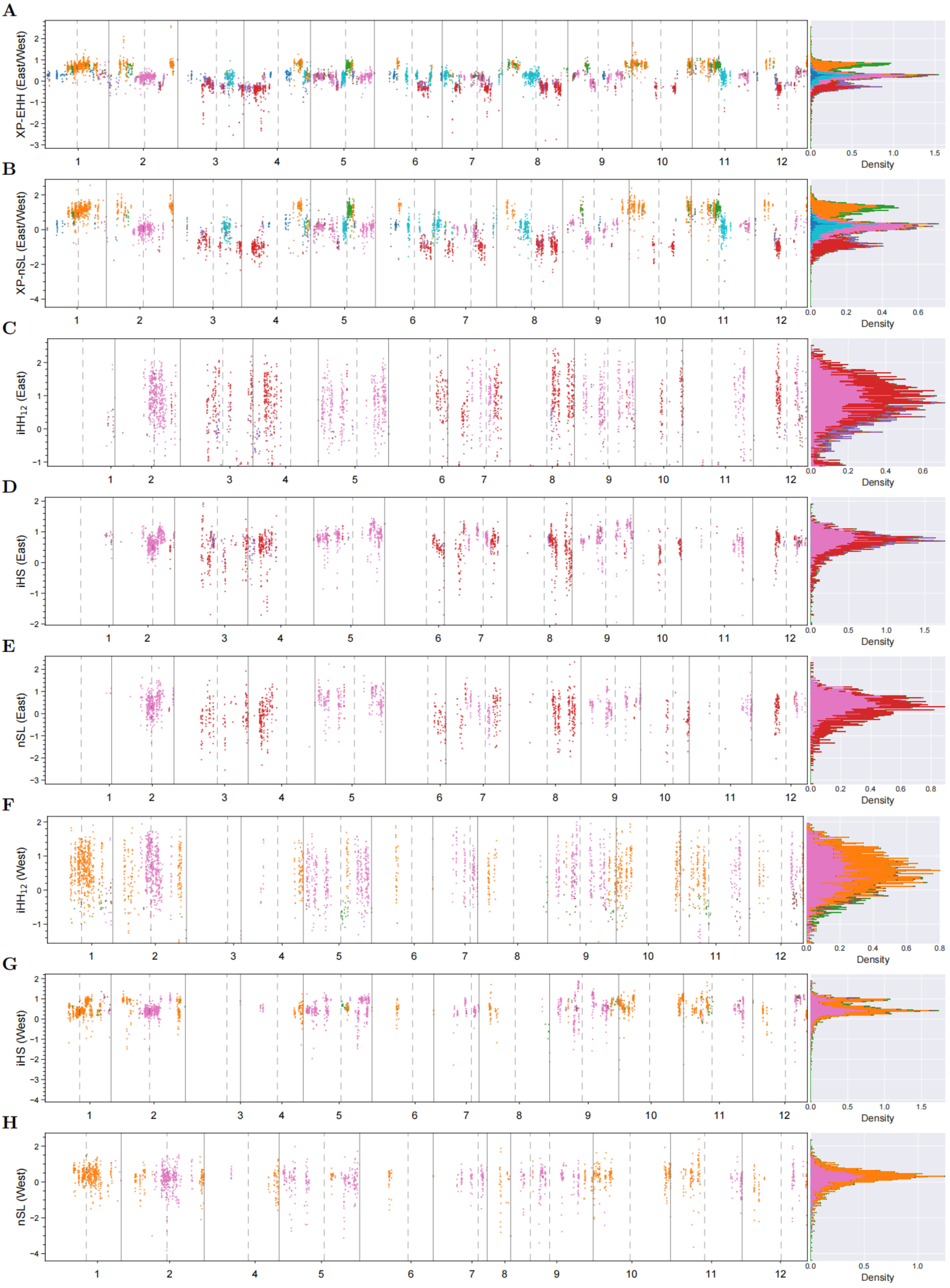


**Supplementary Figure 35. Manhattan plots of positive selection signals by selscan. A–B)** Cross-population signals including XP-EHH (A) and XP-nSL (B) between East and West populations, with the West population as the reference. Highly positive values represent potential positive selection in the East population, while highly negative values represent potential selection in the West population. **C–E)** $iHH_{12}$, iHS, and nSL for the East population, respectively. **F–H)** $iHH_{12}$, iHS, and nSL for the West population, respectively. Higher positive values represent potential positive selection on derived alleles.

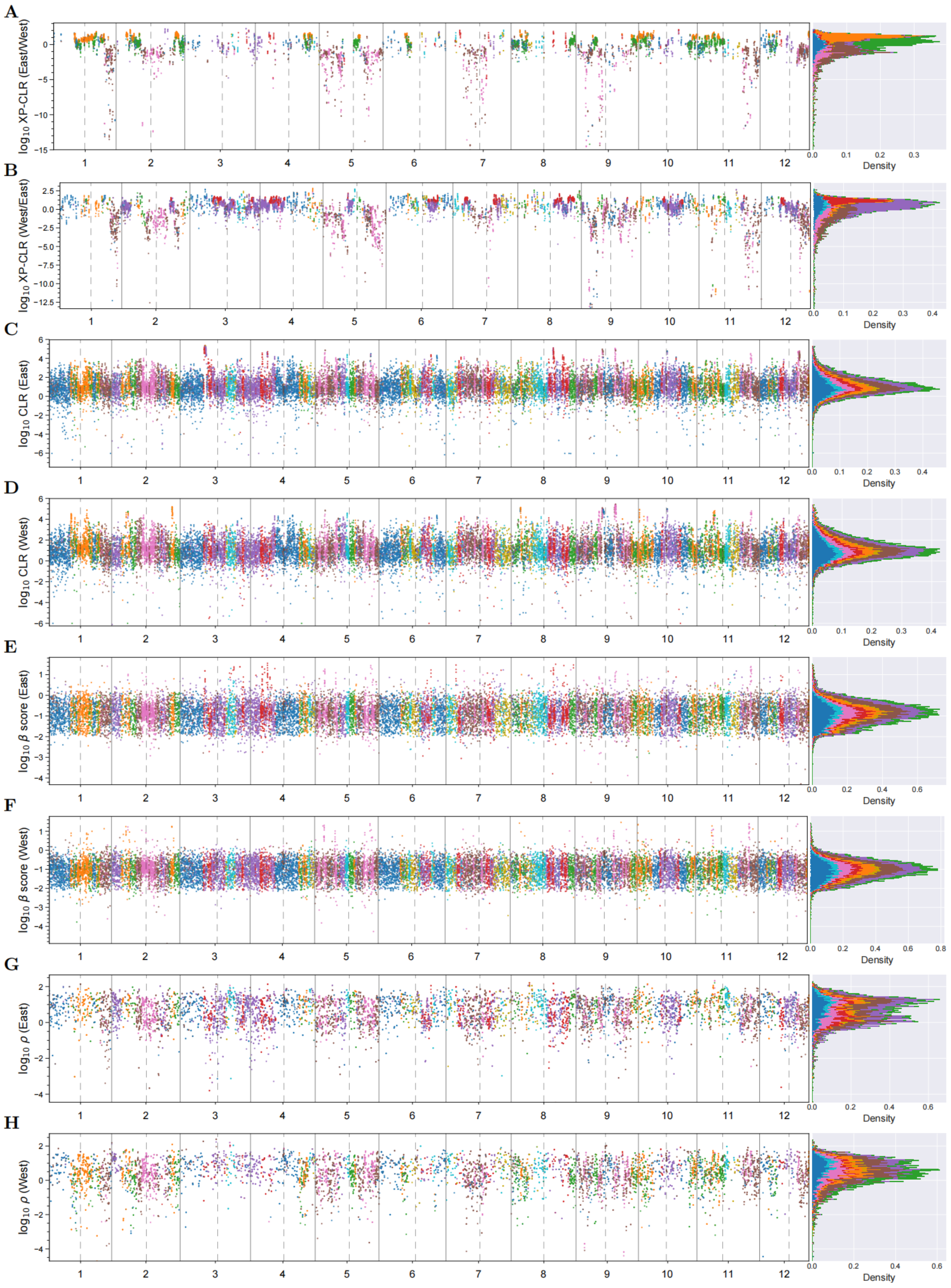


**Supplementary Figure 36. Manhattan plots of positive selection, balancing selection, and population recombination rates. A–B)** XP-CLR signals using West and East populations as reference, respectively; higher values represent potential positive selection in the East and West populations, respectively. **C–D)** CLR values for East and West populations estimated by SweepFinder; higher values represent potential positive selection. **E–F)** $\beta$ scores from betascan; higher values represent signals of potential balancing selection. **G–H)** Population recombination rates ($\rho$) for East and West populations estimated by FastEPRR.

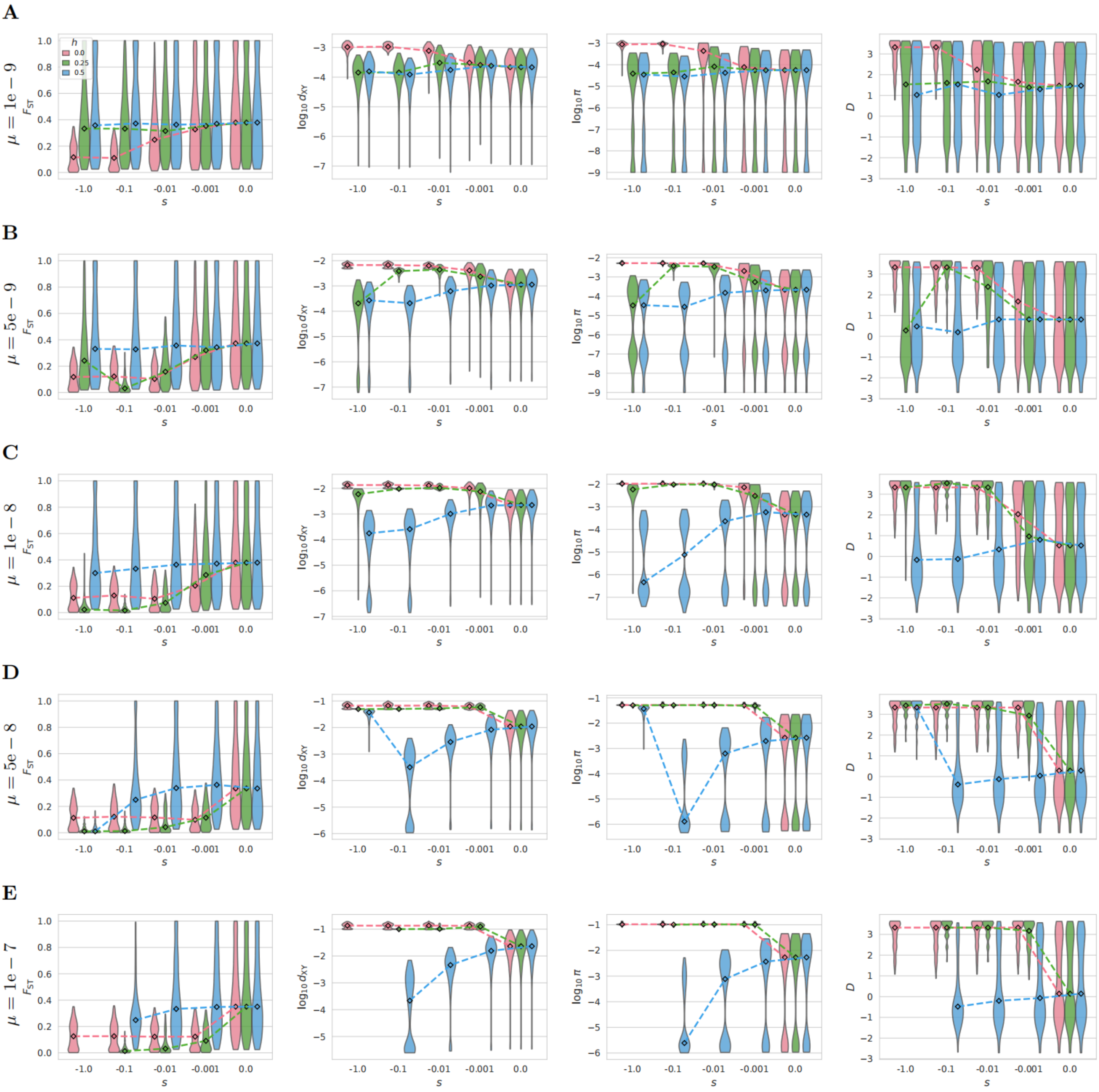


**Supplementary Figure 37. Forward simulations with varied mutation rate ($\mu \in \{1e-9, 5e-9, 1e-8, 5e-8, 1e-7\}$).**

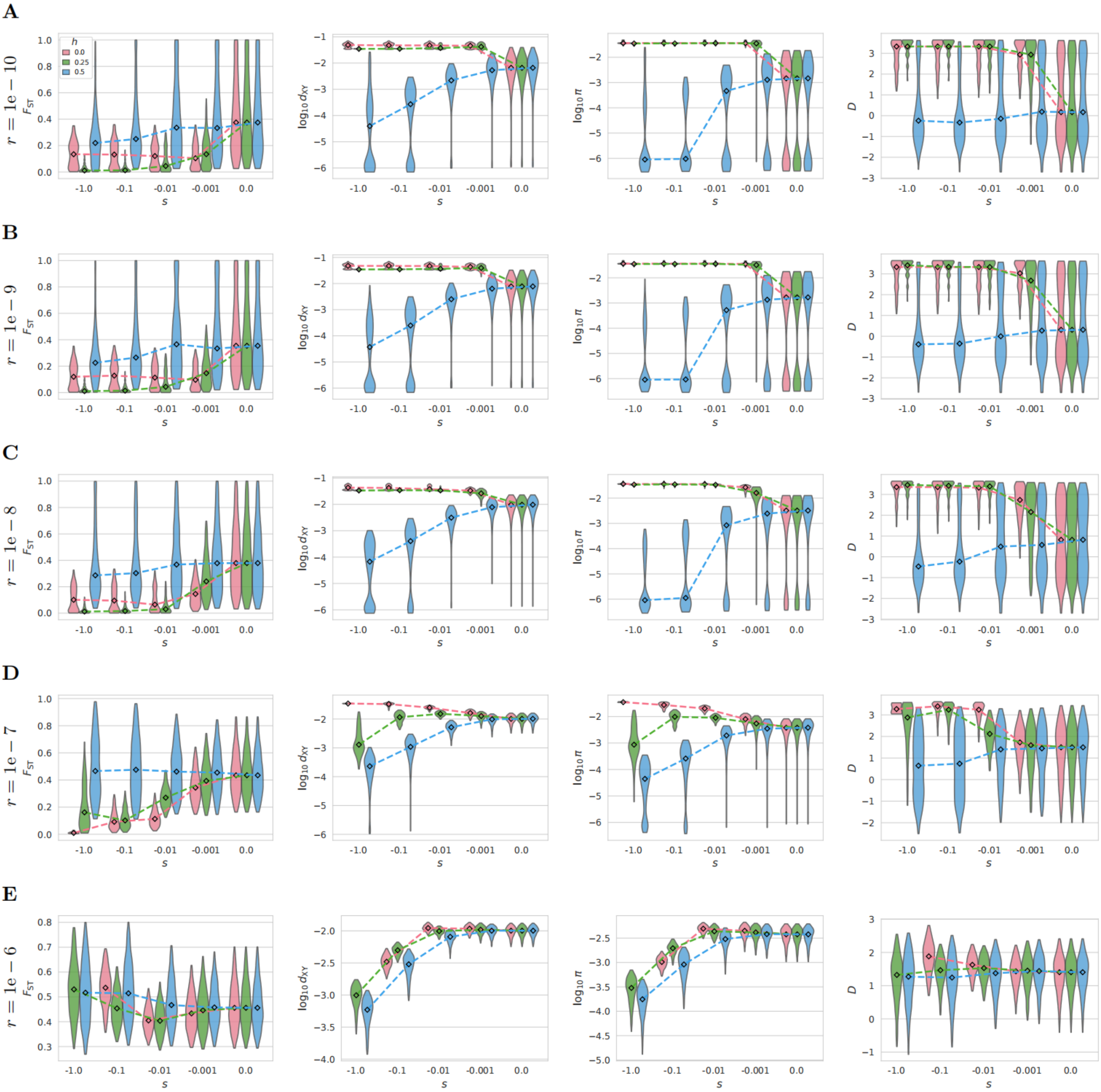


**Supplementary Figure 38. Forward simulations with varied recombination rate ($r \in \{0.01, 0.1, 1, 10, 100\}$ cM/Mb).**

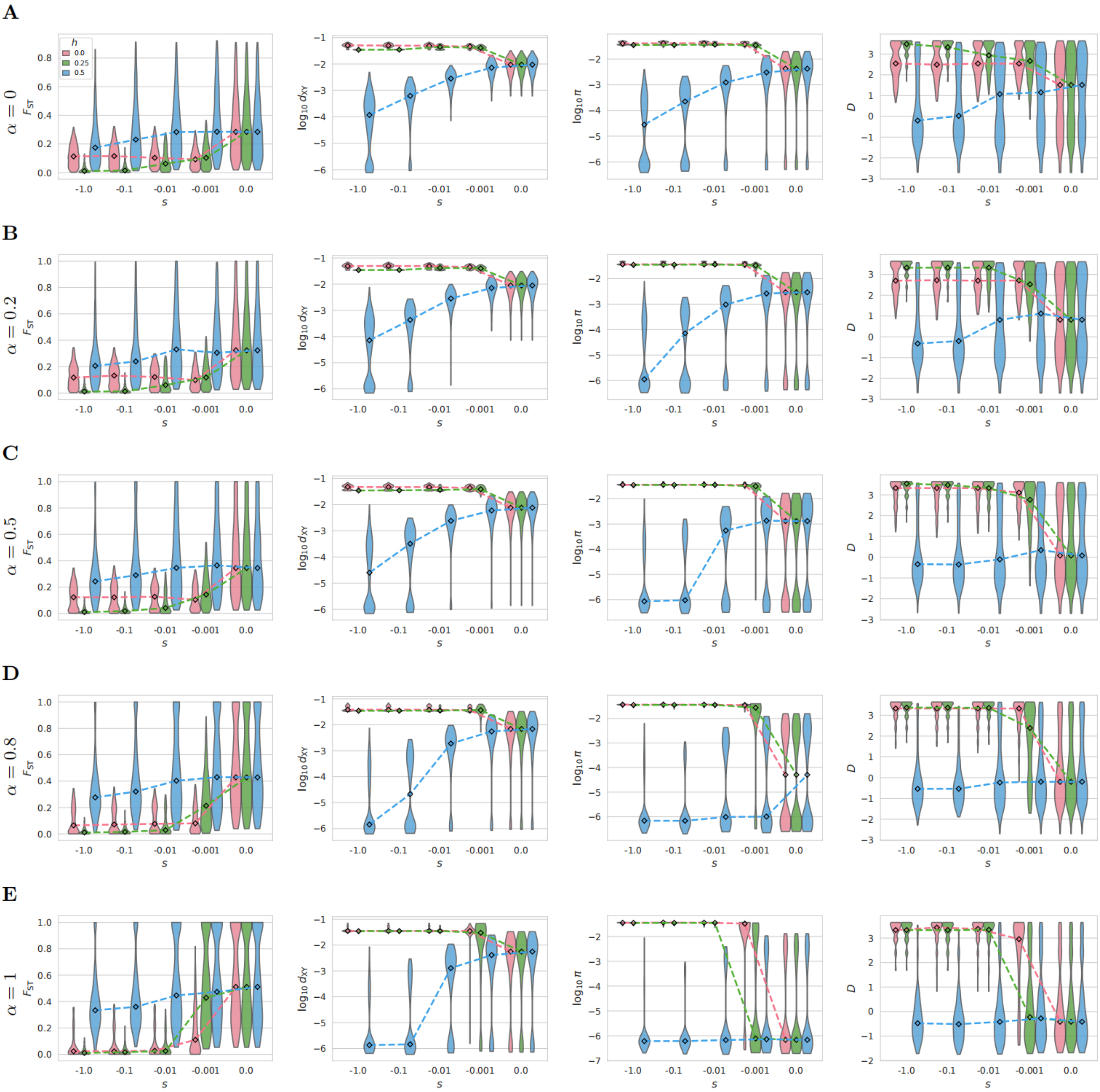

**Supplementary Figure 39. Forward simulations with varied selfing rate ($\alpha \in \{0, 0.2, 0.5, 0.8, 1\}$).**

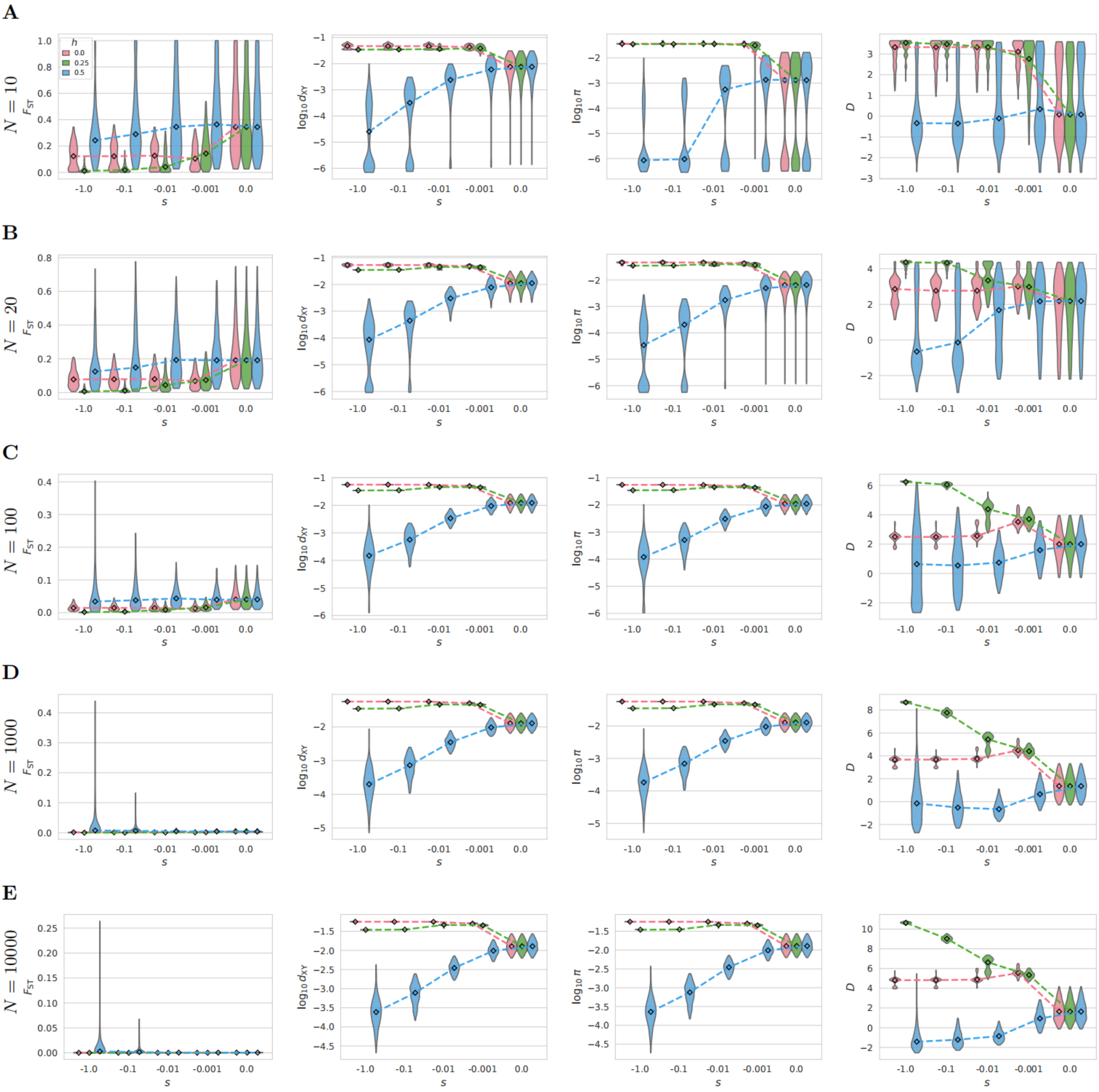


**Supplementary Figure 40. Forward simulations with varied population size ($N \in \{10, 20, 100, 1000, 10000\}$).**

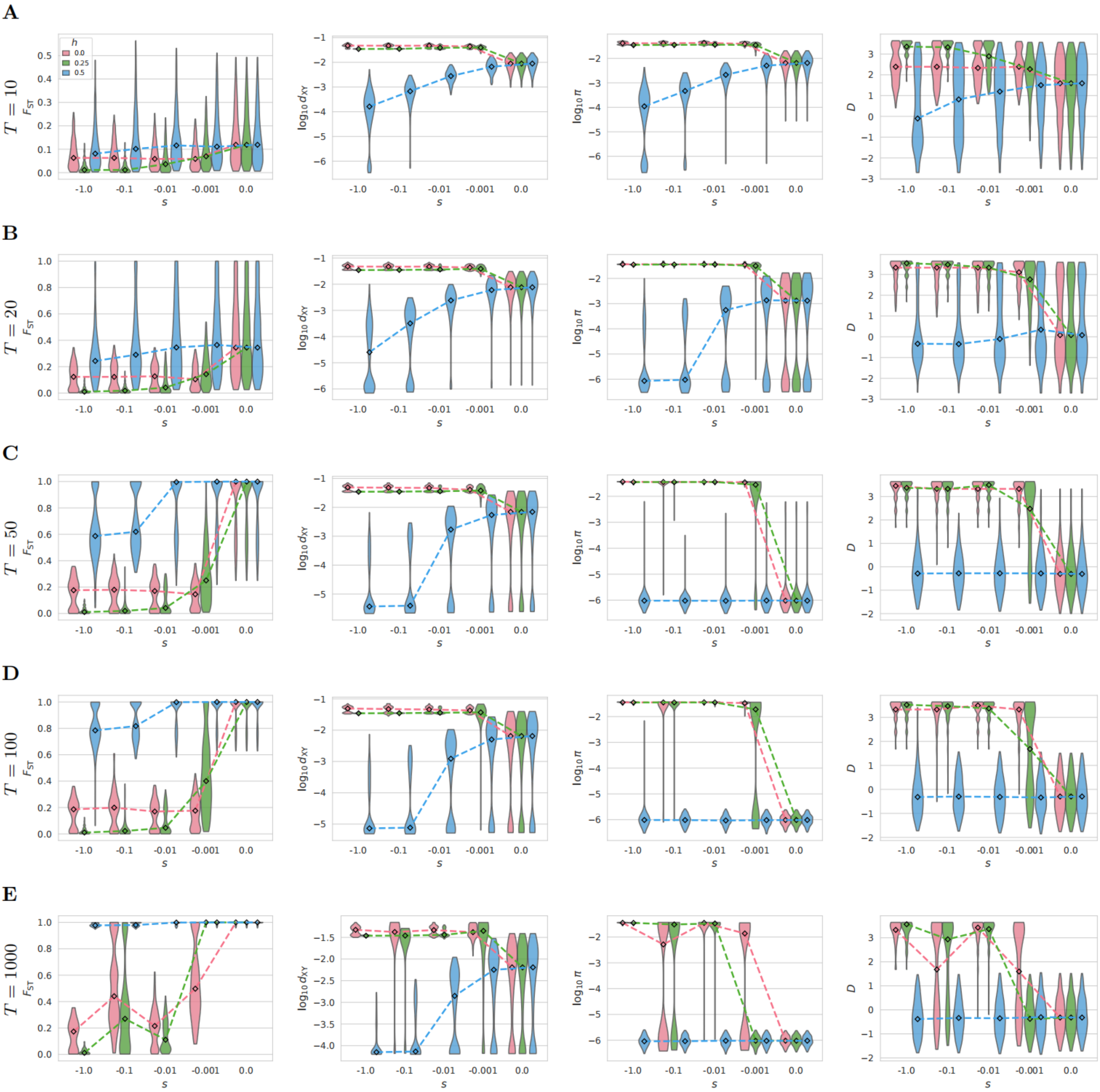


**Supplementary Figure 41. Forward simulations with varied bottleneck duration ($T \in \{10, 20, 50, 100, 1000\}$).**

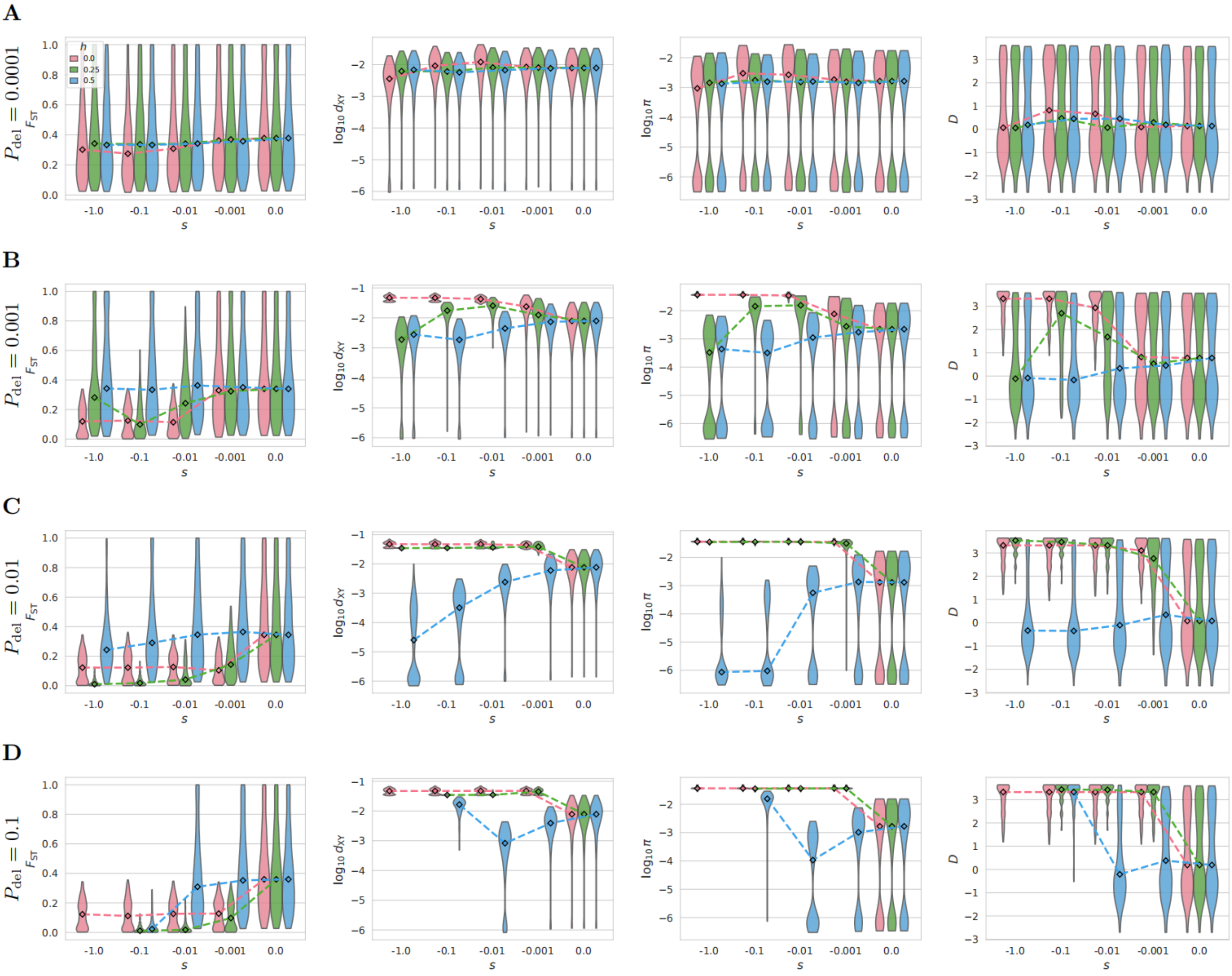


**Supplementary Figure 42.** Forward simulations with varied proportion of deleterious mutations ($P_{\mathrm{del}} \in \{0.01\%, 0.1\%, 1\%, 10\%\}$).

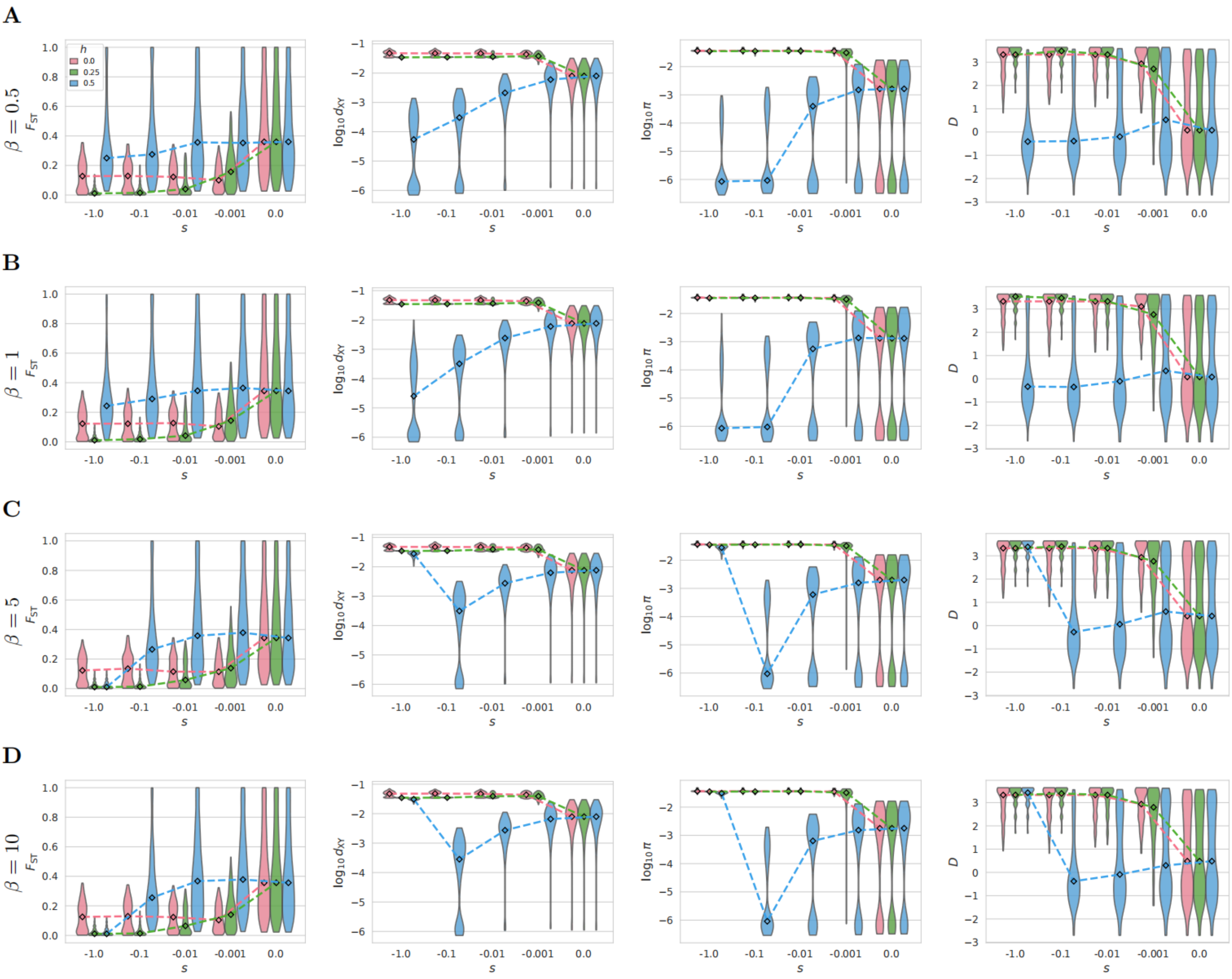


**Supplementary Figure 43. Forward simulations with varied shape of gamma distribution ($\beta \in \{0.5, 1, 5, 10\}$).**

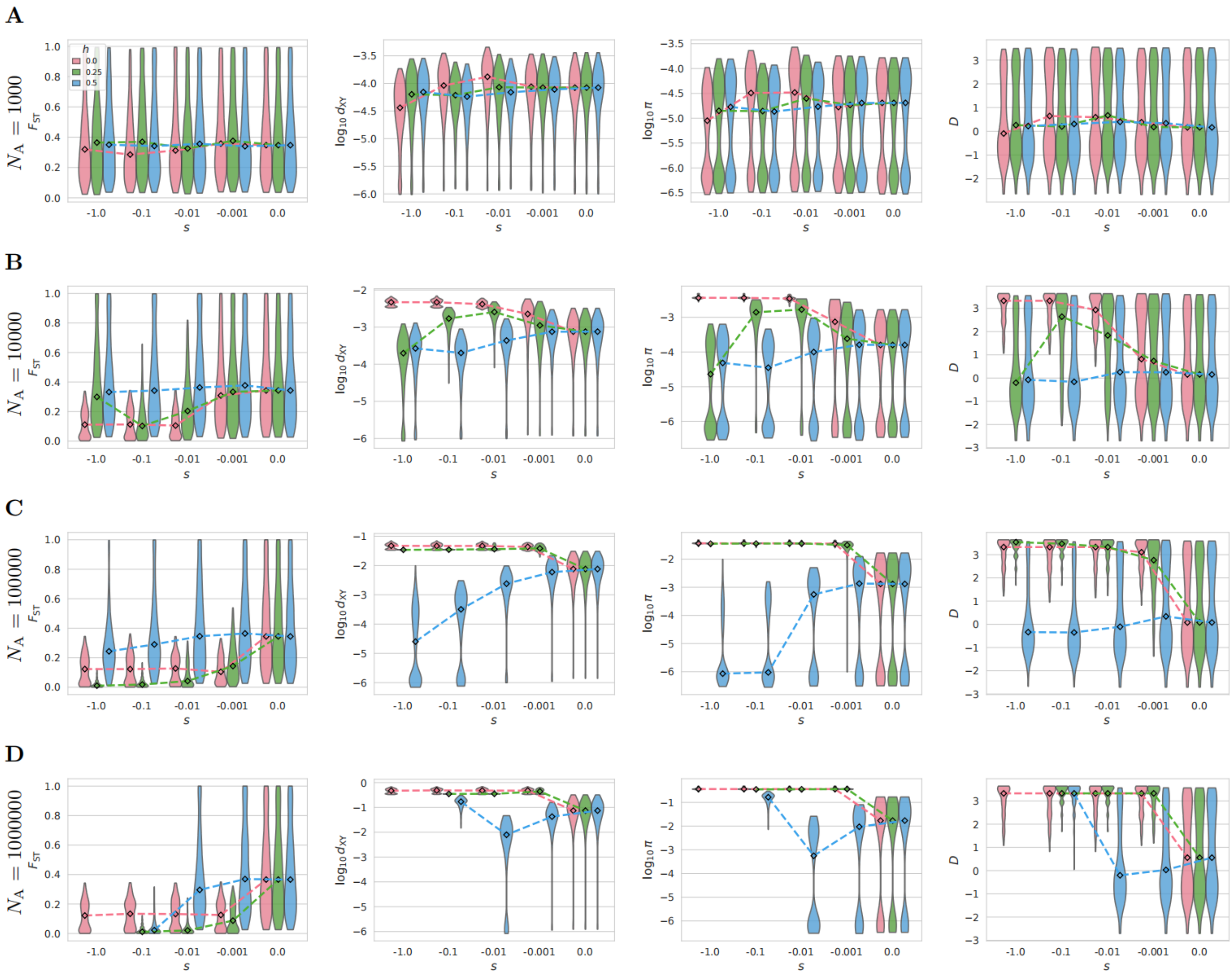


**Supplementary Figure 44. Forward simulations with varied ancestral population size ($N_A \in \{10^3, 10^4, 10^5, 10^6\}$).**

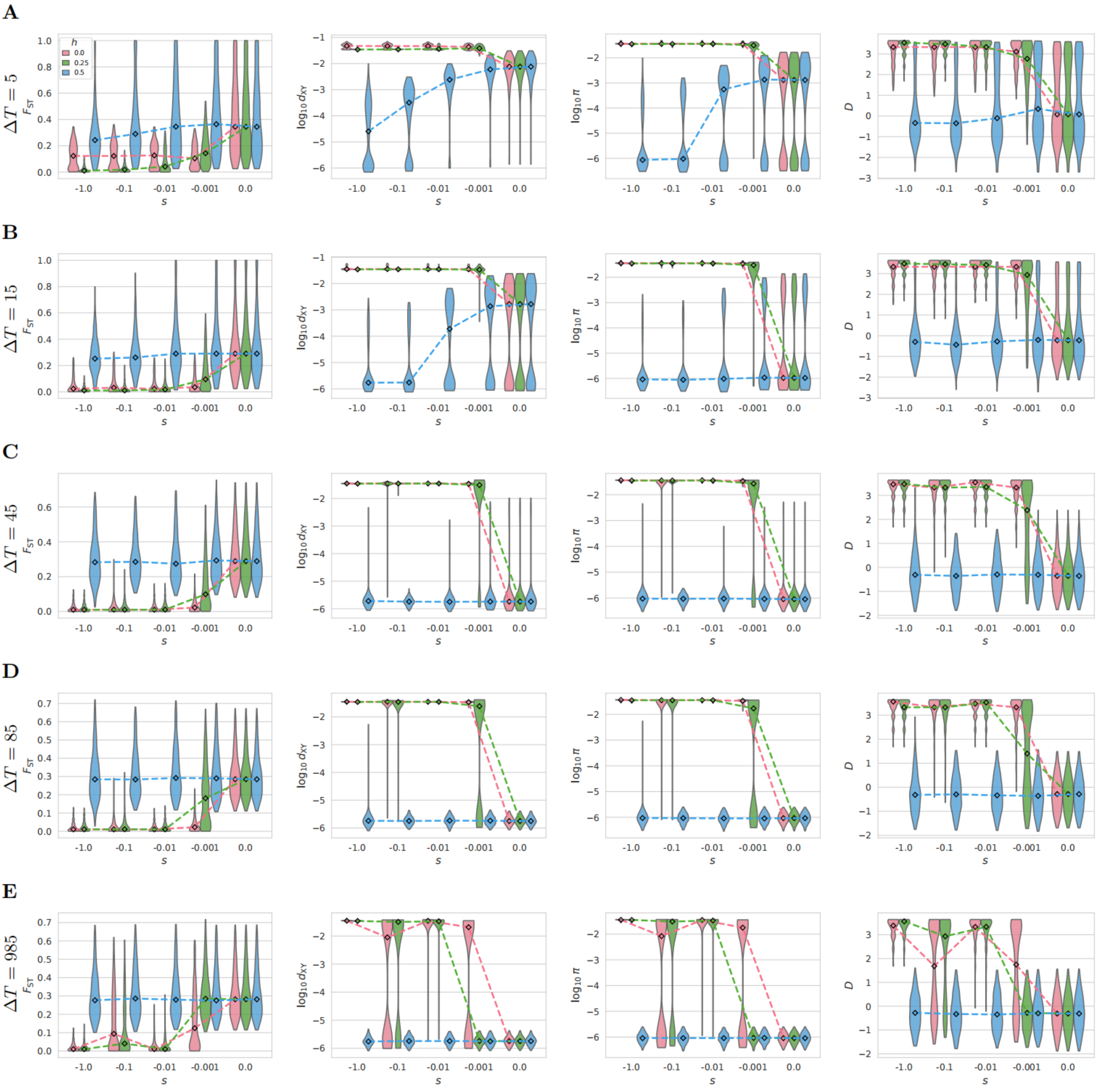


**Supplementary Figure 45.** **Forward simulations with varied lag time between bottleneck and split ($\Delta T \in \{5, 15, 45, 85, 985\}$).**

**Supplementary Table 8**. Estimates of genome-wide genetic diversity in varied plant species.

| Species | Group | Life form | IUCN categories* | Threatened ? | $\pi$ | Reference |
|---|---|---|---|---|---|---|
| *Pinus squamata* | gymnosperm | tree | CR | Yes | 3.32e-5 | This study |
| *Pinus bungeana* | gymnosperm | tree | LC | No | 1.68e-3 | This study |
| *Pinus gerardiana* | gymnosperm | tree | NT | Yes | 3.05e-3 | This study |
| *Cupressus gigantea* | gymnosperm | tree | VU | Yes | 2.01e-3 | 32 |
| *Cupressus duclouxiana* | gymnosperm | tree | DD | No | 3.08e-3 | 32 |
| *Ginkgo biloba* | gymnosperm | tree | EN | Yes | 2.57e-3 | 33 |
| *Alsophila latebrosa* | fern | tree | CR | Yes | 1.37e-2 | 34 |
| *Alsophila costularis* | fern | tree | NE | No | 9.8e-3 | 34 |
| *Alsophila spinulosa* | fern | tree | NE | No | 1.8e-3 | 34 |
| *Acer yangbiense* | angiosperm | tree | EN | Yes | 3.13e-3 | 35 |
| *Rhododendron griersonianum* | angiosperm | tree | NE | No | 1.94e-3 | 36 |
| *Rhododendron delavayi* | angiosperm | tree | LC | No | 1.30e-2 | 36 |
| *Buddleja alternifolia* | angiosperm | tree | LC | No | 9.13e-3 | 37 |
| *Malania oleifera* | angiosperm | tree | VU | Yes | 3.87e-3 | 38 |
| *Magnolia sinica* | angiosperm | tree | CR | Yes | 1.49e-2 | 39 |
| *Tetracentron sinense* | angiosperm | tree | DD | Yes | 1.1e-2 | 40 |
| *Ostrya rehderiana* | angiosperm | tree | CR | Yes | 1.66e-3 | 30 |
| *Ostrya chinensis* | angiosperm | tree | LC | No | 2.79e-3 | 30 |
| *Cercidiphyllum japonicum* | angiosperm | tree | LC | No | 1.1e-3 | 41 |
| *Dipteronia dyeriana* | angiosperm | tree | EN | Yes | 5.15e-4 | 29 |
| *Dipteronia sinensis* | angiosperm | tree | DD | No | 2.98e-3 | 29 |
| *Liriodendron chinense* | angiosperm | tree | NT | Yes | 6.89e-4 | 42 |
| *Liriodendron tulipifera* | angiosperm | tree | LC | No | 5.6e-5 | 42 |
| *Populus ilicifolia* | angiosperm | tree | VU | Yes | 8.0e-4 | 43 |
| *Davidia involucrata* | angiosperm | tree | NE | Yes | 5.85e-3 | 44 |
| *Dasiphora fruticosa* | angiosperm | tree | NE | No | 3.1e-3 | 45 |
| *Quercus rex* | angiosperm | tree | LC | No | 7.10e-3 | 46 |
| *Quercus sichourensis* | angiosperm | tree | CR | Yes | 4.39e-3 | 46 |
| *Hopea hainanensis* | angiosperm | tree | EN | Yes | 3.31e-3 | 47 |
| *Hopea reticulata* | angiosperm | tree | EN | Yes | 3.97e-3 | 47 |
| *Vitis vinifera* | angiosperm | tree | LC | No | 5.49e-3 | 48 |
| *Sapindus delavayi* | angiosperm | tree | LC | No | 6.3e-2 | 49 |
| *Sapindus rarak* | angiosperm | tree | LC | No | 7.1e-2 | 49 |
| *Bretschneidera sinensis* | angiosperm | tree | EN | Yes | 6.36e-2 | 50 |
| *Artocarpus nanchuanensis* | angiosperm | tree | CR | Yes | 1.33e-3 | 51 |
| *Thlaspi arvense* | angiosperm | herb | NE | No | 5.7e-4 | 52 |
| *Citrullus colocynthis* | angiosperm | herb | NE | No | 6.75e-3 | 53 |
| *Citrullus mucosospermus* | angiosperm | herb | NE | No | 7.92e-4 | 53 |
| *Boehmeria nivea* | angiosperm | herb | NE | No | 3.85e-3 | 54 |
| *Ricinus communis* | angiosperm | herb | LC | No | 1.95e-3 | 55 |

| | | | | | | |
|---|---|---|---|---|---|---|
| *Prunus persica* | angiosperm | tree | LC | No | 1.5e-3 | 56 |
| *Phoenix dactylifera* | angiosperm | tree | LC | No | 9.2e-3 | 57 |
| *Gossypium raimondii* | angiosperm | tree | EN | Yes | 1.2e-4 | 58 |
| *Setaria italica* | angiosperm | herb | NE | No | 2.33e-3 | 58 |
| *Citrullus lanatus* | angiosperm | herb | NE | No | 2.41e-3 | 58 |
| *Sorghum bicolor* | angiosperm | herb | LC | No | 2.49e-3 | 58 |
| *Brachypodium distachyon* | angiosperm | herb | NE | No | 2.67e-3 | 58 |
| *Glycine soja* | angiosperm | herb | NE | No | 2.74e-3 | 58 |
| *Arabidopsis thaliana* | angiosperm | herb | NE | No | 3.15e-3 | 58 |
| *Populus trichocarpa* | angiosperm | tree | LC | No | 3.17e-3 | 58 |
| *Capsella rubella* | angiosperm | herb | NE | No | 3.27e-3 | 58 |
| *Medicago truncatula* | angiosperm | herb | LC | No | 5.14e-3 | 58 |
| *Oryza rufipogon* | angiosperm | herb | LC | No | 6.36e-3 | 58 |
| *Prunus davidiana* | angiosperm | tree | LC | No | 9.88e-3 | 58 |
| *Cucumis sativus* | angiosperm | herb | NE | No | 1.32e-2 | 58 |
| *Zea mays* | angiosperm | herb | LC | No | 1.39e-2 | 58 |
| *Citrus reticulata* | angiosperm | tree | NE | No | 1.50e-2 | 58 |
| *Populus tremula* | angiosperm | tree | LC | No | 1.47e-2 | 59 |
| *Populus tremuloides* | angiosperm | tree | LC | No | 1.6e-2 | 59 |
| *Ziziphus jujuba* | angiosperm | tree | LC | No | 2.19e-3 | 60 |
| *Betula pendula* | angiosperm | tree | LC | No | 8.8e-3 | 61 |
| *Malus domestica* | angiosperm | tree | NE | No | 2.2e-3 | 62 |

* CR, Critically Endangered; EN, Endangered; VU, Vulnerable; NT, Near Threatened; LC, Least Concern; DD, Data Deficient; NE, Not Evaluated.

Note: This table is based on previous compilations[30,36] with additional records added in this study.

### Supplementary Methods

#### Supplementary Method 1. Inference of contemporary $N_e$ and historical demographic dynamics

To confirm the GONE analysis, we performed demographic inference using two updated linkage disequilibrium (LD)-based methods: currentNe2 and GONE2[63]. From the SNP dataset without Minor Allele Count filtering, we extracted SNPs for each population and applied rigorous quality control using VCFtools (parameters: --max-missing 0.8 --mac 1). For each population, we randomly subsampled 1 million SNPs as input for both currentNe2 and GONE2. To ensure consistency, both methods were executed with identical parameters: a recombination rate of -r 0.12 and a maximum sample size of -s 100000. To account for potential population structure, we also enabled the -x option. The entire pipeline—from SNP extraction to the final analysis—was repeated 200 times to generate confidence intervals and mitigate stochastic sampling bias.

For long-term demographic inference, we employed coalescent-based methods, Stairway Plot v2.1[64] and the Pairwise Sequentially Markovian Coalescent (PSMC)[65], both of which were performed by excluding gene regions to approximate site neutrality. These analyses adopted a mutation rate of 3.5e-8 per site per generation[6]. For the Stairway Plot, the folded SFS was estimated from the VCF dataset using easySFS[66]. For the PSMC analysis, we selected the four individuals with the highest sequencing depth per population to ensure robust consensus calling. Consensus sequences were generated using BCFtools, with sites filtered for a read depth of 5x to 50x using vcfutils.pl. The PSMC model was executed with parameters -N30 -t15 -r5 and an atomic time interval pattern of -p "4+25*2+4+6".

#### Supplementary Method 2. Demographic Modeling and Divergence Inference

To infer the divergence history and subsequent population size fluctuations between the East and West populations, we employed two coalescent-based approaches: fastsimcoal2 v2.6[67] and DILS[68]. For all analyses, a mutation rate of 3.5e-8 per site per generation was applied.

For the fastsimcoal2 analysis, we constructed an un-folded two-dimensional joint site frequency spectrum (2D-SFS) using realSFS in ANGSD[69], restricting the data to 14,651,899,749 intergenic sites to mitigate the effects of selection. We performed 100,000 coalescent simulations to estimate the expected 2D-SFS and log-likelihood. Global maximum likelihood estimates were obtained from 50 independent runs, and the optimal demographic model was identified using the Akaike Information Criterion (AIC).

For the DILS analysis, to optimize computational efficiency, we utilized 4-fold degenerate sites (4D sites) partitioned into 4,628 genomic windows of 5 Mb each. For the two-population analysis, DILS automatically evaluated four competing models: Strict Isolation (SI), Ancient Migration (AM), Isolation with Migration (IM), and Secondary Contact (SC). For single-population inferences, the software compared constant size, expansion, and contraction models. Based on the goodness-of-fit assessments, the variable population growth models demonstrated superior performance and were selected for final parameter estimation.

#### Supplementary Method 3. Reconstruction of ancestral sequences

To reconstruct ancestral states, we extracted sequence data directly from the VCF files (dataset 1), selecting the individual with the highest sequencing depth as the representative for each species: BPS02 for *P. bungeana*, TBPS03 for *P. gerardiana*, and YS22 for *P. squamata*. Using *P. albicaulis* as the outgroup and following the previously established species tree topology, ancestral sequence reconstruction was performed for each chromosome using the empirical Bayesian method (-asr) in IQ-TREE[70]. To maintain high stringency, ancestral states were assigned only to sites with a posterior probability greater than 0.99; sites falling below this threshold were marked as ambiguous ("N"). The reconstructed sequence at the crown node of the three ingroup pines was defined as the ancestral state for all downstream analyses. Ultimately, this process recovered 1,628,411,634 high-confidence ancestral sites, representing 55.82% of the total genomic positions.
